# Capturing non-local through-bond effects in molecular mechanics force fields: II. Using fractional bond orders to fit torsion parameters

**DOI:** 10.1101/2022.01.17.476653

**Authors:** Chaya D. Stern, Jessica Maat, David L. Dotson, Christopher I. Bayly, Daniel G. A. Smith, David L. Mobley, John D. Chodera

## Abstract

Accurate small molecule force fields are crucial for predicting thermodynamic and kinetic properties of drug-like molecules in biomolecular systems. Torsion parameters, in particular, are essential for determining conformational distribution of molecules. However, they are usually fit to computationally expensive quantum chemical torsion scans and generalize poorly to different chemical environments. Torsion parameters should ideally capture local through-space non-bonded interactions such as 1-4 steric and electrostatics and non-local through-bond effects such as conjugation and hyperconjugation. Non-local through-bond effects are sensitive to remote substituents and are a contributing factor to torsion parameters poor transferability. Here we show that fractional bond orders such as the Wiberg Bond Order (WBO) are sensitive to remote substituents and correctly captures extent of conjugation and hyperconjugation. We show that the relationship between WBO and torsion barrier heights are linear and can therefore serve as a surrogate to QC torsion barriers, and to interpolate torsion force constants. Using this approach we can reduce the number of computationally expensive QC torsion scans needed while maintaining accurate torsion parameters. We demonstrate this approach to a set of substituted benzene rings.

## 1 Introduction

Molecular mechanics (MM) methods rely on empirical force fields inspired by Newtonian physics to describe the potential energy of the system, and are widely used to study larger systems with 10^3^–10^6^ atoms [29]. They are sufficiently computationally efficient and accurate to study biologically relevant systems, provide atomistic details of mechanisms involving biomolecular conformational dynamics in solution, and reliably predict thermodynamic properties such as binding free energies [10, 49, 56]. However, given the larger chemical space that small molecule force fields must cover to adequately represent drug-like molecules and common metabolites, their development—in terms of achieving desired accuracy over the chemical space of interest—has lagged behind protein force fields [27, 81].

### 1.1 The torsional functional describes the potential energy of internal rotation

In MM force fields, the potential energy is constructed with terms for bond stretching, angle bending, internal rotations, electrostatics and Lennard-Jones for attractive and repulsive forces [8, 11, 36]. The free parameters in these functionals are generally fit to reproduce experimental and quantum chemical (QC) data. In class I molecular mechanics force fields [14, 42] (e.g. CHARMM [8], AMBER [11], and OPLS [36]), the torsional potential is given by a truncated Fourier cosine series expansion:

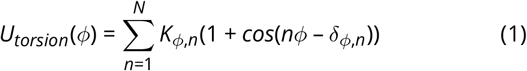

where *ϕ* denotes a single torsion, and the sum is over multiplicities *n*. The parameters *K_ϕ,n_* determine the barrier heights, the multiplicities *n* determine the number of minima in that term, and the phase angles *δ_ϕ,n_* determine the phase offset for each term. In most popular molecular mechanics force fields, the *N* can go up to 6, and the phase angles *δ* are usually set to 0° or 180° to ensure the potential is symmetric about *ϕ* = 0° [23].

The true torsional energy about a bond is determined by a combination of local and non-local effects from conjugation, hypercojugation, sterics, and electrostatics [20, 39, 58, 76]. In most force fields, non-local through-space steric and elecrostatic interactions, or non-bonded interactions beyond 1-4 atoms, are accounted for by non-bonded terms. For the 1-4 atoms, most forcefields include a scaled 1-4 electrostatics and LJ to account for 6-12 LJ and electrostatics at such close range [11]. The torsional energy profile should only capture close range 1-4 non-bonded interaction and conjugation or hyperconjugation effects that perturb these contributions. The torsion parameters should not correct for long-range through-space interactions.

This torsion functional, which models internal rotation, is particularly challenging to parameterize for small molecules, and is generally the least transferable relative to other valence terms for several reasons [32, 73, 76]. First, torsion parameters are usually fit to computationally expensive QC torsion scans, which introduces a bottleneck to setting up simulations if this data must be generated for each new molecule of interest. Second, torsions are ‘soft’, flexible degrees of freedom, compared to bond and angles. Relatively small variations to the torsional potential surface can strongly influence molecular conformation distributions [32]. Given how critical torsion parameters are in determining conformation distributions in simulations, it is prudent for them to be accurately parameterized. Third, torsional potentials can be strongly influenced by distal substituents due to changes in conjugation or hypercojugation, a nonlocal effect very difficult to represent in a force field that uses only the local chemical environment to define parameters [67].

When parameterizing a molecular system, a typical class I MM force field assigns atom types—which capture an atom’s atomic number and chemical environment—to atoms in the system [35, 72]. The aim of these atom types is to enable transferability of parameters to other atoms in similar chemical environments. A *torsion type* is defined by the quartet of bonded atom types of the four atoms involved in the torsion [71, 74]. However, these atom types are generally defined by their local chemical environment, which leads to locally-defined torsion types. Therefore, non-local through-bond effects such as conjugation and hypercojugation are difficult to capture [73, 76]. The inability of traditional torsion types to capture such effects and the torsional energy profile’s sensitivity to distal chemical changes are contributing factors to the poor transferability of torsion parameters [73, 76].

To address this issue of poor parameter transferability, many practitioners have employed bespoke parameterization workflows where parameters are fit to quantum chemical torsion drives generated for a specific molecule. Many automatic and semi-automatic tools exist to generate these bespoke parameters, such as GAMMP [32], ffTK [47], Paramfit [7], and QUBEKit [31]. When only a few molecules need to be parameterized, these tools are very useful for aiding researchers in setting up molecular systems in a systematic and reproducible manner. However, the parameters generated from these tools are not meant to be generalizable. The large computational expense of these methods render them impractical when large virtual libraries of molecules need to be parameterized.

Another approach to overcome a lack of transferability in general torsion types is to expand the set of new atom types in an attempt to adequately capture torsion profiles [13]. However, with traditional atom type based force fields, new atom types are generally added in an unsystematic way, and always lead to a proliferation of other force field terms that can result in errors or an explosion in the number of parameters [48].

Atom type-independent force fields seek to overcome both the proliferation of force field parameters and transferability issues by moving away from the restrictions imposed by encoding chemical environments within types assigned to single atoms. The SMIRNOFF force field [48] uses standard cheminformatics SMARTS patterns for direct chemical perception of valence terms, replacing the use of atom types by matching valence terms within their chemical context directly. For torsion types, the SMIRNOFF format matches torsion environments in a hierarchical manner, allowing new, more specific SMARTS patterns to be crafted for a particular class of torsion environments without the need for introducing new atom types. These SMARTS patterns can be created at different levels of granularity for torsion types without worrying about proliferation of parameters impacting other force field terms.

An alternative approach to achieving torsion type generalization by explicitly capturing electronic effects such as hyperconjugation is the Hyperconjugation for Torsional Energy Quantification (H-TEQ) [9, 40, 76] approach. H-TEQ uses chemical principles of conjugation, hyperconjugation, and electronegativity of the atoms involved in the torsion to model torsion energies based on atomic environment without the need to directly fit torsion parameters from quantum chemical calculations.

In this study we combine these concepts to achieve a parsimonious, transferable approach to generalizable torsions for molecular mechanics force fields : To perceive general torsion chemical environments, we make use of the SMARTS torsion typing offered by the SMIRNOFF approach; and to accommodate perturbations to the quantitative torsion barriers in a manner that accounts for distal electronic effects, we make use of the Wiberg Bond Order (WBO), a measure of electronic population overlap between atoms in a bond to model the extent of conjugation. Below, we describe the findings that led to this approach, and explore its ramifications in detail.

### 1.2 Fractional bond orders describe the extent of bonding between two atoms

In quantum chemical formulations, a molecule is treated as a system of individual particles without explicit chemical bonds—nuclei and electrons. Organic chemists, however, think of molecules as atoms held together by covalent bonds. The concept of molecules as chemical graphs is a powerful mental model in chemistry, based on centuries of chemical observations and knowledge that chemists employ when thinking about molecules.

Many quantum chemists such as Pauling [55], Coulson [12], Mulliken [50], Wiberg [79], Mayer [44], Jug [37], and Politzer [57] have worked on bridging the gap between the physical and chemical conception of atoms in molecules by an *a posteriori* analysis of the wave function to arrive at a fractional bond order that is consistent with the chemical concept of the order of a chemical bond [45]. Given that *fractional bond orders* try to make this connection [46], it is not surprising that they capture important chemical properties that can be useful in many applications, specifically force field parameterization. Indeed, in the MMP2 [66], MM3 [4] and MM4 [52] force fields, a Variable Electronegativity SCF derived bond order for π-systems was used to fit bond length, bond force constants, and twofold torsional force constants.

Here, we use the Wiberg Bond Order (WBO) [79], which can be obtained at negligible additional computational cost from a semiempirical calculation like the AM1-BCC [33, 34] calculations commonly used to generate partial charges, and enable accurate extrapolation of torsion force constants to account for distal electronic effects. First we show that WBOs for the central torsion bond are a good indicator of the electron density around the central bond and are linearly related to torsion barrier heights. Then we show that it is possible to use this relationship to interpolate torsion force constants for the same torsion type in different chemical environments. Lastly, we generate SMARTS patterns for the needed torsion types and demonstrate this approach on a set of substituted phenyl rings.

## 2 Results and Discussion

### 2.1 Torsion energy barriers are sensitive to the chemical environment, which can be influenced by remote substituents

In most class I MM force fields, torsions are defined by the atom types for the quartet of atoms involved in the torsion [8, 11, 36, 78]. Atom types encode chemical environments of atoms that usually only incorporate the local environment. However, the quartet of atom types does not always capture all relevant chemistry influencing the torsion profile, especially when the effects are nonlocal; atoms contributing to hyperconjugation, delocalization, or other nonclassical effects may not be part of the quartet involved in the torsion yet can influence the torsional profile [73]. In particular, with conjugated systems, distal electron-donating or -withdrawing substituents can exert a strong effect on torsional barrier height.

Simple examples can help illustrate this, such as the biphenyl example in different protonation states shown in ***Figure 1***A. While the MM torsional profiles are all the same (***Figure 1***D), the QC torsional profiles are different for each protonation state (***Figure 1***C). The torsional energy barrier increases relative to the neutral state for the cation, anion, and zwitterion, in that order. The profile changes qualitatively as well. For the neutral molecule, the lowest energy conformer is slightly out of plane, at 150° and 20°. For the others, the lowest energy conformer is at 180°. In the neutral molecule, the slightly out-of-plane conformer is preferred to accommodate the proximal hydrogens. In the other cases, the increasing double-bond character of the conjugated central bond (shown for the zwitterion in ***Figure 1***B) makes the planar conformer preferred.

**Figure 1.**
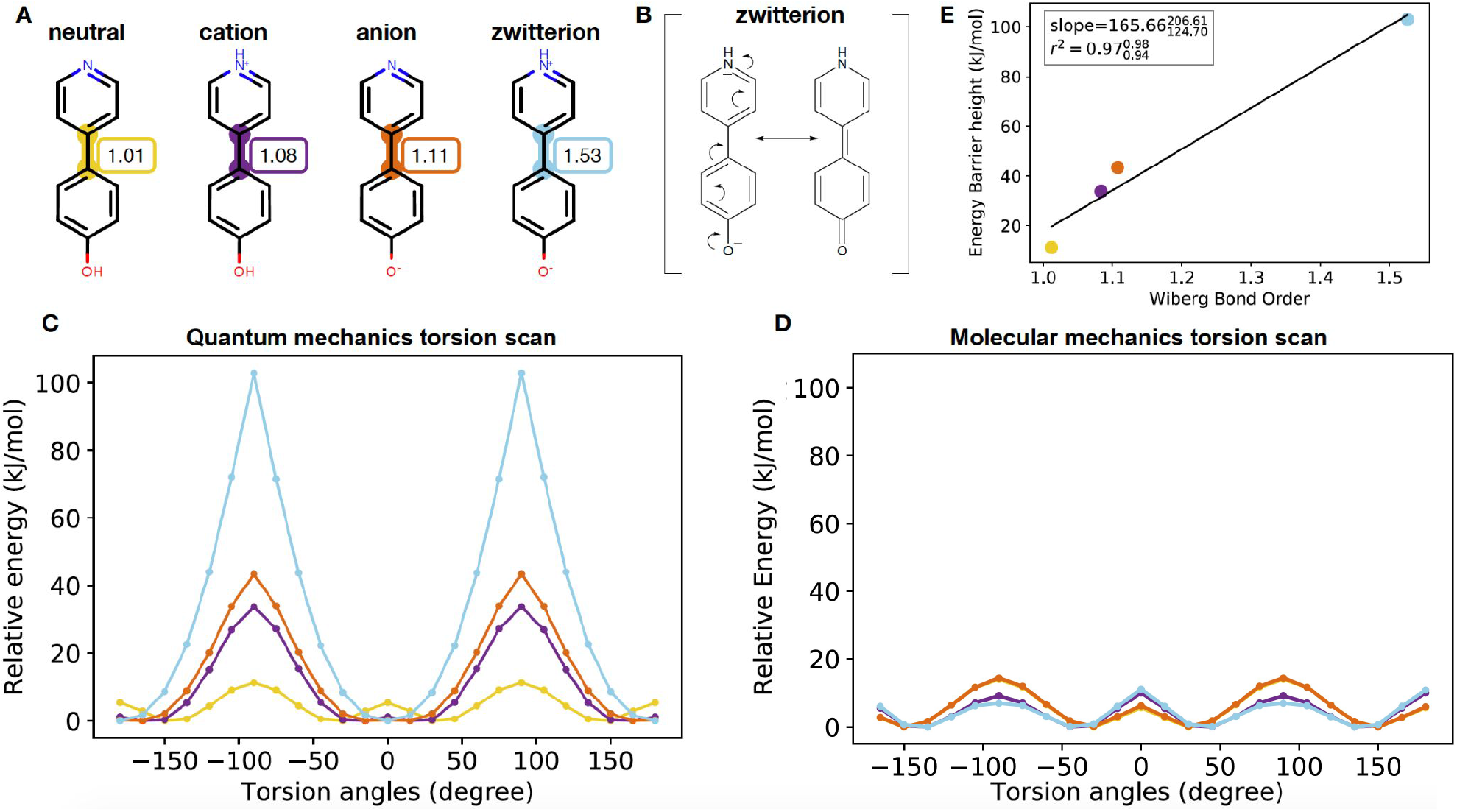
Torsion profiles can be sensitive to remote substituents, and scale linearly with Wiberg bond order about the central bond. **[A]** Biphenyl protonation states and tautomers are shown in order of increasing Wiberg bond order for the central bond. **[B]** The resonance structure of the biphenyl zwitterion shows why the central bond is highly conjugated. The Wiberg bond order and torsion scan for this bond (see **A** and **C**) reflect the nature of this resonance-indced conjugation. **[C]** Relative quantum chemical (QC) energy profile as a function of torsion angle around the central bond computed via QCArchive [65] at B3LYP-D3(BJ) / DZVP level of theory. The colors of the QC scan corresponds to the highlighted bonds in **A**. **[D]**) Same as **C**, but a corresponding molecular mechanics (MM) energy profile computed via the openff-1.0.0 force field. **[E]** Torsion barrier heights scale roughly linearly with Weiberg bond order (WBO) about the central bond. The color of the data points correspond to the highlighted bonds in **A**.

This trend poses problems to generalized torsion force field parametrization. Most general force fields consider the central bond in all tautomers equally rotatable so their MM torsion profiles are all the same (***Figure 1***D), while the QC scan clearly shows that they are not. This illustrates one of the fundamental limits of atom types in classical force fields: At what point in this series should a new atom type be introduced? In this case, remote changes three bonds away from the torsion central bond gradually perturbed the conjugated bond from being highly rotatable to non-rotatable as the conjugation increased.

### 2.2 The Wiberg bond order (WBO) quantifies the electronic population overlap between two atoms and captures bond conjugation

The Wiberg bond order (WBO) is an electronic bond property that is calculated using atomic orbitals (AOs) that are used as basis sets in quantum and semi-empirical methods [45]. WBOs originally started within the CNDO formalism [79], but has been extended to other semi-empirical methods such as AM1 [15] and PM3 [68]. The WBO is a measure of electron density between two atoms in a bond and is given by the quadratic sum of the density matrix elements over occupied atomic orbitals^1^ on atoms A and B:

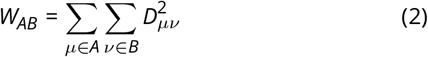

where **D** is the electron density matrix and the quadradic sum is taken over occupied orbitals *μ* and *v* of atoms A and B in the bond.

#### 2.2.1 The WBO is an inexpensive surrogate for changes in the chemical environment that modify torsion profiles

Since the WBO can be calculated from an inexpensive AM1 calculation, is indicative of a bond’s conjugation, and is correlated with torsional energy barrier height, it is attractive to consider using it for interpolating or extrapolating torsion energy barrier parameters. However, WBOs are known to be conformation-dependent [53, 80], so we further investigated this dependence to understand if WBOs will be a robust descriptor suitable for this purpose. In addition, we also investigated the generality of the observed linear relationship between torsional energy barrier and WBO of the central bond (as seen in ***Figure 1***E). In this section, we will first discuss our findings—and present a solution—to the conformation dependence, and then discuss the generality of the WBO linear relationship with torsional barrier height.

#### 2.2.2 Conformation-dependent variance of WBOs are higher for conjugated bonds

Because they are a function of the electron density, WBOs are necessarily conformation-dependent. However, not all bond WBOs change the same way with conformation. As we will show, we found that WBOs for conjugated bonds are more sensitive to conformation and that bonds involved in conjugated systems have WBOs that are correlated with each other. This makes sense in terms of our qualitative understanding of conjugation strength depending on the alignment of π orbitals across the conjugated system: as the change of a distal torsion disrupts the π orbital alignment, the strength of that conjugation on the local torsional barrier decreases.

To investigate how WBOs change with conformation, we used OpenEye Omega [28] to generate conformers for a set of kinase inhibitors (SI ***Figure 3***) and calculated the WBO for each conformation from a B3LYP-D3(BJ) / DZVP [5, 18, 21, 22] geometry optimized calculation using Psi4 [54]. Omega is a knowledge-based conformer generator that uses a modified version of MMFF94s [24] to score conformations. It has been shown to accurately reproduce crystallographically observed conformers in the Platinum benchmark dataset [17].

***Figure 2*** illustrates the results for the FDA-approved kinase inhibitor gefitinib (***Figure 2***A), a representative drug-like molecule. ***Figure 2***B shows the distribution of WBOs for all rotatable bonds color-coded with the colors used to highlight the bonds in gefitinib (Figure 2A). Single carbon-carbon bonds and carbon-nitrogen bonds formed by atoms numbered 10–13 are more free to rotate than conjugated bonds, reflected by low-variance WBO distributions close to unity.

**Figure 2.**
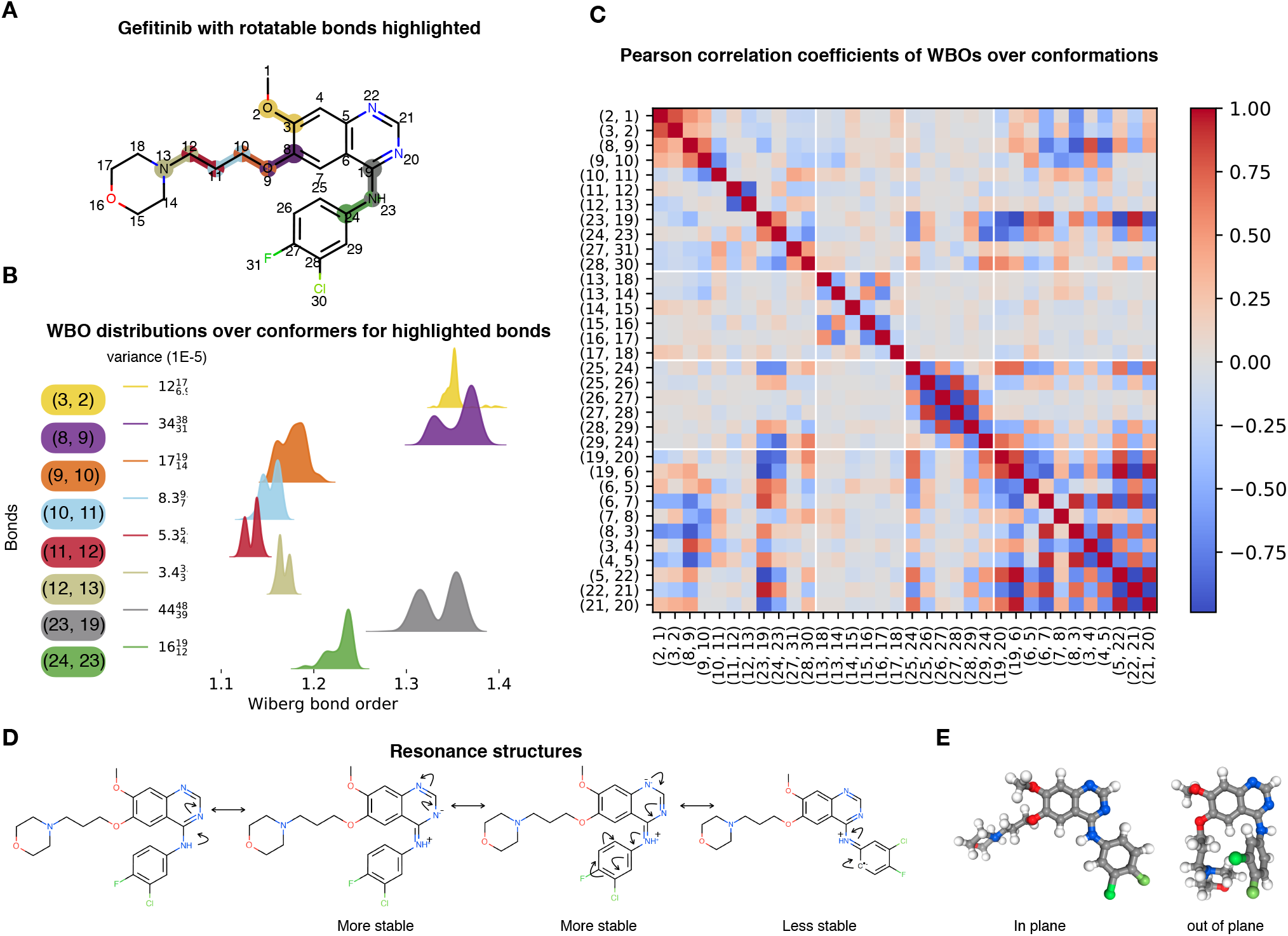
Variance and correlations of Wiberg bond order distributions with respect to conformations are higher for conjugated bonds. **[A]** The FDA-approved kinase inhibitor gefitinib, with its rotatable bonds highlighted and numbered to correspond with figures **B** and **C**. **[B]** WBO distributions over 232 conformations of the highlighted, rotatable bonds. The colors of the distributions correspond to the colors in the highlighted bonds in **A**. The variance and their 95% confidence interval are shown on the left (with exponent base of 1E-5). The single non conjugated bonds (blue, (10, 11), red (11, 12), and olive (12, 13)) have lower variance than conjugated bonds (yellow (3, 2), purple (8, 9), orange (9, 10), grey (23, 19), and green(24, 24)). Conformers were generated with OpenEye Omega. **[C]** Correlation plot of WBOs every bond in Gefitinib against WBOs of all other bonds over 232 conformations. The white lines indicate ring systems. Bonds in conjugated systems have higher correlations between their WBOs (see the aromatic ring systems in the two lower right diagonal squares). Both bonds (23, 19) (grey) and (24, 23) (green) have WBOs that are correlated with their neighboring ring systems, but bond (23, 19) are more correlated with the ring systems than the green bond (24, 23). **[D]** Resonance structures of gefitinib illustrate why the grey bond (23, 19) has higher variance than the green bond (24, 23) even if both bonds are conjugated. When the double bond is on bond (23, 19), the negative charge is on a nitrogen which is the more stable form, vs the resonance structure where the double bond is on (24, 23) with the negative charge on a carbon. **[E]** The conformations of the molecule for the highest WBO and lowest in the distribution. The mode with higher WBOs has bond (23, 19) in plane with the quinazoline heterocycle which allows for stronger conjugation while the mode with lower WBOs has the bond out of plane so there is less electron population overlap in out of plane conformation.

Bonds involving the ether oxygens and aromatic rings (formed by atoms numbered 1–3, 8–10, 19, and 23–24 in ***Figure 2***A and B) exhibit higher variance. It is interesting to note the difference in the WBOs for the conjugated bonds formed by the nitrogen between the quinazoline and chlorofluorophenyl (bonds formed by atoms numbered 19, 23 and 23, 24 in ***Figure 2***A and B). Both of these bonds are conjugated with their neighboring ring systems. However, while the distribution of WBOs for bond 23–19 (the grey distribution ***Figure 2***B) has two clear modes of almost equal weights, the WBO distribution for bond 24–23 has lower variance (***Figure 2***B). This is in agreement with the resonance structures shown in ***Figure 2***D.

The resonance structures that have the double bond on the bond closer to the quinazoline (bond 19–23 ***Figure 2***A and D) are more stable because the negative charge is localized on an electronegative nitrogen atom. When the double bond is on the neighboring 23–24 bond (***Figure 2***D, last resonance structure), the negative charge is localized on an aromatic carbon, which is less stable. The results are similar for other kinase inhibitors examined, as shown in SI ***Figure 3***.

In addition, when we inspected the conformations associated with the highest and lowest WBO in the grey distribution (***Figure 2***B), we found that conformations with lowest WBO on bond 19–23 had that torsion out of plane while the conformation with the highest WBO had the torsion in plane, allowing conjugation (***Figure 2***E). We found similar results from WBOs calculated from quantum chemical (QC) torsion scans with Psi4. Figure 6 shows the WBO for each point in the corresponding QC torsion scans. The WBOs are anti-correlated with the torsional potential energy, in line with chemical intuition: Conjugation stabilizes conformations and leads to more electronic population overlap in bonds [80]. At higher energy conformers, the aromatic rings are out of plane and cannot conjugate, therefore the WBO is lower for those conformers. At lower energy conformations, the rings are in plane and can conjugate so the WBO is higher. We found that these trends are similar when using semi-empirical methods such as AM1 (SI Figure 4). For other levels of QC theory, as well as for the related concept of Mayer bond order [45], the results can quantitatively differ—this is further discussed in the SI section A.1.

#### 2.2.3 Bonds in conjugated systems have highly correlated conformation-dependent WBOs

We found that certain WBOs are strongly correlated or anticorrelated with each other, indicating strong electronic coupling. As bonds in one conformation gain electron population overlap, coupled bonds will decrease in electron population overlap, and vice versa. ***Figure 2***C shows the Pearson correlation coefficient for each bond WBO against all other bond WBOs over 232 conformations for gefitinib. There is a clear structure in this correlation plot: The square formed by bonds from atoms 24–29 shows that the alternating bonds in the aromatic ring (25–29) are strongly anti-correlated with each other (***Figure 2***C.

This trend is different in the ring formed by atoms 13-18, which is not aromatic (***Figure 2***A and C). In this ring, bonds 13–18, 13–14, 16–15 and 16–17 (which involve electron rich atoms O and N) have Pearson correlation coefficients with absolute values higher than for the other bonds in the ring, but lower than the bonds in the aromatic ring. The bonds involved in the methoxy groups (atoms 1–3 and 8–10) are correlated with each other and also correlated to the quinazoline heterocycle, albeit not as strongly. The bonds between the chlorofluorophenyl and quinazoline follow the same trend as their WBO distribution and resonance structures. The bond closer to the quinazoline (bond 23–19) has WBO distribution correlated with the quinazoline, while the bond closer to the chlorofluorophenyl (bond 23–24) is not as strongly coupled with the quinazoline.

The trends are similar for other kinase inhibitors examined, as shown in SI ***Figure 3***.

#### 2.2.4 The electronically least-interacting functional group (ELF) method provides a useful way to capture informative conformation-independent WBOs

As we have shown, the WBO is conformation-dependent, and this dependency can also be highly informative of the electronic couplings within a system. ***Figure 3*** shows the distribution of standard deviations of the conformation-dependent WBO distribution in blue. Most of the standard deviations fall below 0.03, which is encouragingly small. In contrast, when the standard deviation of WBOs is taken over changes to WBO due to remote chemical changes, the stds are higher as shown in the pink distribution in ***Figure 3***. The pink distribution was calculated over 366 distributions of 366 bonds in different chemical environments of 140,602 fragments. These fragments were taken from the validations set of `fragmenter`, an automated approach that uses WBOs to fragment molecules for QC torsion scans with minimal perturbation to the QC torsion profile [67].

**Figure 3.**
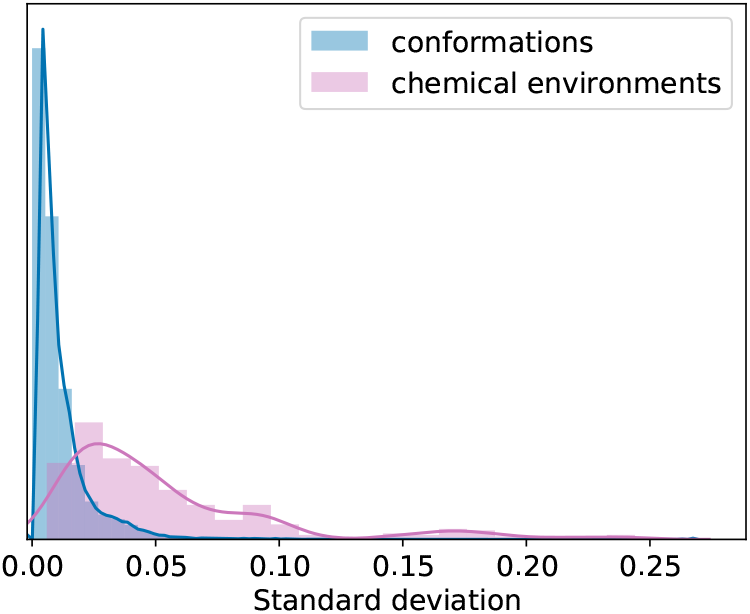
The distribution of standard deviations of WBO distributions is tighter when the distribution is over changed in WBO due to conformations than for changes in chemical environment. The distribution of standard deviations of WBO distributions over conformations is shown in blue. The distribution of standard deviations of ELF10 WBO (explained in subsection 2.2.4) distributions for the same bond in different chemical environments is shown in pink. The changes in WBO due to conformations are smaller than the changes in WBO due to chemical changes around the bond. The fragmenter validation set [67] was used to generate these distributions. The blue standard deviation distribution was calculated over 140,602 WBO dependent distributions (this is the number of individual fragments in the dataset). The pink standard deviation distribution was calculated over 366 distributions of 366 bonds in different chemical environments (of the 140,602 fragments)

It can become computationally expensive to calculate the WBO for all conformations; if we aim to use WBOs as a descriptor for torsional barrier heights in a reproducible way, we need a way to capture informative conformation-independent WBOs. The Electronically Least-interacting Functional groups (ELF) conformation selection scheme implemented in the OpenEye Toolkit ‘quacpac’ module [qua]^2^ resolves the issue of sensitivity of molecular mechanics electrostatic energies from QC derived charges.

The ELF10 method begins with a large set of conformers for the molecule. MMFF94 charges [24] are assigned to the molecule, set to their absolute value, and then single-point Coulomb electrostatic energies evaluated for each conformer. The lowest-energy 2% of conformers are selected, and if there are more than 10, from these the most diverse 10 are selected. For this final conformer set (up to 10 conformers), the AM1 WBOs and charges for each conformer are averaged (by bond and by atom, respectively) over conformers, and the bond charge corrections (BCCs) are applied to the charges [33, 34]. This method yields a set of AM1-BCC atomic partial atomic charges and bond WBOs for the molecule which are relatively insensitive to the initial choice of conformer set, and which mitigate two pathologies of AM1-BCC charges: peculiar charges resulting from strong intramolecular electrostatic interactions (e.g., from internal hydrogen bonds or formal charges) and simple conformational sensitivity.

This method can also be applied to produce WBOs that are insensitive to conformers. To check how well AM1 ELF10 estimated WBOs recapitulate the bond orders, we calculated WBOs from AM1 ELF10 calculations for all bonds in a set of molecules shown in SI ***Figure 4***. The distribution in Figure 4 corresponds closely with bond order. The density at ~0.7 correspond to bonds involving sulfur and phosphorous since these are weaker and longer bonds. The mode at ~1.0 corresponds to C-H and C-C bonds, the mode close to 1.5 corresponds to bonds in aromatic rings, the mode close to 2.0 corresponds to double bonds, and finally the triple bonds form the last peak.

**Figure 4.**
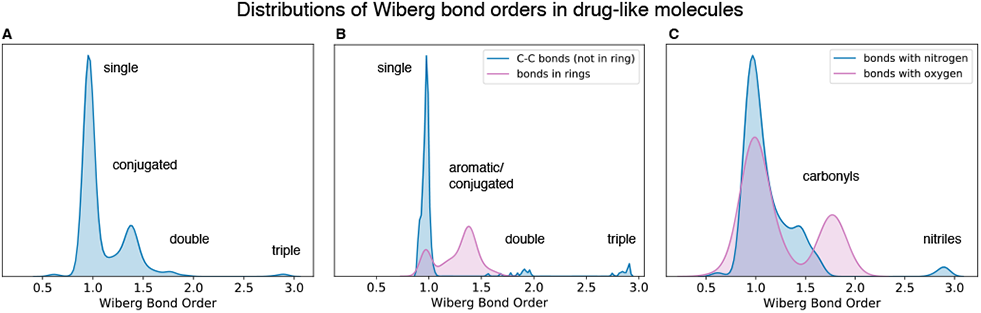
Distribution of WBO in drug-like molecules is concentrated near chemically sensible values. **[A]** The distribution of all WBOs for all bonds in molecules in set. The modes at WBO of one, two, and three correspond to single, double, and triple bonds. The density between one and two correspond to aromatic and conjugated bonds. The mode at ~0.7 correspond to bonds that include sulfur and phosphorous which are longer, weaker bonds. **[B]** The blue distribution includes carbon–carbon bonds that are not in rings. The modes at one, two, and three correspond to single, double and triple bonds. The pink distribution include bonds that are in rings. The mode at one corresponds to single bonds and the density between one and 1.5 are aromatics. **[C]** The blue distribution includes bonds that have either one or two nitrogens. Many of these bonds are conjugated as demonstrated by the density around 1.5. The density at three corresponds to nitriles. The pink distribution include bonds that have oxygens. The mode at two corresponds to carbonyls.

***Figure 4***B and D separate out different kinds of bonds to more clearly illustrate effects captured by the WBO. ***Figure 4***B shows carbon - carbon bonds not in rings (blue) and bonds in rings (pink). The carbon-carbon distribution has distinct modes at one, two and three corresponding to single, double and triple bonds. There is also a smaller mode at 1.5 that corresponds to conjugated bonds. The pink distribution includes bonds in rings and has modes at one and 1.5 which corresponds to aliphatic and aromatic rings, respectively. ***Figure 4***D shows distributions with bonds that have nitrogens (blue) and oxygens (pink). The peaks occur at chemically sensible values; for nitrogen, peaks appear at 1, 1.5, and 3, corresponding to single, conjugated, and triple bonds; for oxygen, peaks appear at and 1 and 2, corresponding to single and carbonyl bonds. For the rest of this section, we focus on the robustness and generalizability of ELF10 WBOs.

#### 2.2.5 WBOs are a robust signal of how torsional barrier heights depend on remote chemical substituents

To investigate how resonance and electronic effects from remote substituents can perturb the torsional energy profile of a bond, we took inspiration from the Hammett equation [25] of reactions involving benzoic acid derivatives. The Hammett equation relates meta- and para-benzoic acid substituents to the acid’s ionization equilibrium constants:

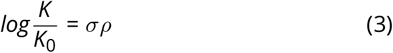

Here, *σ* is a substituent constant, *ρ* is a reaction constant, and *K*_0_ is the reference equilibrium constant when the substituent is hydrogen. This approach aims to isolate the resonance and inductive effects of substituents from the steric effects of a reaction.

Here, we generated a combinatorial set of meta- and para-substituted phenyls and pyridine (***Figure 5***A) with 26 functional groups that cover a wide range of electron donating and withdrawing groups. We then calculated the AM1 ELF10 WBO for the bond joining the functional group to the aromatic ring (highlighted green in ***Figure 5***A) for all functional groups which resulted in 133 (26 * 5 + 3) WBOs for each functional group in different chemical environments. This allowed us to isolate the effect on a bond’s WBO from remote chemical environment changes, defined as a change more than two bonds away, from other effects such as sterics and conformations. The resulting WBO distributions are shown in ***Figure 5***B. (Details on generating and accessing this set are provided in the 4.)

**Figure 5.**
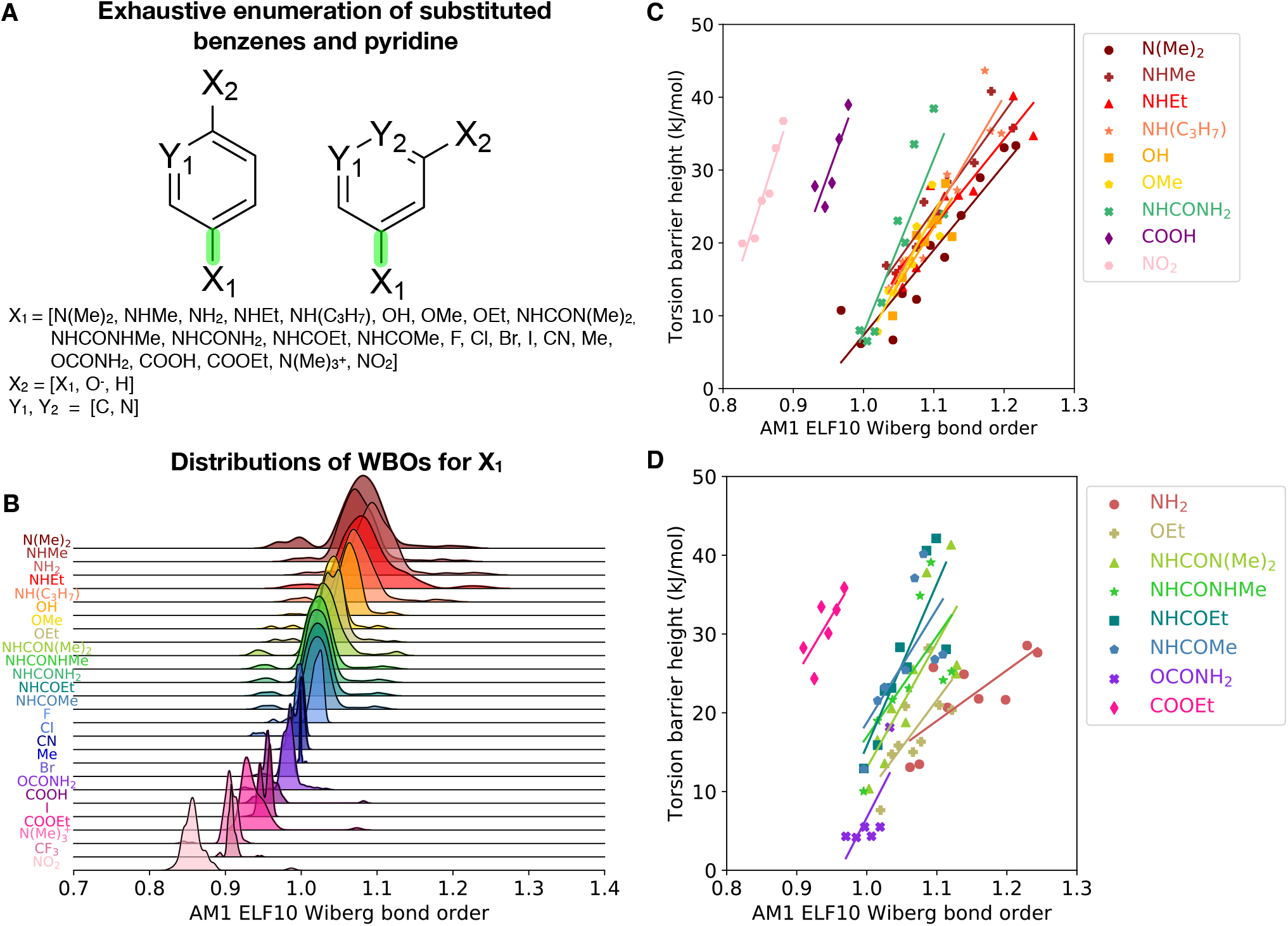
AM1 ELF10 Wiberg bond orders correlate with torsion barrier heights in related molecules. **[A]** Systems and functional groups used in the substituted phenyl set. The functional groups were chosen to span a large range of electron donating and withdrawing groups. **[B]** AM1 ELF10 WBO distributions for the bond between the phenyl ring and *X*_1_ in different chemical environments. **[C]** Selected QC torsion scan barrier heights vs AM1 ELF10 WBOs. These lines all had *r*^2^ > 0.7. **[D]** Same as **C** but these series did not fit the lines as well (*r*^2^ < 0.7).

This substituted phenyl set illustrates several points:

1. Changes in WBO due to modifications of remote substituents can be quite large compared to WBO shifts due to conformational change.
2. The WBOs capture resonance effects.
3. The linear relationship between WBO and torsional energy barrier seems to generalize to many functional groups.

These points are discussed in more detail below.

#### 2.2.6 Changes in WBO due to modifications of remote substituents can be quite large compared to WBO shifts due to conformational change

***Figure 3*** shows two distributions of WBO standard deviations. The blue distribution is the distribution over WBO standard deviations due to conformational changes. The pink distribution is the WBO standard deviations of the same torsion in different chemical environments. These distributions were taken from the validation set for `fragmenter`, an automated approach that uses WBOs to identify fragments that preserve the local torsion environment for reduced-cost quantum chemical torsion scans [67]. When we compare these two distributions, we find that the shifts in ELF10 WBO for remote chemical environment modifications are usually bigger than the shifts in WBO that arise from changes in conformation. This allows us to use ELF10 WBOs as a useful surrogate to capture modifications in chemical environment, and to interpolate or extrapolate their effect on torsion barrier heights.

#### 2.2.7 The WBOs capture resonance effects

It is interesting to note that the decreasing WBOs for more electron donating groups are anticorrelated with increasing Hammett substituent constants. In SI ***Figure 5***, the AM1 ELF10 WBOs of the bonds between the functional group and benzoic acid are plotted against their Hammett meta and para substituent constants (values were taken from Hansch et al. [26]). Functional groups that are more electron donating will have more electron density on the bond attaching the functional group to the benzoic acid. The resonance and/or inductive effect destabilizes the benzoate and increases its pKa, which corresponds to lower substituent constants.

#### 2.2.8 The linear relationship between WBO and torsional energy barrier seems to generalize to many functional groups

To investigate how these long range effects—as captured by the WBO—reflect changes in torsional potential energy barriers, we ran representative QC torsion scans for 17 of these functional groups SI (***Figure 6***). We did not run QC torsion scans for the following functional groups:

- functional groups lacking a torsion (such as halogens)
- functional groups that were congested (such as trimethyl amonium)
- functional groups where the WBOs did not change by more than 0.01 for different functional groups at the meta or para position (such as methyl)

We chose the representative molecules for the 17 functional groups by sorting the molecules within each functional group by their WBO and selecting molecules with minimum WBO difference of 0.02. All of the resulting QC torsion scans are shown in SI ***Figure 6***. Table 1 lists the slopes and associated statistics for the fitted linear models of barrier height as a function of WBO.

**Table 1.**
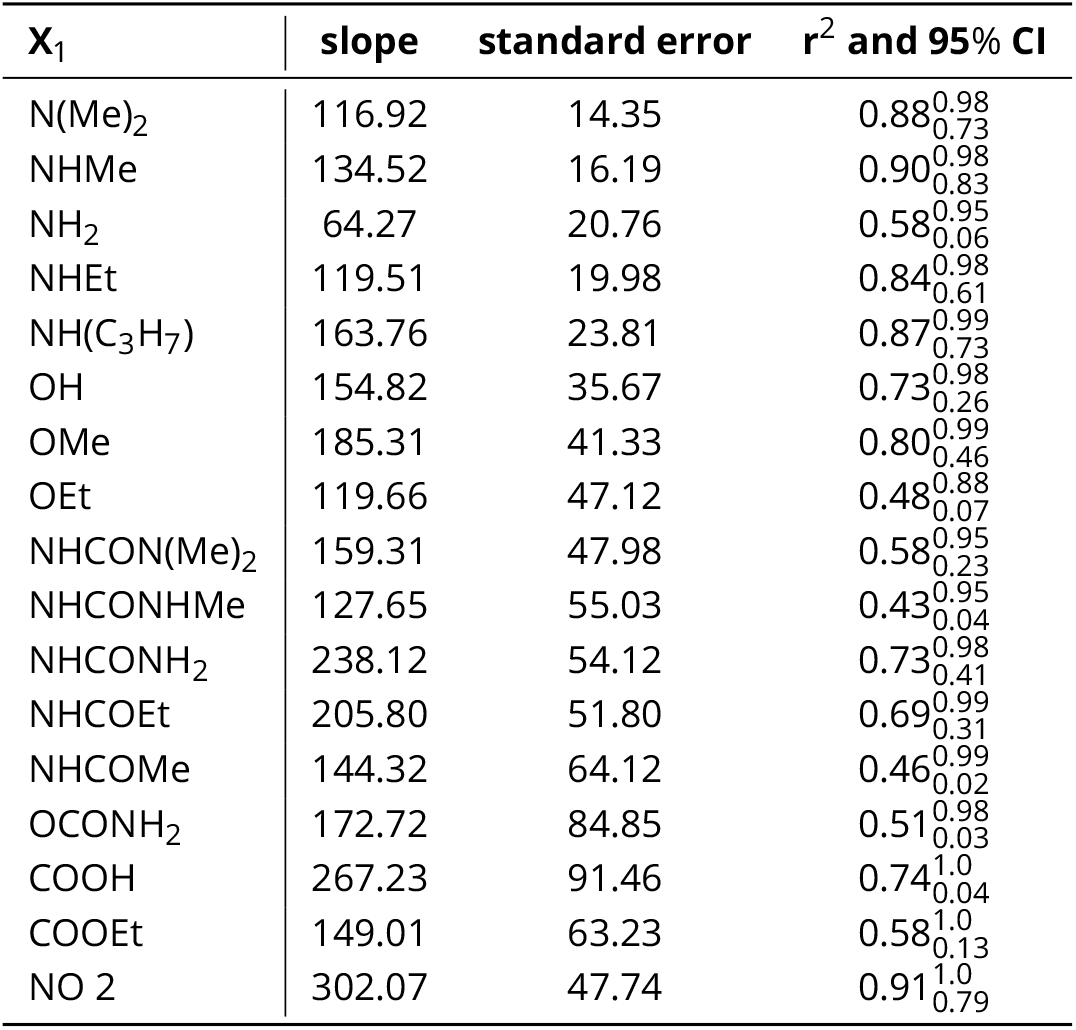
Slope and associated statistics for torsion barrier height vs WBO for selected functional groups.

#### 2.2.9 Conformation-dependent WBOs differentiate between though-space and through-bond non-local effects on QC torsion scans

We show a representative series of quantum chemical torsion scans in ***Figure 6***, and the corresponding conformation-dependent WBO for each conformation in the scan. A QC torsion scan contains contributions of through-bond effects such as conjugation and/or hyperconjugation, as well as through-space effects such as steric and electrostatic interactions [20, 76]. In this section, we show how WBOs can be used to characterize through-space and through-bond effects in QC torsion scans.

**Figure 6.**
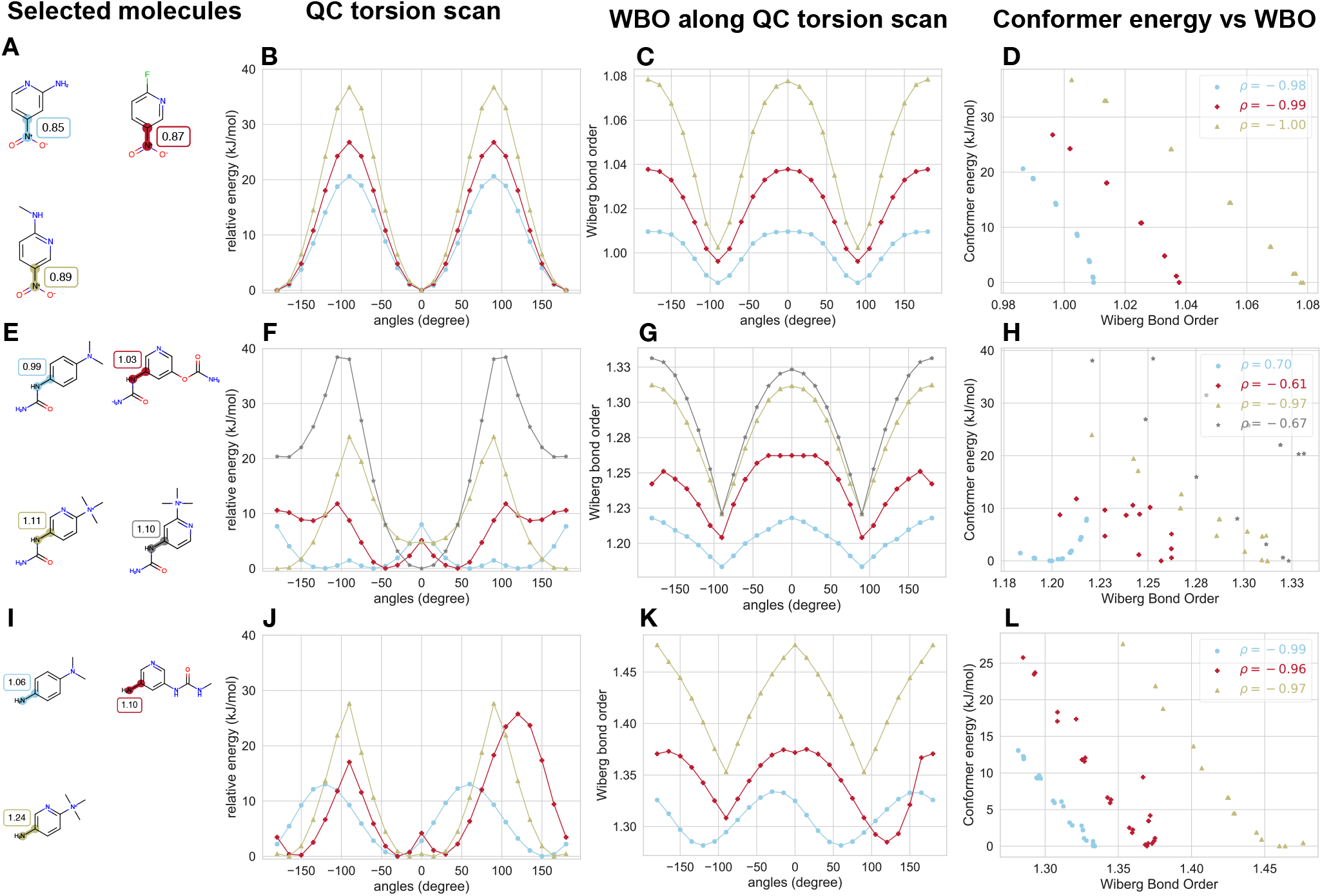
Wiberg bond orders are anticorrelated with conformer energies from QC torsion scans. **[A]** Selected series of molecules with central torsion bonds connecting the nitro group to the phenyl ring highlighted and labeled with AM1 ELF10 WBOs. **[B]** QC torsion scans for nitro series in different chemical environments shown in **A**. The color of the scans correspond to the colors of the highlighted bonds in **A**. **[C]** Wiberg bond orders calculated at each point in the QC torsion scan using the same level of theory. **[D]** Energies of conformers produced during the QC torsion scan plotted against it WBO. All molecules in this series have WBO that are anti-correlated with their QC torsion scan conformer energies. Pearson correlation coefficients (*ρ*) are shown in the upper right legend. **[E]** Same as **A** but with urea at the *X*_1_ position. **[F]** Same as **B** but for urea in a series of different chemical environment. Both profiles and energy barriers change with AM1 ELF10 WBOs. In addition, the grey scan has higher energy barriers than the olive scan but its ELF10 WBO is lower. **[G]** Same as **C** but for urea. Here, the WBO scans all have the same profiles while the QC torsion scan does not. **[H]** Same as **D** but here the WBO scans are not always anti-correlated or not as strongly anti-correlated. The WBO scan profiles do not change because the changes in the QC torsion scan captures spatial effects while the WBO scans capture conjugation. **[I]** Same as **A** with amino at the *X*_1_ position. **[J]** Same as **B** but for amino in different chemical environments. While the gold scan is symmetric around 0°, the red and blue scan are not. The Blue scan is shifted and the red scan has a higher barrier on one side. **[K]** Same as **C** but for amino. Here the WBO scans are anticorrelated with the QC torsion scans even if the QC scans have different profiles. **[L]** Same as **D** but for amino. Here the WBO scans are anticorrelted with the QC torsion scans.

As a molecule is rotated about its bond to generate a QC torsion scan, changes in conjugation and/or hyper-conjugation, the conformation of the rest of the molecule, and non-local, through-space interactions contribute to the potential energy surface that is then used to fit MM torsion parameters. The relaxation of the orthogonal degrees of freedom can result in hysteresis of the torsion profile. To avoid this issue we used waterfront propagation to minimize hysteresis [61] The torsion parameters in classical force fields are intended to include corrections for both conjugation—a through-bond electron delocalization phenomenon that is not well-modeled in classical force fields—and the simple treatment of 1–4 non-bonded steric and electrostatic interactions. To increase their transferability, torsion parameters should not include corrections for non-bonded interactions beyond 1–4 atoms. However, in general, it is difficult to separate the contributions of sterics and conjugation in a QC torsion scan. In this section, we characterize steric and conjugation and/or hyperconjugation contributions to QC scans using the corresponding WBO scans. Below is a summary of these observations from the substituted phenyl torsion scans:

1. When QC torsion scans are anti-correlated with conformation-dependent WBOs calculated for conformations in the scan, differences in QC torsion scans for the same torsion types in different chemical environments are a result of non-local, through-bond effects.
2. If changes in QC torsion scans are not accompanied by highly anti-correlated WBOs, the changes are due to non-local, through-space, effects.
3. When AM1 ELF10 WBOs do not obey the linear relationship with barrier heights observed in ***Figure 5*** for the same torsion type in different chemical environments, they are generally caused by non-local, through-space interactions.

***Figure 6*** shows three series of QC torsion scans for the same torsion type in different chemical environments to illustrate this. ***Figure 6*** A, E and I show three torsion types (nitro, urea and amino) in different chemical environments with their associated AM1 ELF10 WBO. ***Figure 6***B, 6F, and 6J show their QC torsion scans, Figure 6C, ***Figure 6***G, and 6K show their conformation-dependent WBOs along the torsion scan, and ***Figure 6***D, 6H, and 6L show the correlations between conformer energies and their conformer-dependent WBOs.

***Figure 6***A-D show what is generally the expected behavior of QC torsion scans for the same torsion types in different chemical environments as shown in ***Figure 5***. The QC torsional profiles in ***Figure 6***B are all the same, while the torsional barrier heights increase with increasing AM1 ELF10 WBOs. WBOs calculated for every conformer of the QC torsion scan generate a profile that is generally anti-correlated to QC torsion scans, as shown in ***Figure 6***C and 6D. This is in line with chemical intuition: Increased conjugation is a result of increased electron population overlap, which stabilizes conformations, decreasing their energy.

The second derivative of WBOs along a QC torsion scan profile also changes depending on how strongly the bond conjugates at its lowest energy conformation. For bonds with higher AM1 ELF10 WBOs (***Figure 6***A and 6E) which indicates increased conjugation, the rate of change in the WBO scans are higher than for the same torsion types in environments where the bond does not conjugate as strongly (***Figure 6***C and 6G). This also makes sense with respect to chemical intuition: At high energy conformations, where electronic orbitals are not oriented to conjugate, WBOs of the same bond in different chemical environments will be closer to each other than at lower energy, where electronic delocalization is modulated by distal chemical substituents. In other words, conjugation is disrupted similarly for the different chemical environments, but the extent of conjugation is different because of different remote substituents.

***Figure 6***E-H shows a different series of the same torsion type in different chemical environments. There are several differences between this example and the example discussed in the previous paragraph. One, while the torsion types are equivalent for all four molecules shown in ***Figure 6***E, the QC torsion scans do share the same profile. Relative heights of minima and maxima differ, or new minima and maxima are observed. In addition, their corresponding conformation-dependent WBO scans are not as strongly anti-correlated with the torsion scans (***Figure 6***H) as in ***Figure 6***D. Interestingly, the WBO scans share similar profiles (Figure 6G). In this example, the urea is a more bulky functional group than the nitro and amino group in ***Figure 6***A and 6I. The red and grey molecules also have bulky groups at the meta position relative to the torsion being driven. This creates distinct steric clashes for the different molecules, changing their QC torsion energy profiles relative to each others. However, the conformation-dependent WBO scans have similar profiles for the molecules in this series because the non-local through-bond effects are similar. These WBO scans are not as anti-correlated to the QC scan as in the previous examples (−0.98, −0.99, −1.00 vs 0.7, −0.61, −0.97 and −0.67). The blue scan is actually correlated instead of being anti-correlated. While the electrons can conjugate when the torsion is in a planar position, the clashes of the proximal hydrogen and oxygen increase the energy. This small barrier at 0° does not exist in the grey and gold scan because the trimethylamonium is electron withdrawing, that the urea group can better conjugate with the phenyl ring; this stabilization is sufficient to overcome the steric interactions.

This series also shows an example of molecules that do not follow the trend shown in ***Figure 5*** where increasing AM1 ELF10 WBO corresponds to increasing torsion barrier heights. The grey torsion scan (***Figure 6***F) has torsion barrier heights that are ~15 kJ/mol greater than the gold torsion scan, while its AM1 ELF10 WBO is lower than the gold molecules (1.10 vs 1.11). This occurs because the trimethylamonium is a bulky group at the meta position, and it interacts with the carbonyl in the urea which causes the the barrier heights to increase. We observed this trend of bulky groups on the meta position clashing in most other cases where the AM1 ELF10 WBOs did not follow the trend of increasing torsion barrier heights (SI ***Figure 6***).

***Figure 6***I-L show yet another series of torsion scans that exhibit different behaviors than both examples already discussed. In this example, the QC torsion energy profiles shown in ***Figure 6***J are all distinct. Notably, the reflection symmetry around zero torsion angle is lost in the blue and red scan, albeit in a different manner. The blue scan shifted such that the minima are not at 0° and 180° but the barrier heights are equivalent. In the red scan, one barrier is ~10 kJ/mol higher than the other. However, in these scans, the corresponding WBO scans are anti-correlated with the torsion energy profile scans (***Figure 6***L). In these cases, these changes are due to changes in conjugation, rather than non-local steric interactions. All of these molecules contain a trivalent nitrogen that can assume both a pyramidal and planar conformation, depending on the amount of electron density in the lone pair. The more electron density there is on the lone pair, the greater the angle of the pyramidal nitrogen will be. If the lone pairs conjugate with other *π* electrons, the trivalent nitrogen adopts a planar conformation for two reasons: *(1)* To accommodate the conformation needed for conjugation, and *(2)* because there is less electron density in the lone pair of the nitrogen now that the electrons are delocalized.

***Figure 7*** shows how the angle of the trivalent nitrogen involved in the torsion changes over the course of the torsion scan. The blue scan, which corresponds with the molecule with a low AM1 ELF10 WBO relative to other bonds of the same torsion type (1.06), does not conjugate with the phenyl ring and remains in a pyramidal with an angle of ~35°. This creates a chiral center, causing loss of reflection symmetry around zero torsion angle [30], as seen in the blue torsion scan in ***Figure 6***J. The red scan—which also corresponds to a molecule with a lower AM1 ELF10 WBO relative to other bonds with this torsion type (1.10)—does not conjugate and remains pyramidal throughout the entire scan. However, in this scan, the chirality of the pyramidal nitrogen does flip, but then does not flip again at 100°, which can explain why the barrier heights are so different from one another. Lastly, the gold scan—which corresponds to a molecule with a relatively high AM1 ELF10 WBO (1.24)—does conjugate at a torsion angle of zero and the trivalent nitrogen becomes planar with an improper torsion angle of zero.

**Figure 7.**
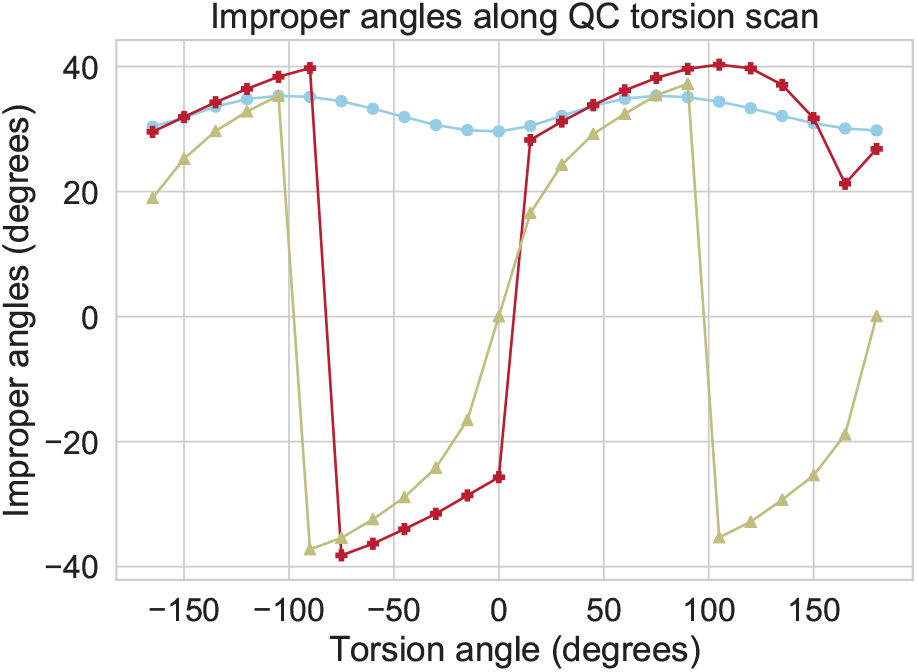
Improper torsion angles can be coupled with torsion angles being driven in torsion scans. Improper angles of pyramidal nitrogen involved in the torsion scan in ***Figure 6***J. In the blue torsion energy profile scan, the trivalent nitrogen is in a pyramidal conformation for the entire scan and the pyramid does not inter-convert. In the red scan, the trivalent nitrogen is also in a pyramidal conformation for the entire scan, but the pyramid does inter-convert. In the gold scan, the pyramidal nitrogen becomes planar at torsion angles 0° and 180° as the amino group conjugates with the phenyl.

The general trend we found when looking at other molecules in the subsituted phenyl set (SI ***Figure 6***) was that when trivalent nitrogen were involved in a torsion scan, changes in the torsion profile relative to the same torsion type in different chemical environments, were due to changes in non-local through-bond effects. These changes were also universally reflected in the corresponding WBO scans that remained anti-correlated with the QC scan. Specifically, torsions involving trivalent nitrogens with relatively lower AM1 ELF10 WBOs for the central bond, were more likely to exhibit such changes. In all cases where such changes were observed, the trivalent nitrogens did not form planar conformations at appropriate points in the scan to conjugate with the neighboring phenyl ring.

Calculating conformation-dependent WBOs for a torsion scan is computationally inexpensive relative to a full QC scan, and the information gleaned from it can be helpful in differentiating non-local through-bond and through-space effects. In general, when the conformation-dependent WBO scan is not strongly anti-correlated with the QC torsion scan, the QC scan contains through-space steric effects. When the conformation-dependent WBO scans are strongly anti-correlated with QC scans, especially when those profiles have loss of symmetry and the torsion atoms include a trivalent nitrogen, the changes in QC torsion profiles relative to QC scans of the same torsion type in different chemical environments are usually a result from through-bond effects and need to be incorporated in classical torsion force field parameters.

### 2.3 Wiberg bond orders can be used to interpolate torsion parameters

Traditional torsion parameters do not consider remote chemical changes and their effects on the torsional barrier height (***Figure 1***D). Therefore traditional parameters incorrectly assign force constants to seemingly similar chemistries. ***Figure 1*** and ***Figure 5*** illustrate that WBOs are able to capture changes in chemical environments. In addition, we have shown that WBOs are linearly correlated with torsion barrier height for similar torsions in different chemical environments (***Figure 5***C and D). In this section we show proof of concept that we can use this linear relationship to interpolate torsion parameters, thus capturing the trends observed in ***Figure 5***C and D. Interpolated torsion parameters utilize WBOs to determine more accurate force constants that consider the electronic properties of not only the substructure of the torsion, but also the entire molecule. Using the linear relationship between the WBO and torsional barrier heights in a series of molecules with the same torsion type, the torsion force constant is determined for every torsion in the series by interpolating along the line. Support for utilizing calculated WBOs via AM1 ELF10 is now included in the [SMIRNOFF specification](https://github.com/openforcefield/openff-toolkit/blob/master/The-SMIRNOFF-force-field-format.md). The [OpenFFToolkit](https://github.com/openforcefield/openff-toolkit) implements the use of these parameters and calculated WBOs to parameterize interpolated torsions for a set of molecules with the same torsion in different chemical environments. Detailed usage and implementation details can be found in the 4.

In order to show that using WBO to ineterpolate torsion force constants can improve torsion parameters in a force field, we performed a proof of concept experiment with a set of SMARTS patterns. For this experiment, we used the substituted phenyl dataset shown in Figure 5 because of how well the data captures remote chemical changes effects on torsion QC scans. The SMARTS patterns are used in the force field to hierarchically assign torsion parameters through a chemical substructure search, as encoded in the SMIRNOFF format [48]. To determine candidate SMARTS patterns to capture these trends, we started with the pre-existing SMARTS patterns for the torsions types in Open Force Field 1.3.0. We assigned these torsion types to the substituted phenyl set and looked at the resulting torsion barrier heights as a function of WBO. Particularly, we examined the data in ***Figure 8*** showing torsion barrier heights vs WBO, and then color-coded the data based on which parameters were used for the different torsions. A good set of SMARTS patterns of torsion types ought to correctly capture the different trends shown ***Figure 5***C and D. We then used our chemical intuition to refine different SMARTS patterns/torsion types and come up with proposals which increasingly separated the different trends in ***Figure 8***, eventually arriving at the different SMARTS patterns tested here. For example, our chemical intuition would lead us to separate out C-N, C-C and C-O central bonds, and we felt carboxylic acids would probably need to be separated from other C-C central bonds, so we tested these combinations and observed how their use would affect the separation of the different trends in Figure 8. Each attempt then led to further improvements. Particularly, if a proposed SMARTS pattern was too general, we used that parent SMARTS pattern to create more specific patterns. Our goal, here, was not to obtain a perfect and general force field, but simply to separate the different trends in our available data for this proof-of-concept test. We named the the interpolated torsion parameters in the experiment with the torsion ID TIG1-10 which stands for “Torsion Interpolated General”, which is how we will identify and refer to the proposed interpolated torsion parameters in this paper. Our final chemical groupings are shown in ***Figure 8***. Some of the trend lines such as TIG9 and TIG10 have almost identical slopes as shown in ***Figure 5***C and 5D, but we separated them because of chemical differences of the Oxygen; ether in TIG9 and alcohol in TIG10, though further refinements may be needed as part of generalizing this work.

**Figure 8.**
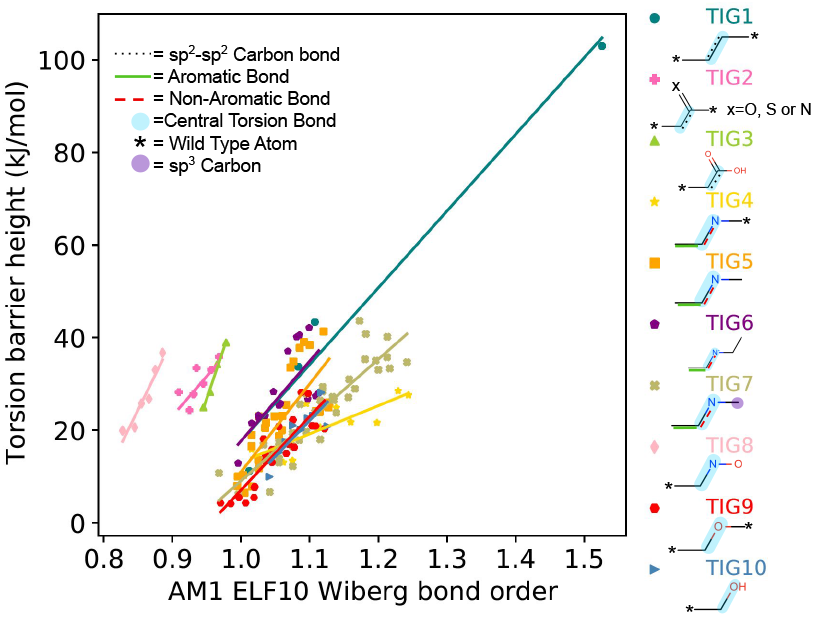
A set of interpolated torsion parameters applied to the substituted phenyl dataset. The set of interpolated torsion parameters (TIGs) that were generated based on chemical similarities and existing torsion parameters in the Open Force Field v1.3.0. The plot shows the AM1 ELF10 Wiberg bond order versus the torsion barrier height (kJ/mol) for the set of molecules in the substituted phenyl dataset. The points and the corresponding interpolation lines are colored and grouped based on their TIG (Torsion Interpolated General) parameter ID matches. The fitting data and SMARTS patterns are reported in table 2. The unmarked carbons represent wild type carbons. Wild type bonds and atoms indicate that there can be any atom or any bond type. There are single, double, *sp*^2^-*sp*^2^, aromatic, non-aromatic, and wild type bonds for the parameters. Further details regarding the SMARTS patterns, wild card bonds and associated statistics are included in Table 2. It is important to note that this data shows the barrier height for the full torsion profile, whereas the desired barrier height in the force field is typically the barrier height for the residual, as discussed in the text.

For the substituted phenyl dataset, we needed to create 10 interpolated torsion parameters to capture all of the chemical diversity in the set. We named these parameters Torsion Interpolated General (TIG) parameters 1–10. We show the slope, standard error, r^2^, and 95% confidence interval for each TIG in Table 2.

**Table 2.**
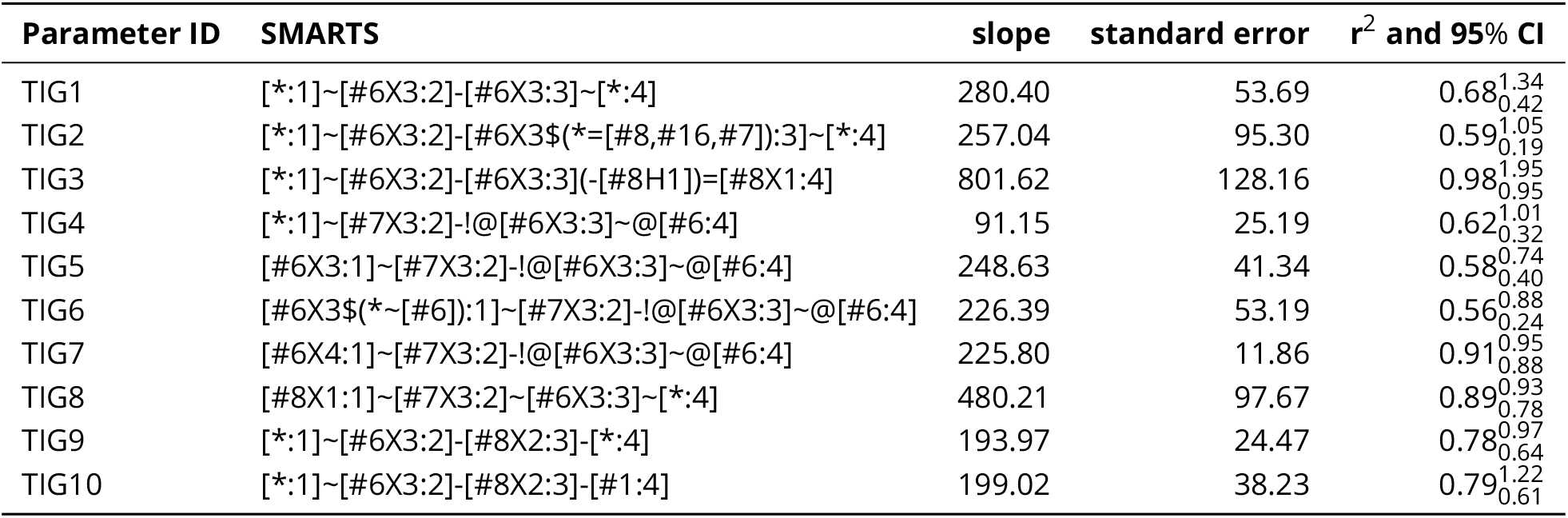
Slope and associated statistics for torsion barrier height vs WBO for interpolated torsion parameters grouped by SMARTS patterns. The results in this table are from performing a linear regression fit on the ELF10 AM1 Wiberg bond order versus the torsion barrier height using SciPy. The linear regression fits are grouped by torsion parameter matches of molecules from the substituted phenyl dataset. As in Figure 8, these fits are for trends capturing the full torsional barrier height, and are not the force constants used in the force field (as discussed in the text).

#### 2.3.1 Interpolated torsion parameters reduce errors in MM torsion parameters without introducing more torsion types

After coming up with a proposed set of new torsion types (Figure 8; Table 2), we wanted to test whether these new torsion types would actually improve the accuracy of describing torsions. The data in Figure 8 and Table 2 shows how these parameters could capture trends in the *full* MM potential as torsions are driven; however, MM torsional potentials instead need to capture the *residual* – the difference between the quantum mechanical torsional potential and the MM energy profile for that same torsion drive, without any MM torsional potential. If the MM energetics accurately captured QM torsion energetics without any MM torsional potential, no MM torsional potential would be needed. This means that the torsional potentials resulting from a fit might need to capture somewhat different energetic behavior.

To test this, we used ForceBalance to fit a force field including the TIG1-TIG10 interpolated parameters with the typing illustrated in Figure 8. Using interpolated torsion parameters, we are able to reduce errors in the MM fitted torsion parameters without the addition of more torsion types. ***Figure 9***A shows the root mean square error (RMSE) of MM vs QM torsion scans for the torsions in all TIGs using torsion parameters from OpenFF 1.3.0 (blue bar) and interpolated torsion parameters (pink bar). In general, interpolated torsion parameters reduce the MM vs QM torsion scan RMSE.

**Figure 9.**
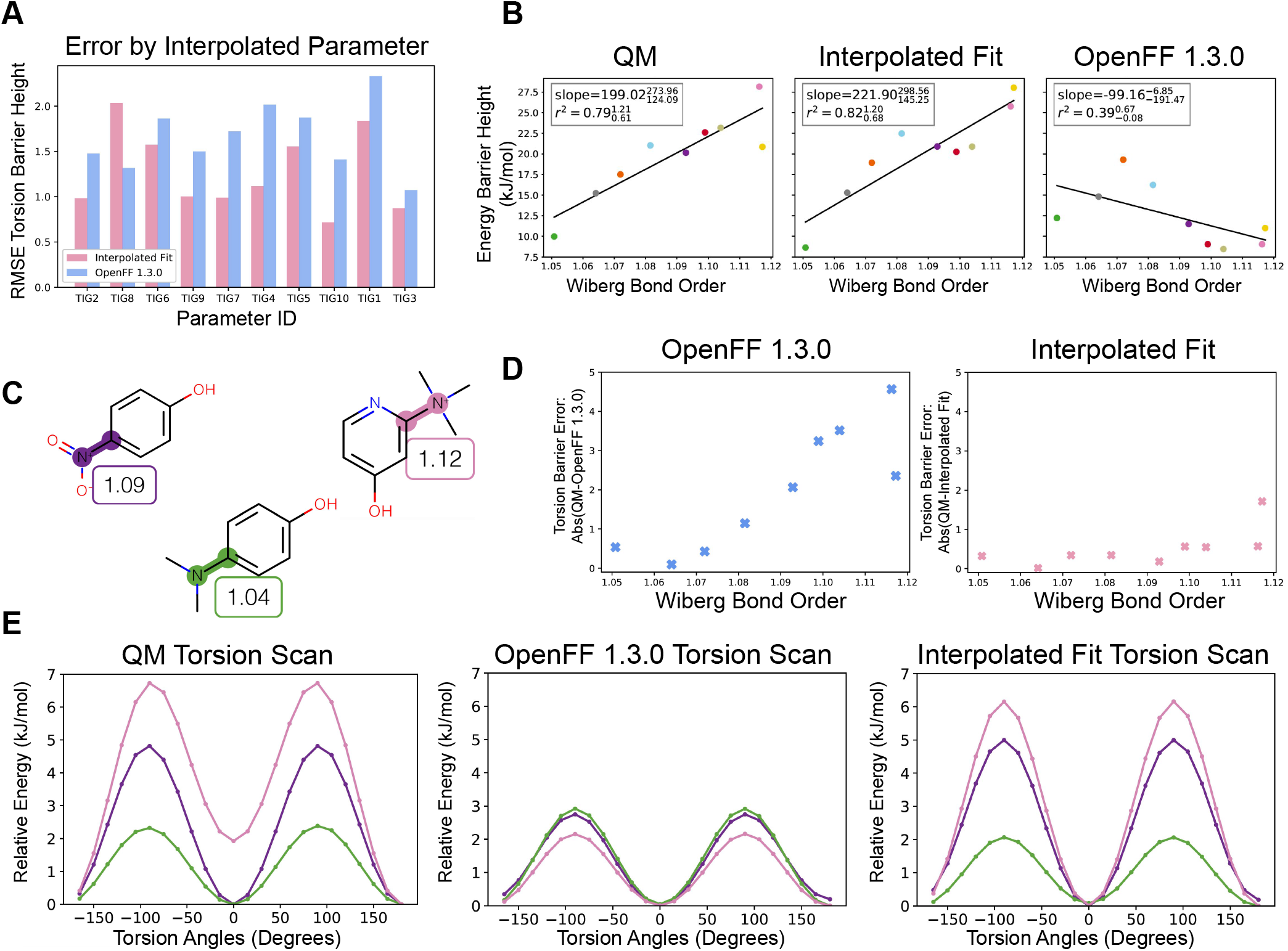
Representative of QC and fitted torsion scans for TIG10 parameter. Using ForceBalance, we performed a fitting experiment to refit the interpolated torsions proposed in ***Figure 8*** using the QC torsion scans from the substituted phenyl dataset as target data. Here, we depict the fitting results for the interpolated torsion parameter with parameter ID TIG9 and the SMARTS pattern `[*:1] [6X3:2]-[8X2:3]-[*:4]`. **[A]** The RMSE for the torsion barrier height calculated from the QM and MM difference of the interpolated fit and OpenFF 1.3.0 fit. The blue bars are the RMSE of the OpenFF 1.3.0 torsion fit and the pink bars are the RMSE of the interpolated torsion fits. The general trend is lower error for the interpolated fit compared to OpenFF 1.3.0 [60]. **[B]** The AM1 ELF10 Wiberg bond order versus torsional energy barrier height for the molecules in the substituted phenyl set that use the TIG10 parameter. The subplots show the QM, interpolated fit, and OpenFF v1.3.0 results for the same set of molecules. Each molecule uses the same respective color in the QM, interpolated fit and OpenFF v1.3.0 subplots. We added a line of best fit to illustrate the improvement of the MM torsion barrier by using an interpolated torsion. **C** Three selected molecules that use TIG10 with their central torsion bond highlighted and ELF10 WBO value. The highlighted color matches with the MM torsion scans in panel **E** and data points in panel **B**. **D** The OpenFF 1.3.0 and Interpolated fit torsion barrier height difference with the QM torsion barrier versus the AM1 ELF10 WBO. **E** QM and MM torsion profiles for three molecules from **C**. The MM torsion profiles consist of the energies from OpenFF v1.3.0 and the interpolated fit force field.

In ***Figure 9***E, we show an example of the improved fit when using interpolated torsion parameters. The torsion profiles in ***Figure 9***E correspond to the torsion about the central bonds highlighted in the three molecules in (Figure 9C). All three of these molecules use the TIG10 parameter, but have varying torsional barrier heights. The OpenFF 1.3.0 MM energies show little variation in the torsion barrier height, but when using torsion interpolation as shown in the interpolated fit torsion scan, we are able to capture torsion barrier height more accurately.

Overall we were able to improve the MM torsion scans relative to the QM scans by using interpolated torsion scans as shown in ***Figure 9***. In ***Figure 9***B and ***Figure 9***D we show the results of the fit for all molecules that use TIG10 in more detail. ***Figure 9***B shows the improvements of the torsion barrier heights on the interpolation line for this parameter. The first panel shows the QM torsional barrier height vs the WBO of the central bond, the second panel shows the MM torsional barrier height derived from interpolated torsion parameters vs the bonds’ WBOs, and the third panel shows the same data as the second but using the torsion parameters from OpenFF 1.3.0. Using OpenFF 1.3.0 torsion parameters resulted in a trend line that is opposite of what we see with QM torsion scans. We were able to correct this by fitting interpolated torsion parameters and predict more accurate MM torsion barrier heights.

***Figure 9***D illustrates that interpolated torsion parameters reduce the error in the MM torsion barrier heights. The first panel shows the absolute difference between the QM and MM torsional barrier height vs WBO. As the WBO increases, the absolute error increases as well. In the second panel, the interpolated torsion parameters alleviate most of this error.

We see a decrease in performance for TIG8 in ***Figure 9***A, as judged by the RMSE value. Although the RMSE increases, by visual inspection of the MM torsion profiles of the interpolated torsion fit and OpenFF 1.3.0 SI ***Figure 7***, we see that the interpolated torsion fit MM profiles show an overall decrease in torsion barrier heights but a better separation of the torsion profiles which is more representative of the QM torsion profiles SI (***Figure 7***). There is also an improvement of the slope of the torsion barrier height as a function of WBO, and of the *r*^2^ value for the interpolated fit in comparison to OpenFF 1.3.0 SI (***Figure 7***).

In order to generalize interpolated torsion parameters, more work is required. In particular, we need a better understanding of which chemical groups show a strong trend between WBO and torsion barrier height, and a good way to separate such trends from confounding factors such as the effects of strong neighboring steric interactions which might obscure trends. Studies that use small simple molecules with conjugation may be a promising avenue for the development of interpolated torsion parameters. When studying WBO versus torsion barrier trends, it appears important to limit studies to simple molecules or carefully select the series which are studied, because more complicated chemical systems often involve non-bonded interactions which can obscure the relationship between WBO and torsion barrier height. The present work shows how WBO can correlate with torsion barrier height for several simple series, but more work is needed before we can produce torsional parameters which will generalize well across a broader region of chemical space.

## 3 Conclusion

We have shown that the WBO is a simple yet highly informative quantity of a bond’s chemical environment. The conformation sensitivity of WBOs is also informative: variance over conformers reports on conjugation, and correlations between bonds captures conjugated systems in molecules. When the WBO is calculated along a QM torsion energy profile scan, it can inform on which effects in the profile are due to steric interactions vs through-bond effects. When conformation dependence is minimized via the ELF10 method, the WBO still remains highly informative with respect to resonance effects, the extent of conjugation, and its relationship with torsion barrier heights. In addition, we also showed that trivalent nitrogen improper torsion angles are coupled with torsion scans.

Since WBOs are linearly related to torsion barrier heights, we demonstrate in a proof of concept that WBOs can be used to derive interpolated torsion parameters such that fewer torsion types are needed to reduce the error in capturing QC torsion energy profiles. We show that this approach allows us to more accurately capture QM torsion energy scans of similar chemistries with different torsion profiles without a proliferation of torsion types. While in this study we only showed in sample improvements of MM torsion scans, we believe that this approach can also be used for fitting out of sample MM torsion parameters of the same torsion type thus reducing the number of expensive QC scans needed to fit torsion parameters.

## 4 Detailed Methods

### 4.1 QCArchive data generation and archiving

The MolSSI QCArchive [64] project [http://qcarchive.molssi.org] is a platform for computing, organizing, and sharing quantum chemistry data. Computations with this platform automatically record all input and output quantities ensuring the reproducibility of all computations involved. In the case of computing with the MolSSI QCArchive instances, all data is automatically hosted at http://qcarchive.molssi.org and can be queried using QCPortal [https://github.com/MolSSI/QCPortal].

#### 4.1.1 Submitting computation to QCArchive

All scripts used to submit calculations for the datasets in this paper are hosted on GitHub online at https://github.com/openforcefield/qca-dataset-submission.

The submission scripts used for the kinase inhibitor and substituted phenyl datasets are listed below:

1. [Kinase inhibitor dataset](https://github.com/openforcefield/qca-dataset-submission/tree/master/submissions/2019-11-27-kinase-inhibitor-optimization)
2. [Substituted phenyl dataset](https://github.com/openforcefield/qca-dataset-submission/tree/master/submissions/2019-07-25-phenyl-set)

#### 4.1.2 Details on QC and MM torsion scans

All torsion scans were computed with TorsionDrive [61], which makes choices of new constrained optimizations to evaluate. The required constrained optimizations were then computed with the geomeTRIC [75] standalone geometry optimizer interfaced to the QCEngine [qce] project.

To ensure a fair comparison between the QC and MM torsion scans, the only change in the torsion scan procedure was to switch out the program, which evaluated the gradient at each step in the geomeTRIC optimization. For QC, gradients were computed at B3LYP-D3(BJ)/ DZVP with the Psi4 [54] program. Our choice of quantum chemical level of theory and basis set was based on benchmarks of quantum chemistry and density functional methods for the accuracy of conformational energies, such as [19, 38, 62]. In these studies it was generally observed that B3LYP-D3(BJ) which includes an empirical dispersion correction [6, 22] is roughly equivalent to MP2 and the wB97X-V functional with nonlocal dispersion [43] in terms of accuracy for conformational energies of small molecules (<30 heavy atoms) in the complete basis set limit, i.e. 0.3-0.4 kcal/mol vs. CCSD(T)/CBS gold standard calculations. Remarkably, when using the DZVP basis set [18] which is equivalent to 6-31G* in size and was optimized for DFT calculations, the accuracy of conformational energy calculations was unchanged compared to the much larger def2-TZVPD and def2-QZVP basis sets [77]. After verification of the published results locally, we chose B3LYP-D3(BJ)/DZVP as the QM method of choice for torsion drives. For molecular mechanics, gradients were run using OpenMM [16] with the OpenFF parsley Force Field (v1.0.0) [60].

### 4.2 Calculating bond orders

#### 4.2.1 AM1 WBO and AM1 ELF10 WBO

To calculate AM1 ELF10 WBO, we used OpenEye’s QUACPAC toolkit [qua] (OpenEye Toolkit version 2019.Apr.2). This implementation provides the ELF10 WBO at the same time as the AM1-BCC charge fitting procedure [33, 34]. For ELF10 WBOs generated in this paper, we used the `get_charges` function in the `chemi.py` module in `fragmenter` versions v0.0.3 and v0.0.4. To calculate AM1 WBO for individual conformers, we used the `OEAssignPartialCharges` with the `OECharges_AM1BCCSym` option from the QUACPAC toolkit for each conformer generated with Omega [ome] (OpenEye version 2019.Apr.2) which is called for the `get_charges` function. For AM1 WBOs calculated to verify the results from the validation set, we generated conformers using the `generate_grid_conformer` function in the `chemi.py` module in `fragmenter` version v0.0.6.

We noticed that AM1 ELF10 WBOs can differ significantly across platforms (Linux vs Mac OS) for triple bonds. All results in this paper were calculated on a Linux machine. The results might be different on a Mac OS.

#### 4.2.2 Wiberg bond order in Psi4

Wiberg-Löwdin bond orders are calculated in Psi4 with the keyword `scf_properties: wiberg_lowdin_indices` using Psi4 version 1.3. All bond orders were computed during the torsion scan computations.

### 4.3 Datasets

#### 4.3.1 Kinase inhibitor dataset

B3LYP-D3(BJ) / DZVP Wiberg-Löwdin bond orders were calculated for 9 kinase inhibitors and Omega generated conformers after a B3LYP-D3P(BJ) / DZVP geometry optimization. The DFT results are available on QCArchive as an `OptimizationDataset` named `Kinase Inhibitors: WBO Distributions`.

The variance of the WBO distributions were calculated using the `numpy` [70] var function version 1.16.2 and their confidence intervals were calculated using arch ‘IIDBootsrap’ function [63] version 4.8.1. To calculate the correlation matrix, we calculated the Pearson correlation coefficient with the `numpy` [70] corrcoef function version 1.16.2. Scripts and data used to generate and analyze this dataset are in https://github.com/choderalab/fragmenter_data/tree/master/wbo-manuscript-figures/kinase_inhibitors_wbos

#### 4.3.2 Subsituted phenyl dataset

The substituted phenyl dataset consists of 3,458 substituted phenyl molecules where the substituents chosen to span a large range of electron donating and withdrawing groups. We arrived at 3,200 molecules by attaching 26 different functional groups to 5 scaffolds (***Figure 5***A) at the X_1_ position, and then attach the 26 functional group (and H) at the X 2 position for a total of 133 molecules per functional group (26 * 5 + 3 (for molecules with H at the X_2_ position)). The AM1 ELF10 WBOs were calculated as described above.

We selected molecules for QC torsion scans as follows:

1. From the 26 functional groups, we only selected molecules from 18 functional groups, skipping X_1_s that either did not have a rotatable bond (fluorine, chlorine, iodine, bromine, nitrile, oxygen), were too congested (triflouromethyl, trimethylamonium) or where the WBOs on the bonds attaching X_1_ to the phenyl ring did not change by more than 0.01 with different chemical group at the X_2_ position (such as methyl).
2. For the 18 functional groups, we chose molecules that were evenly spaced along the WBO range of that functional group, up to 15 molecules. While all the skipped functional groups for X_1_ were allowed to be at X_2_, we did not include the negative oxygen at X_2_ because OpenFF have not yet benchmarked the level of theory to use for anions.
3. After selection, we had 140 molecules that we submitted to QCArchive for both QC and MM torsion scan.

The dataset is available on QCArchive as a `TorsionDriveDataset` named `OpenFF Subsituted Phenyl Set 1`. This dataset also includes the biphenyl torsion scans shown in ***Figure 1***. Scripts used to generate and analyze this dataset can be found in https://github.com/choderalab/fragmenter_data/tree/master/phenyl_benchmark

#### 4.3.3 Interpolating torsion force constants

Given a set of molecules with the common torsion `[*:1]~[#6X3:2]-[#8X2:3]~[*:4]`, such as those given for TIG9 in ***Figure 8***, we can perform a linear fit on the observed torsion barrier height against the calculated AM1 ELF10 Wiberg bond order. The slope and intercept of this fit (Table 2) can then be used to define a torsion parameter. This parameter will give the barrier height for a torsion based on the determined WBO of the torsion’s central bond, and can be applied to molecules outside of this training set that feature the same torsion.

For TIG9, we derived a y-intercept of - 161.3427 kJ/mol and a slope of 193.97 kJ/mol. For a torsion with only a single periodic term, such as that for TIG9, we need only define the torsion height *k*_1_ for bond orders 1 and 2. These *k*_1_ values will define the interpolation and extrapolation line used to obtain *k*_1_ values for WBOs below, above, and between these bond orders. We can define these *k*_1_ values in a single force field parameter shown in ***Figure 10***:

**Figure 10.**
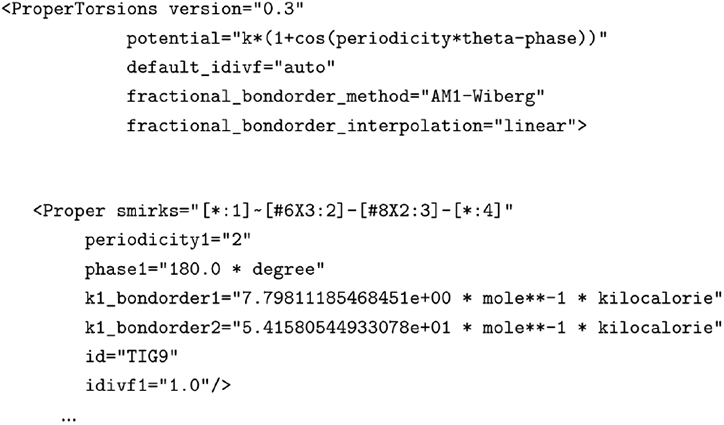
Example parameter file format.

For central bonds that can feature bond orders up to 3, it may make sense to define *k*_1_ for bond orders 1, 2, and 3. Three points defined in the same parameter will give a piecewise-linear interpolation/extrapolation function, in which bond orders 1 and 2 define the barrier heights for bond orders 2 and below and bond orders 2 and 3 define the barrier heights for bond orders 2 and above. It is also possible to define e.g. *k*_1_ for bond orders 2 and 3, but not for bond order 1. For torsions requiring more than one periodic term, such as *k*_1_ and *k*_2_, the same approach as described above for a single *k* applies to each.

Parameterization of molecules via the [OpenFF Toolkit](https://github.com/openforcefield/openff-toolkit) using a force field with the parameter above will apply this interpolation/extrapolation scheme to matching torsions automatically. For more information on the flexibility and limitations of interpolating torsion parameters via SMIRNOFF-based force fields, see the [SMIRNOFF specification](https://github.com/openforcefield/openff-toolkit/blob/master/The-SMIRNOFF-force-field-format.md).

## 5 Code and Data Availability

Scripts to generate the figures in this paper can be found at http://github.com/choderalab/fragmenter_data/wbo-manuscript-figures

## 6 Acknowledgments

CDS would like to acknowledge Josh Fass, Ana Silveira, Rafal Wiewiora, Caitlin Bannan, Alberta Gobbi, Geoff Skillman, and members of the Chodera lab for helpful discussion.

## 9 Author Contributions

Conceptualization: CDS, CIB, JDC

Methodology: CDS, DGAS, JM

Software: CDS, DGAS, DD

Formal Analysis: CDS, JM

Investigation: CDS, JM

Writing – Original Draft: CDS, JM

Writing – Review & Editing: CDS, DM, JDC

Funding Acquisition: CDS, JDC

Resources: JDC, DM

Supervision: JDC, DM

## Author Information

### ORCID

Lee-Ping Wang: 0000-0003-3072-9946

## Appendix A Supplemental Information

### A.1 The Mayer bond order (MBO) extends the Wiberg bond order (WBO) to non-minimal basis sets

The WBO was developed within Pople’s “complete neglect of differential overlap” (CNDO)formalism [59, 79] and is defined as:

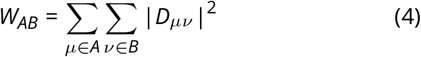

where the quadradic sum of the electron density *D* is over occupied orbitals *μ* and *v* in atoms A and B in the bond. At higher levels of QM theory where non-minimal basis sets are used, *W_AB_* would give values that increase in size with the basis sets and need to normalized for physical quantities. In Psi4 [54], The Löwdin normalization scheme [41] is used to calculate the WBOs. In this scheme, the non-minimal basis set *ϕ* are transformed by a linear transformation 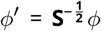 Where **S** is the the overlap matrix.

Mayer [45] incorporates **S** directly in the bond order calculation:

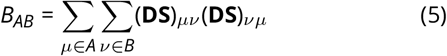

where **D** is the density matrix for occupied orbitals for closed shell systems, and **S** is the overlap matrix. This equation reduces to the WBO when minimal basis sets are used.

### A.2 Comparing MBO and WBO at different levels of QC theory

Psi4 [54] implements both Wiberg-Löwdin and Mayer bond orders. Here, we compare WBO and MBOs at different levels of theory.

#### A.2.1 Differences in conformation-dependent bond order distributions between WBO and MBO at different levels of QC theory

Both the WBO and MBO are functions of electron density, which is conformation dependent. As we have shown in ***Figure 2***, conjugated bonds have higher variance than non-conjugated bonds. To examine both the variance in the quality of individual QC methods electronic densities and WBOs and MBOs effect on bond order values and conformation-dependent variance, we compared WBOs and MBOs calculated from wavefunctions using different QC methods. In ***Figure 1*** we compare AM1 [15] WBOs, HF-3c [69] MBOs and WBOs, and B3LYP(BJ)/DZVP [5, 18, 21, 22] MBOs and WBOs. Figure 1a shows gefitinib, a kinase inhibitors, with its rotatable bonds highlighted to correspond to the colors of the distributions shown in the other panels of ***Figure 1***. The bond order distributions shown in ***Figure 1*** were calculated over 232 Omega [28] generated conformers of Gefitinib, the resulting conformers were optimized at the same level of theory as the subsequent bond order computations.

For AM1, the MBO and WBO methods are equivalent and only Wiberg values were presented. AM1 WBO distributions have lower variance than B3LYP for the non-conjugated bonds, but have significantly higher variance and more modes for the more conjugated bonds.

WBO and MBO distributions calculated from HF-3c optimized structures (Figure 1c and 1d) had lowest variances than other QC methods. In addition, both MBO and WBOs have similar values and shapes of their distributions. This might be a result of HF-3c using a small, Gaussian atomic orbital (AO) basis set. As discussed in the previous section, the MBO reduces to the WBO when orthonormalized basis AO are used. In HF-3c, a small, Gaussian basis set is used which might be the reason why the MBOs and WBOs are so similar.

When using larger basis sets (B3LYP(BJ)/DZVP), the Mayer and Wiberg bond orders differ significantly. First, MBOs have significantly higher variance with respect to conformation. In addition, WBO values are significantly higher than Mayer and AM1 and HF-3c WBOs. The basis sets are Löwdin normalized, but the values still seem to grow with larger basis sets.

#### A.2.2 Differences in conformation-dependent bond order correlation structures between WBO and MBO at different levels of QC theory

As we have shown in Figure 2, conjugated systems have strongly correlated and anti-correlated bond orders with respect to conformations. Figure 2 compares such correlation maps and their resulting structures for the bond orders calculated in Figure 1.

Correlation plots from HF-3c MBOs and WBOs (Figure 2c and 2d) have structures that are very similar to each other. This result is similar to Figure 1c and Figure 1d where the distributions were also very similar. However, the correlation structures of MBOs and WBOs calculated at B3LYP(BJ)/DZVP are different from each other. MBOs show stronger correlations and anti-correlations for local conjugated systems. For example, the bonds in the phenyl ring (alternating correlated and anti-correlated square formed by bonds 24-29) are all alternatively strongly correlated and anti-correlated. In addition, bonds 19–23 and 23–24 are strongly correlated with each other and the quinazoline heterocycle. The WBOs calculated both at B3LYP(BJ)/DZVP and AM1 still show alternating correlated and anti-correlated bond orders for the phenyl ring, albeit not as strongly as seen for MBOs. However, there is also a diffusion of the coupling to bonds outside of the phenyl ring. Bonds 24–25 and 24–29 are also coupled with quinazoline the bonds between the qunazoline and the phenyl (19–23 and 23–24).

**Appendix A Figure 1.**
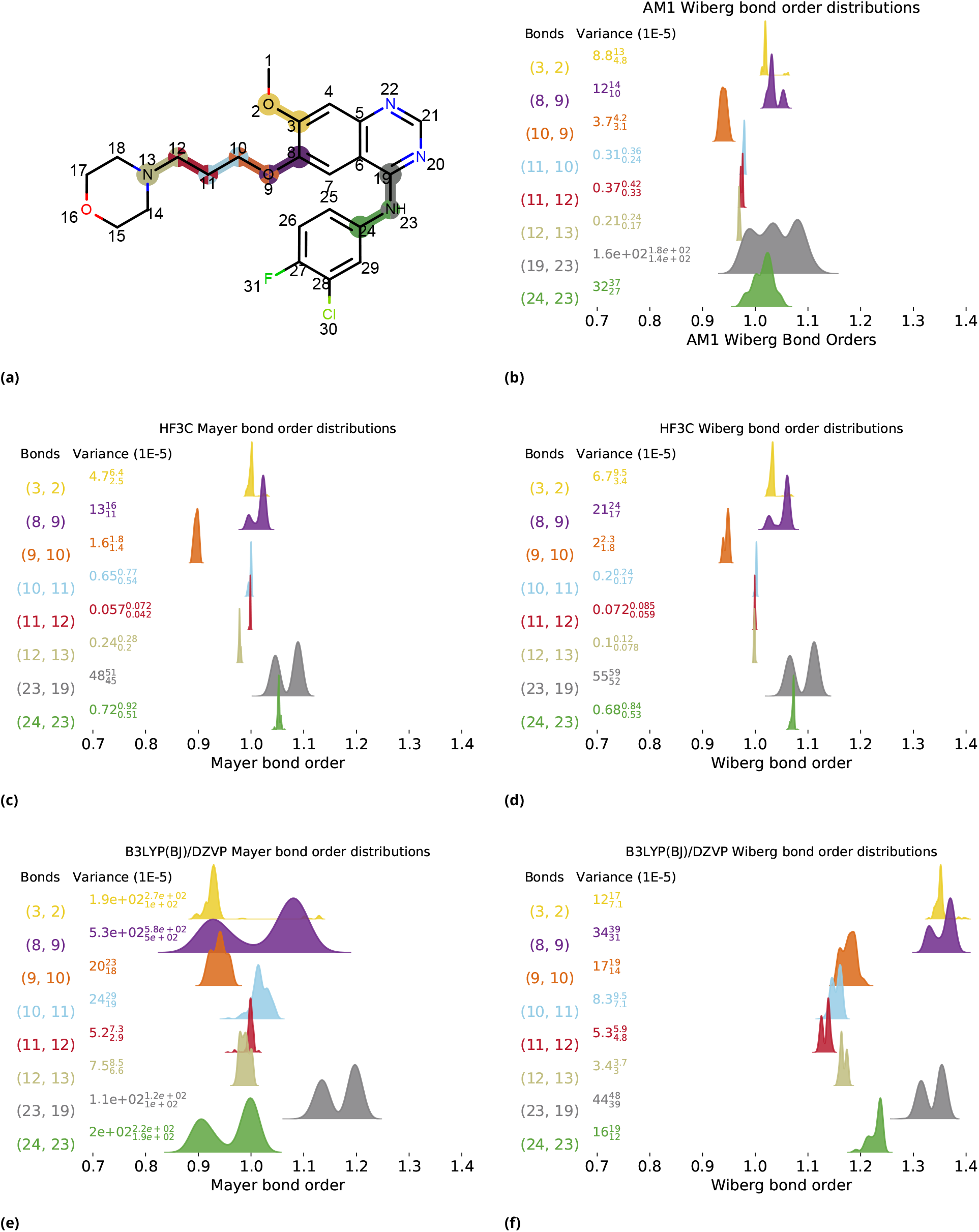
Bond order distributions for rotatable bonds over 262 conformers for Mayer and Wiberg at different levels of QM theory. (a) gefitinb, with atoms numbered highlighted to correspond to numbers and colors on distribution plots; (b) Distribution of AM1 WBOs for rotatable bonds; (c) Same as 1b but for MBOs at HF3C; (d) Same as 1b calculated at HF3C; (e) Same as 1b but for MBOs at calculated at B3LYP; (f) Same as 1b but calculated at B3LYP;

**Appendix A Figure 2.**
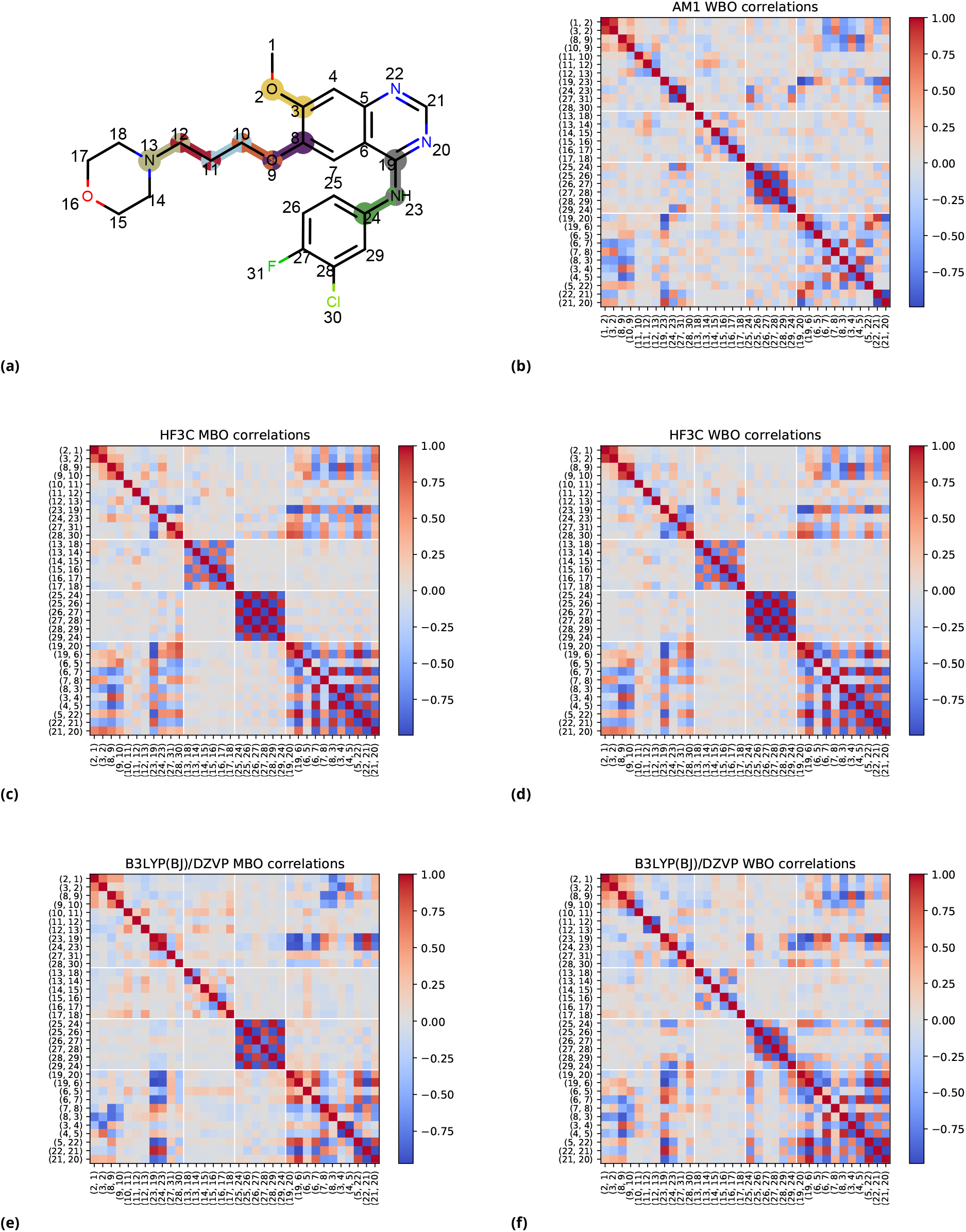
Bond order correlations with respect to conformations for Mayer and Wiberg at different levels of QM theory. (a) Gefitinb, with atoms numbered to correspond to the numbers on the correlation plots; (b) Pearson correlation coefficient of AM1 WBOs for every bond against every bond over 232 conformations; (c) Same as 2b but for MBOs at HF3C; (d) Same as 2b calculated at HF3C; (e) Same as 2b but for MBOs at calculated at B3LYP; (f) Same as 2b but calculated at B3LYP;

From these results, it is not clear which fractional bond order scheme is better at capturing electronic coupling in larger conjugated systems. Further investigation is needed, specifically for deciding if MBO or WBO would be better at differentiating steric and resonance contributions to QC torsion scans discussed in Section 2.2.9. It is encouraging to see that both AM1 and B3LYP(BJ)/DZVP WBOs generate such similar correlation structures.

**Appendix A Figure 3.**
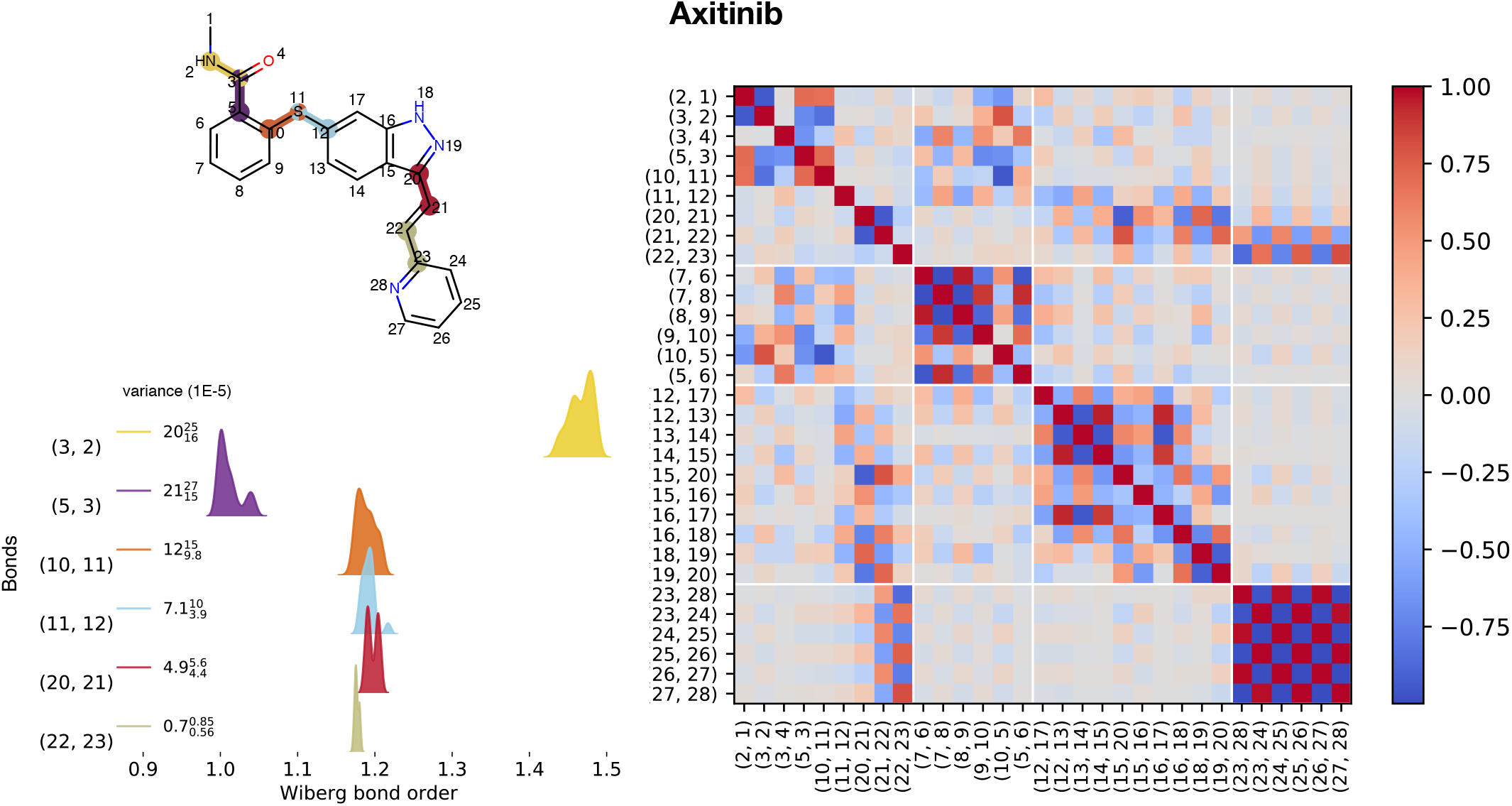
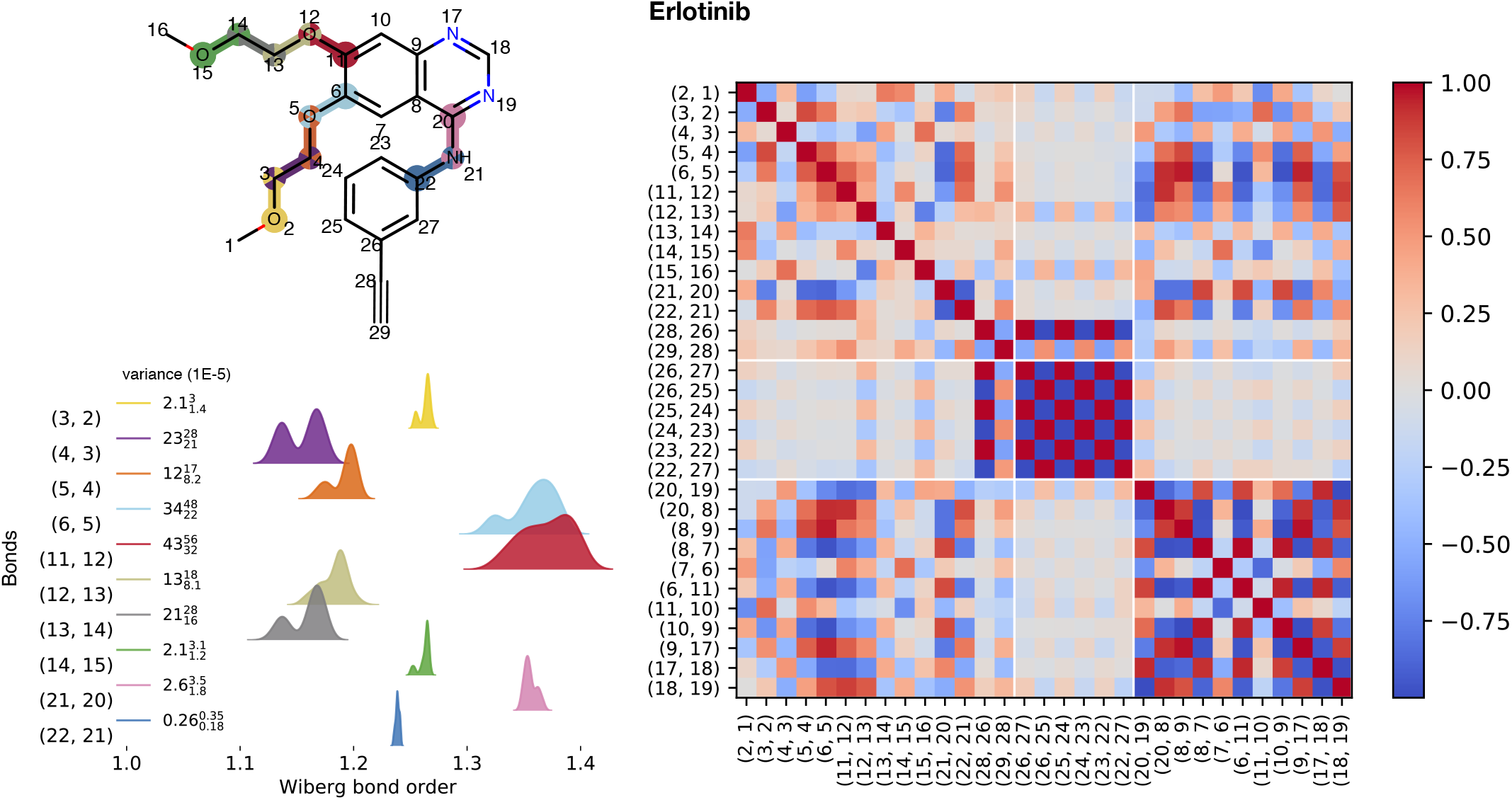
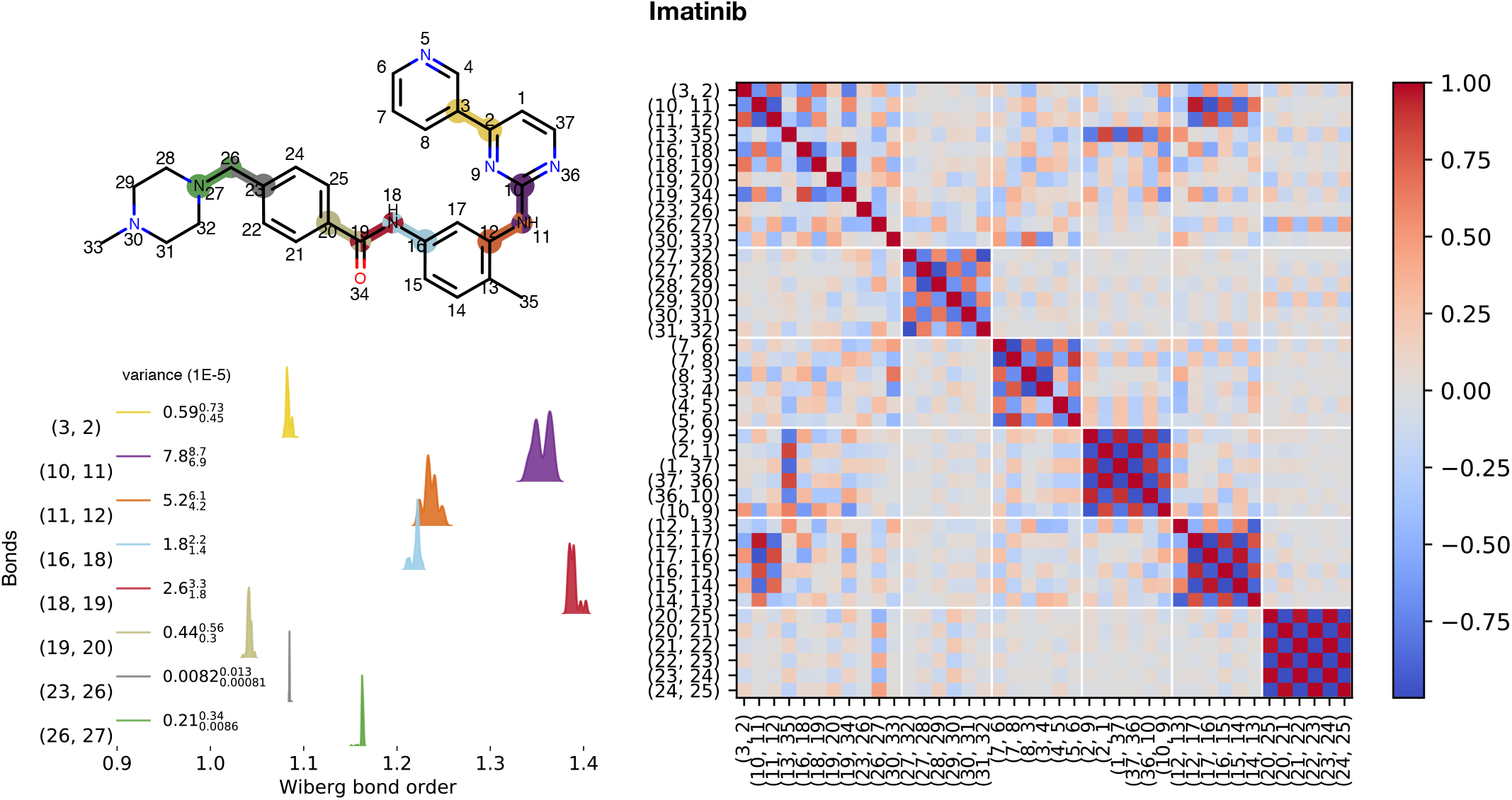
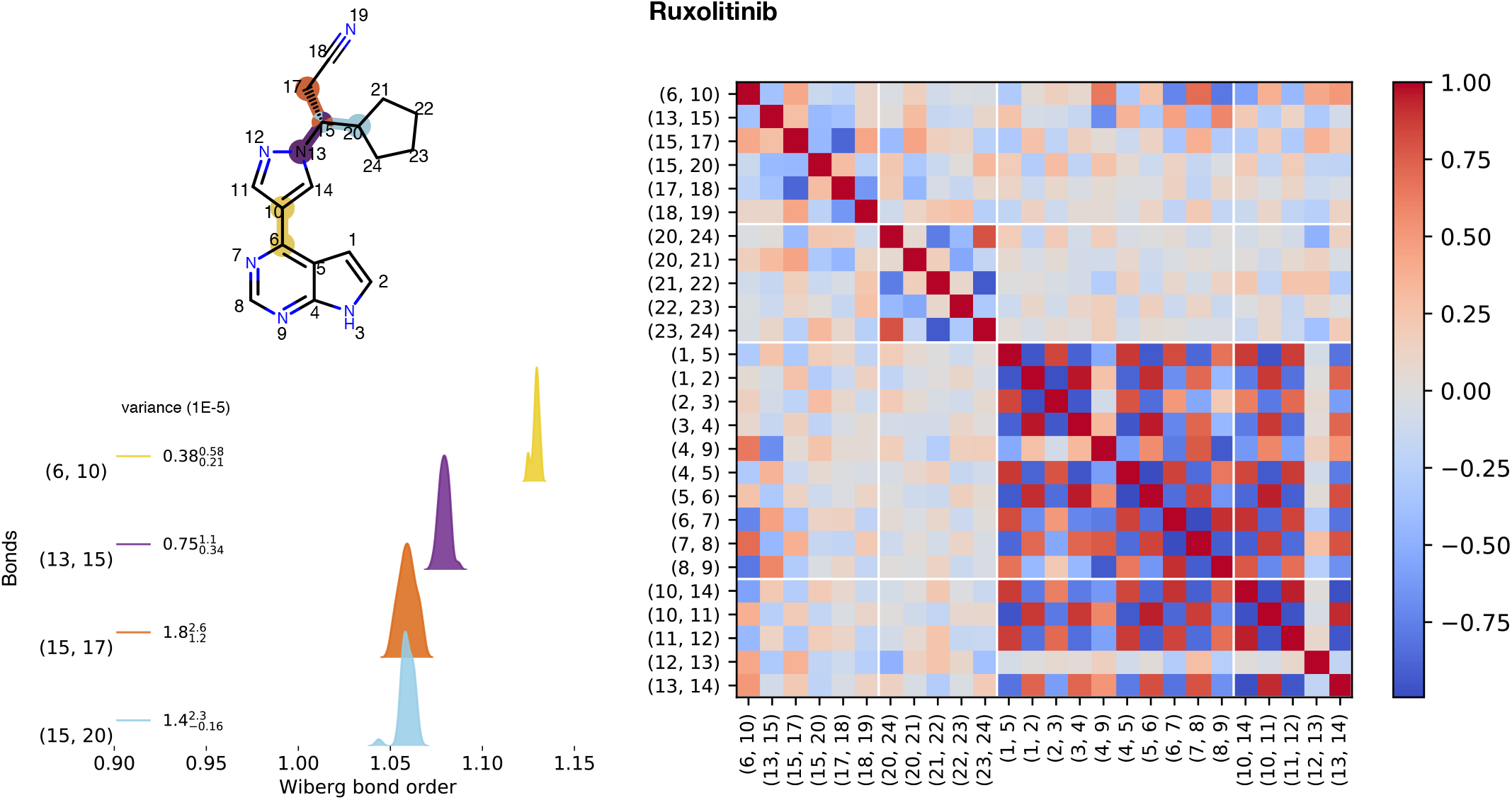
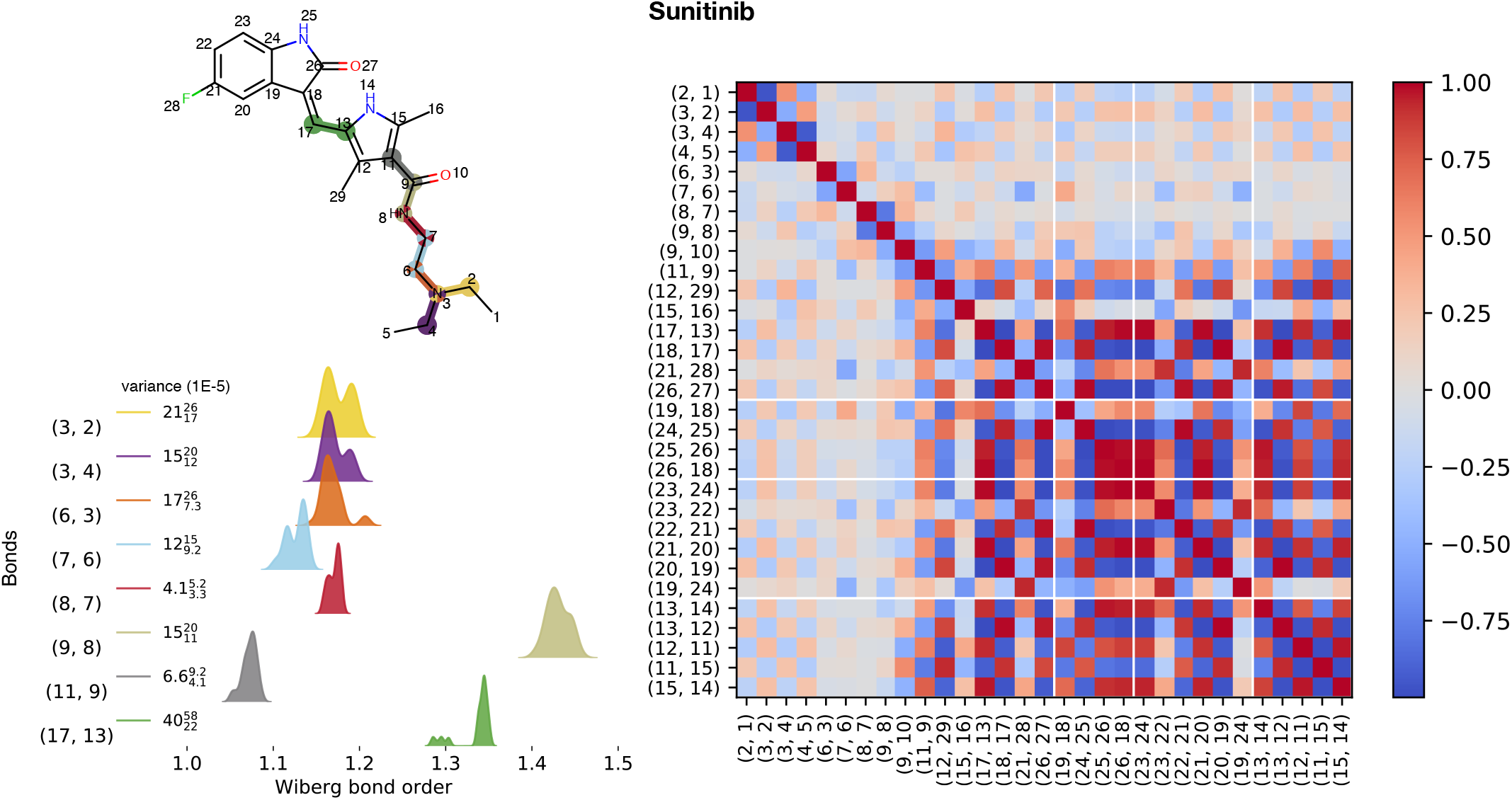
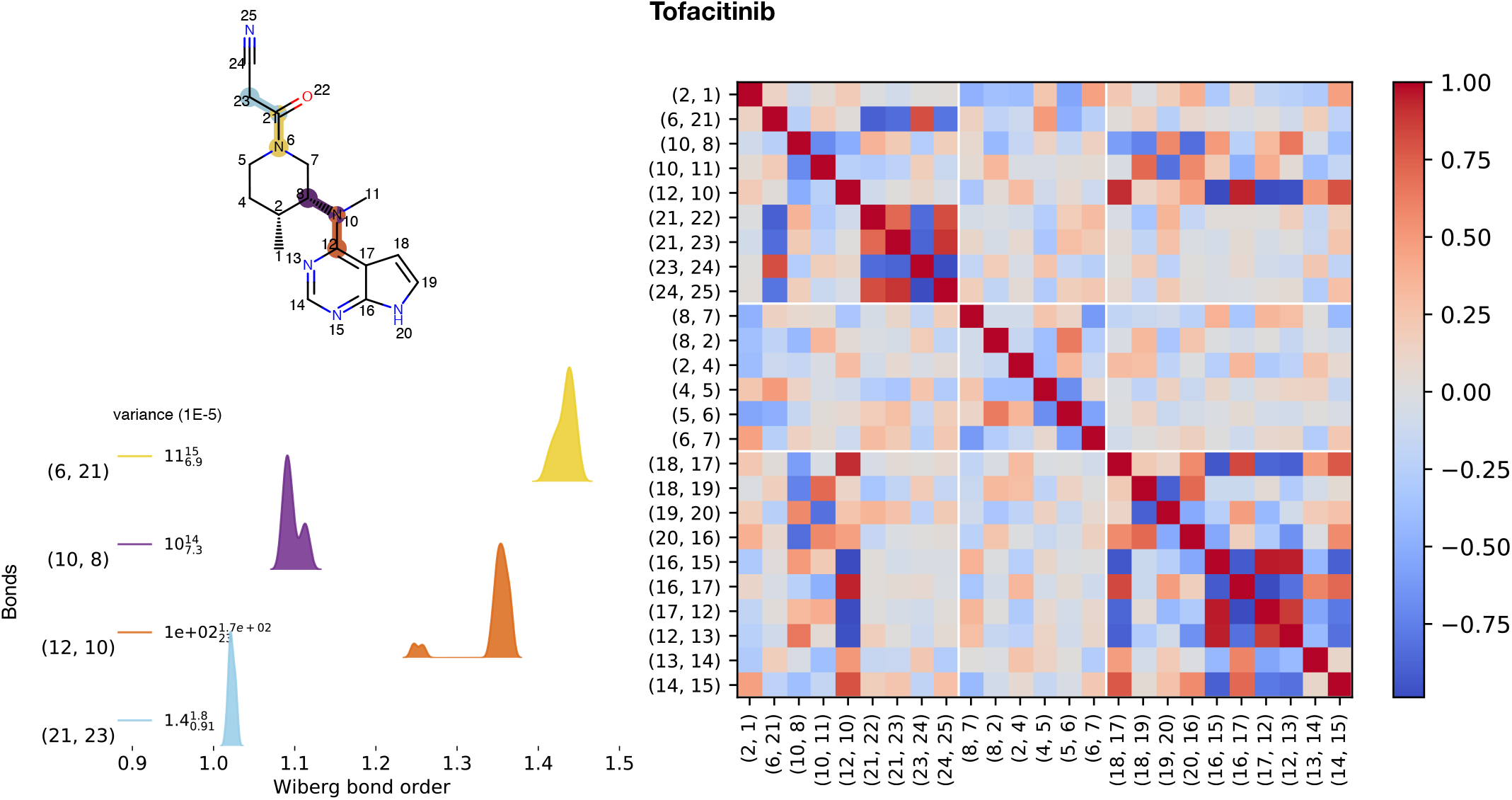
Variance and correlations of WBO distributions with respect to conformations for a set of kinase inhibitors. This figure shows WBO distributions and correlations for a set of kinase inhibitors calculated at B3LYP-D3(BJ) / DZVP. Optimized conformations and their WBOs are on QCArchive (`OptimizationDataset`, named `Kinase Inhibitors: WBO Distributions`)

**Appendix A Figure 4.**
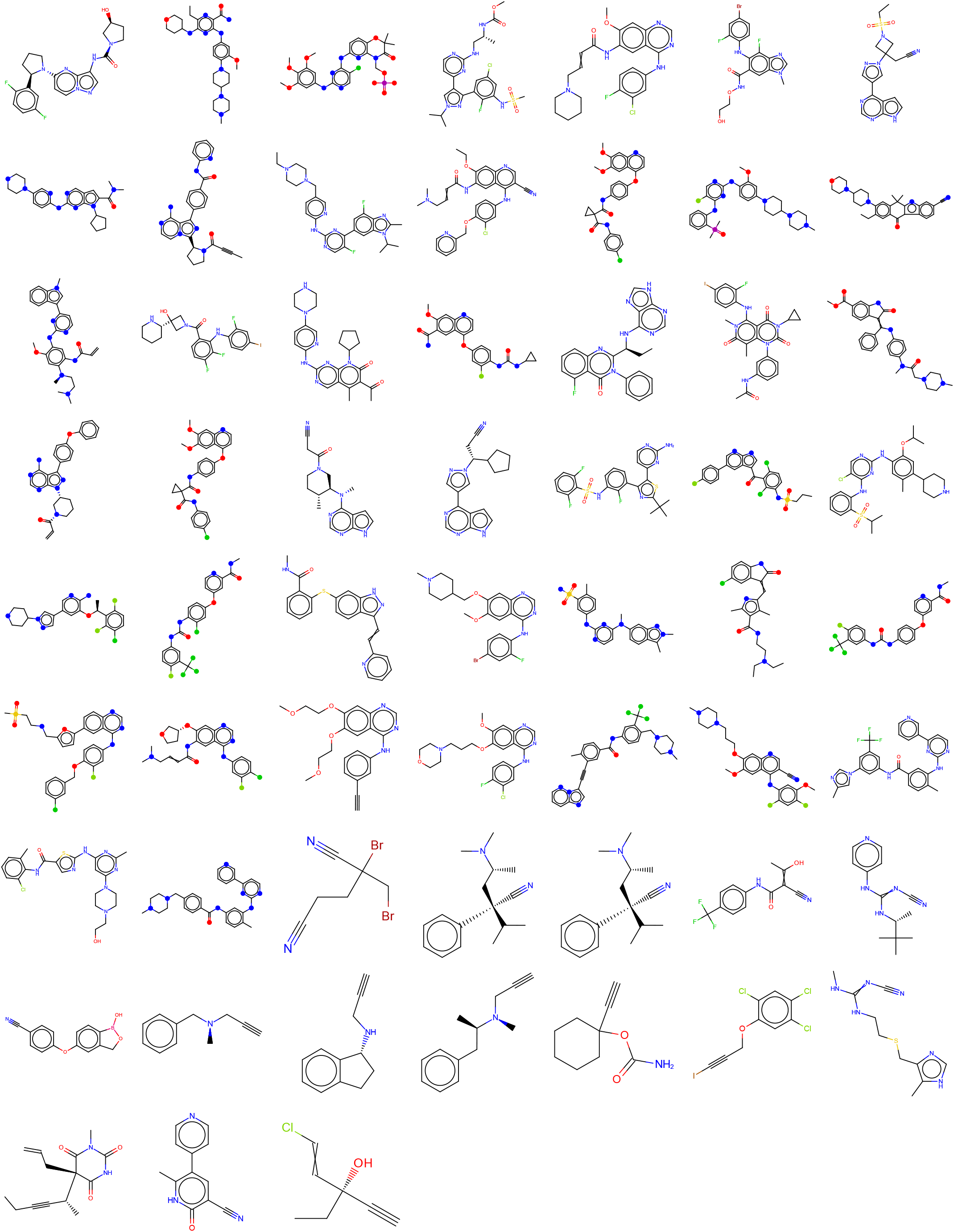
Druglike molecules used to calculate ELF10 AM1 WBOs. This set of molecules were selected to cover bonds of different types and multiplicities. The x axis is the ELF10 WBO for the bond attaching the functional group to the benzoic acid. The Y axis for the left and right panel are *σ_m_* and *σ_p_*, the Hammet meta and para substituent constant.

**Appendix A Figure 5.**
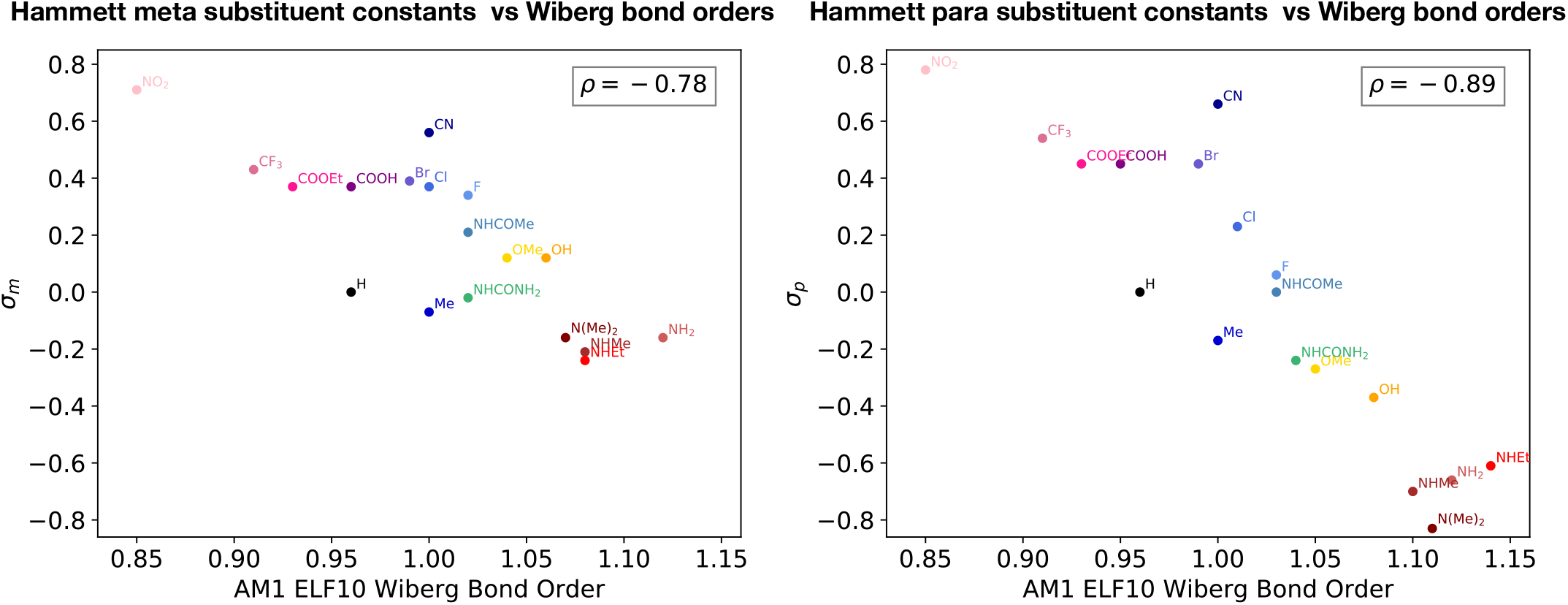
Hammett parameters are anti-correlated with ELF10 WBOs. **[A]** Hammett sigma meta parameters vs AM1 ELF10 WBOs of *X*_1_ meta to carboxylic acid. **[B]** Same as **A** but for para substituents.

**Appendix A Figure 6.**
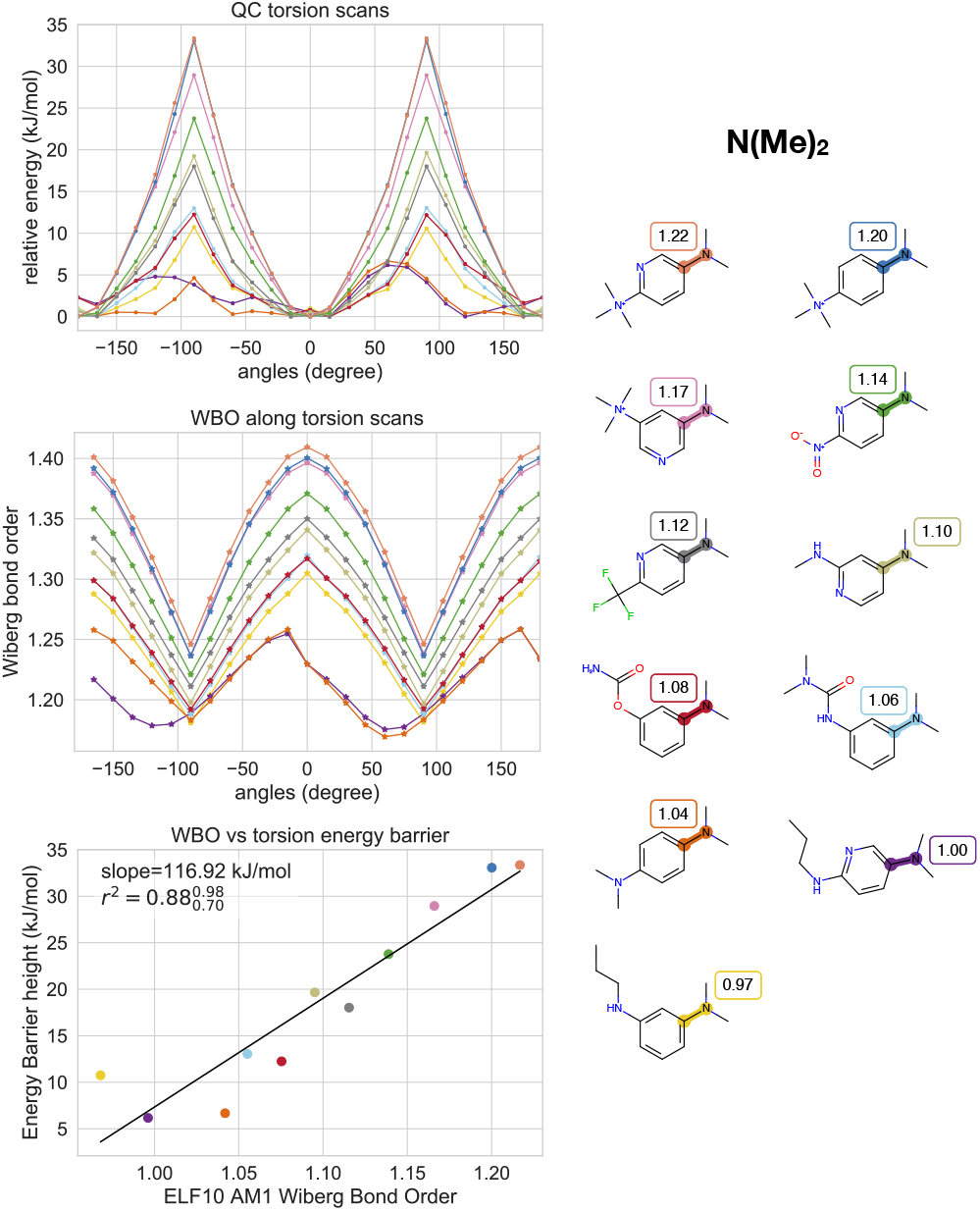
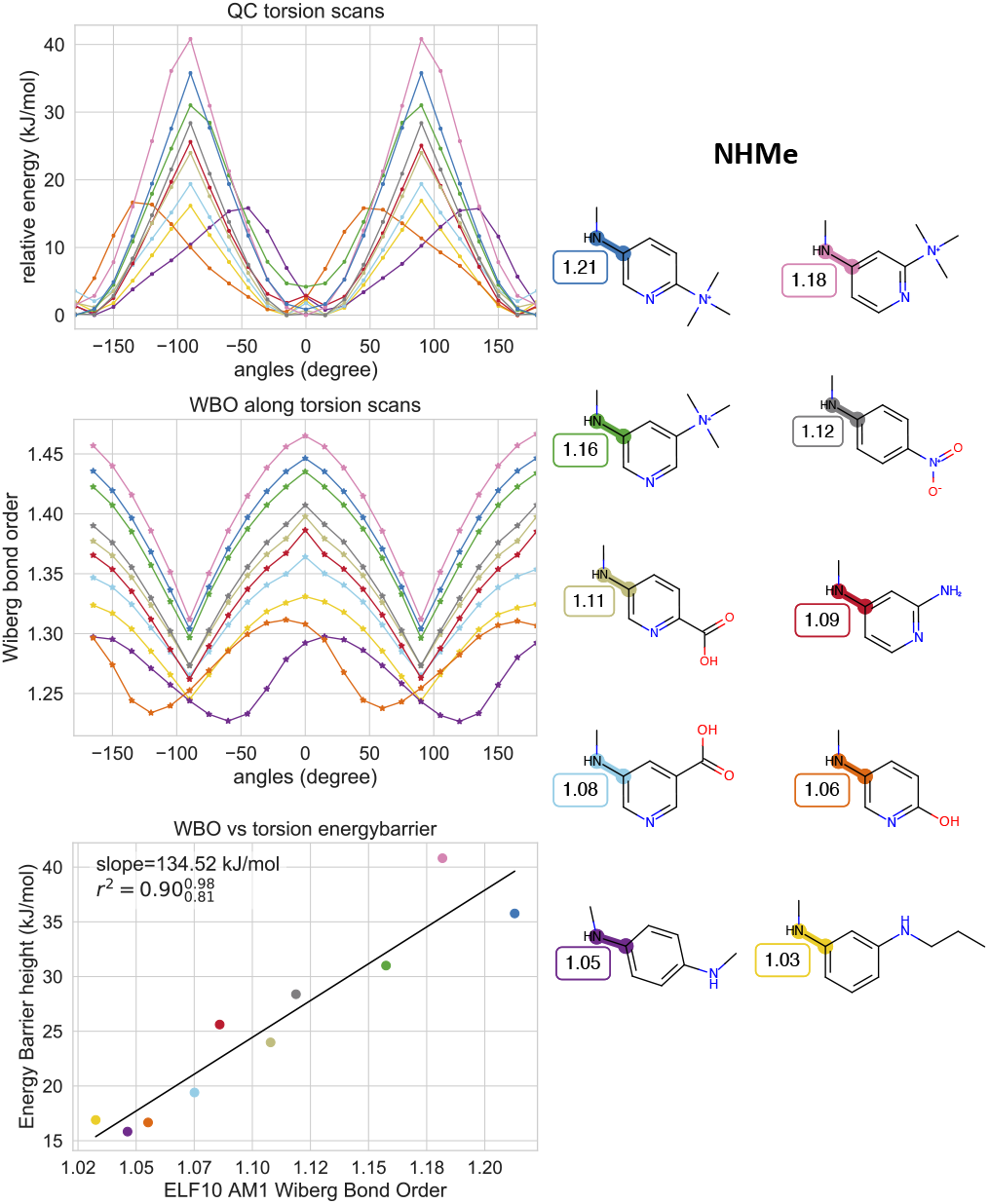
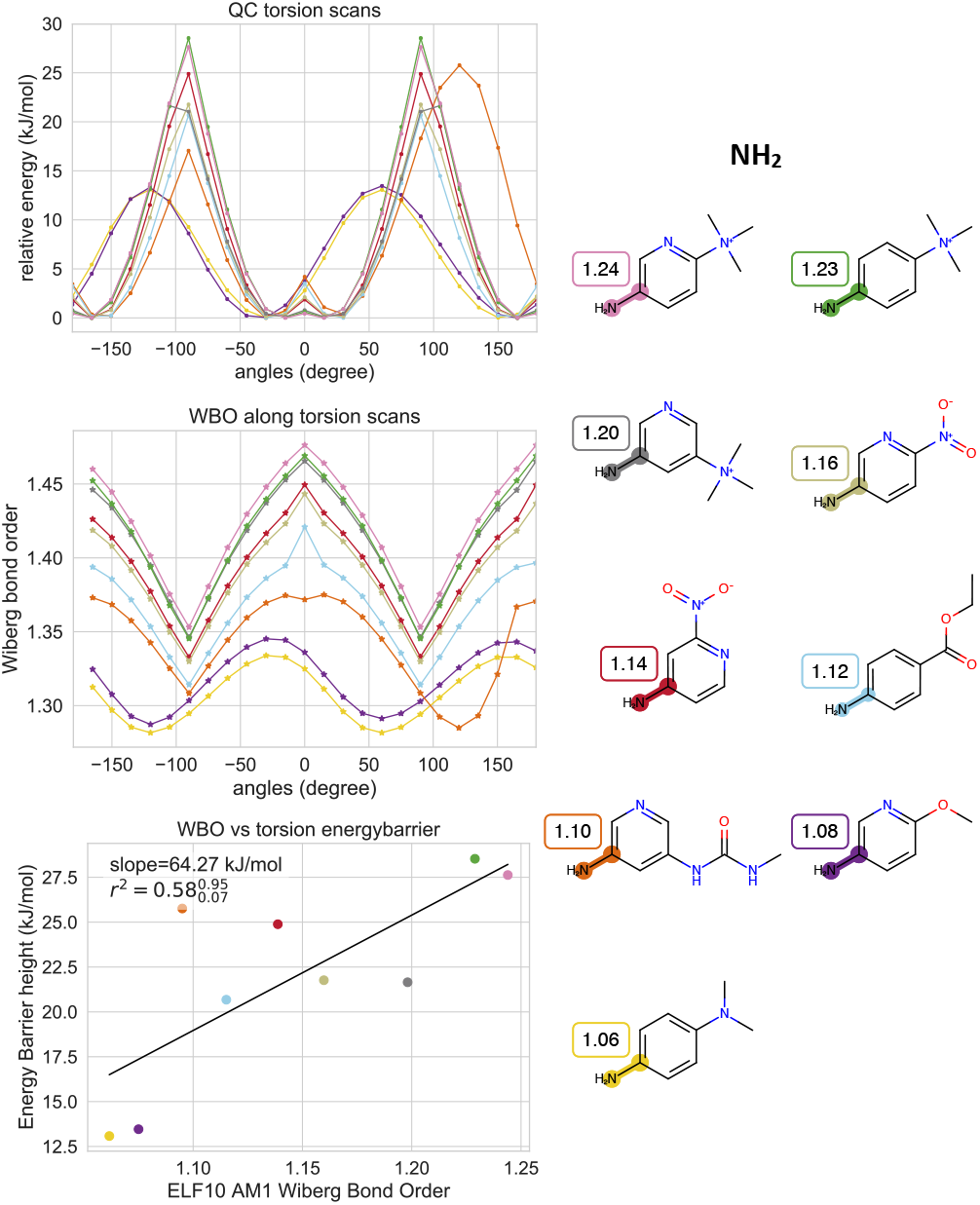
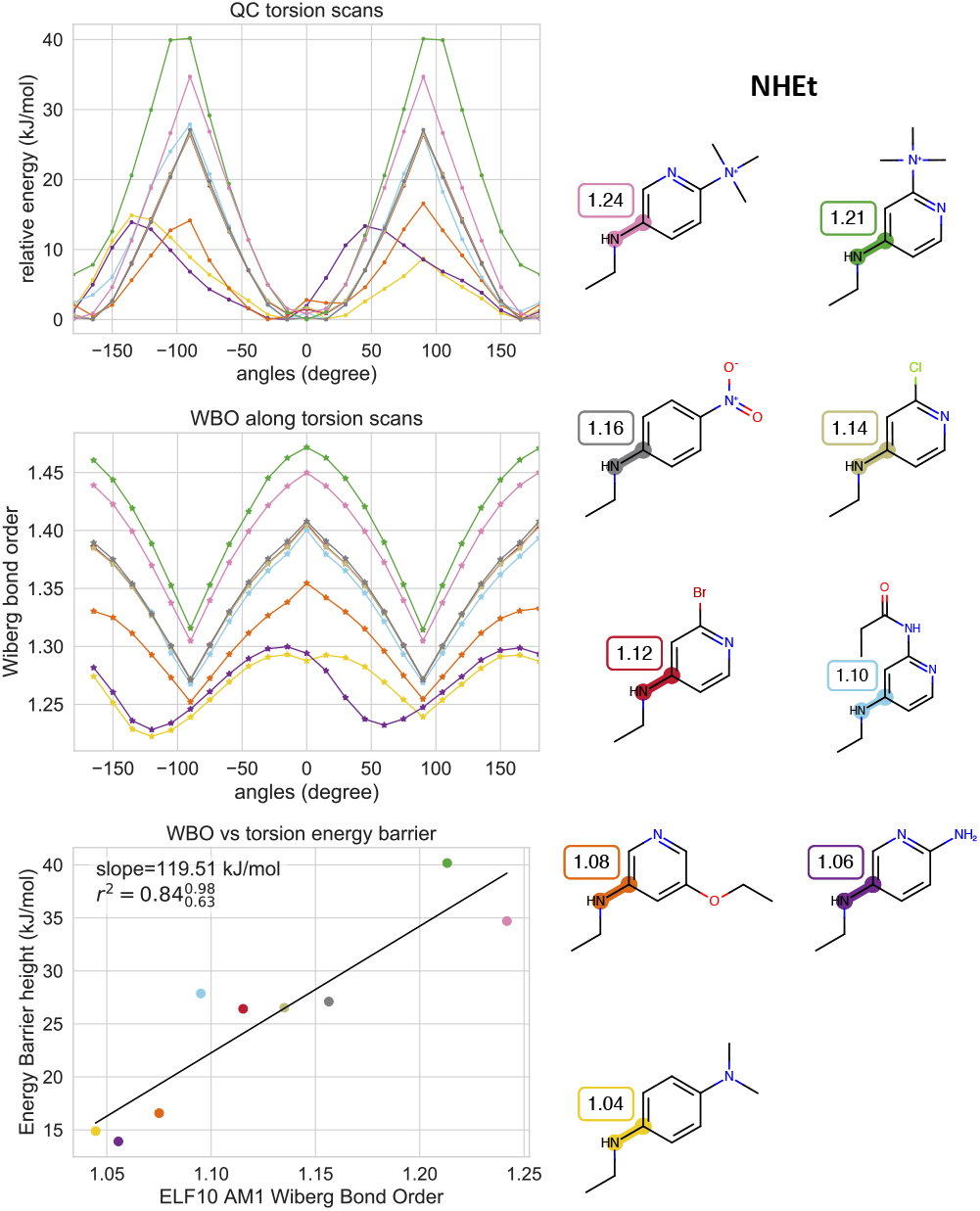
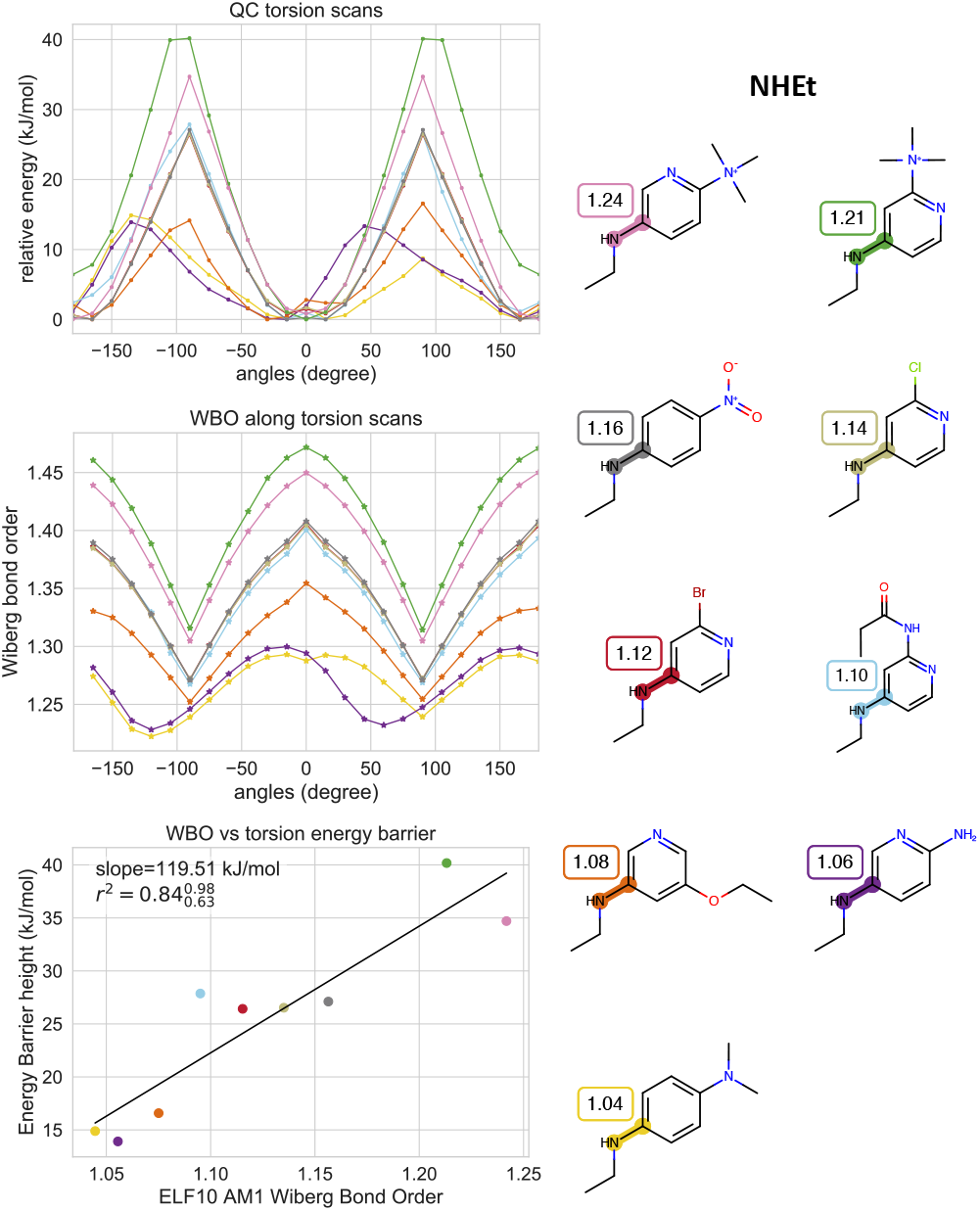
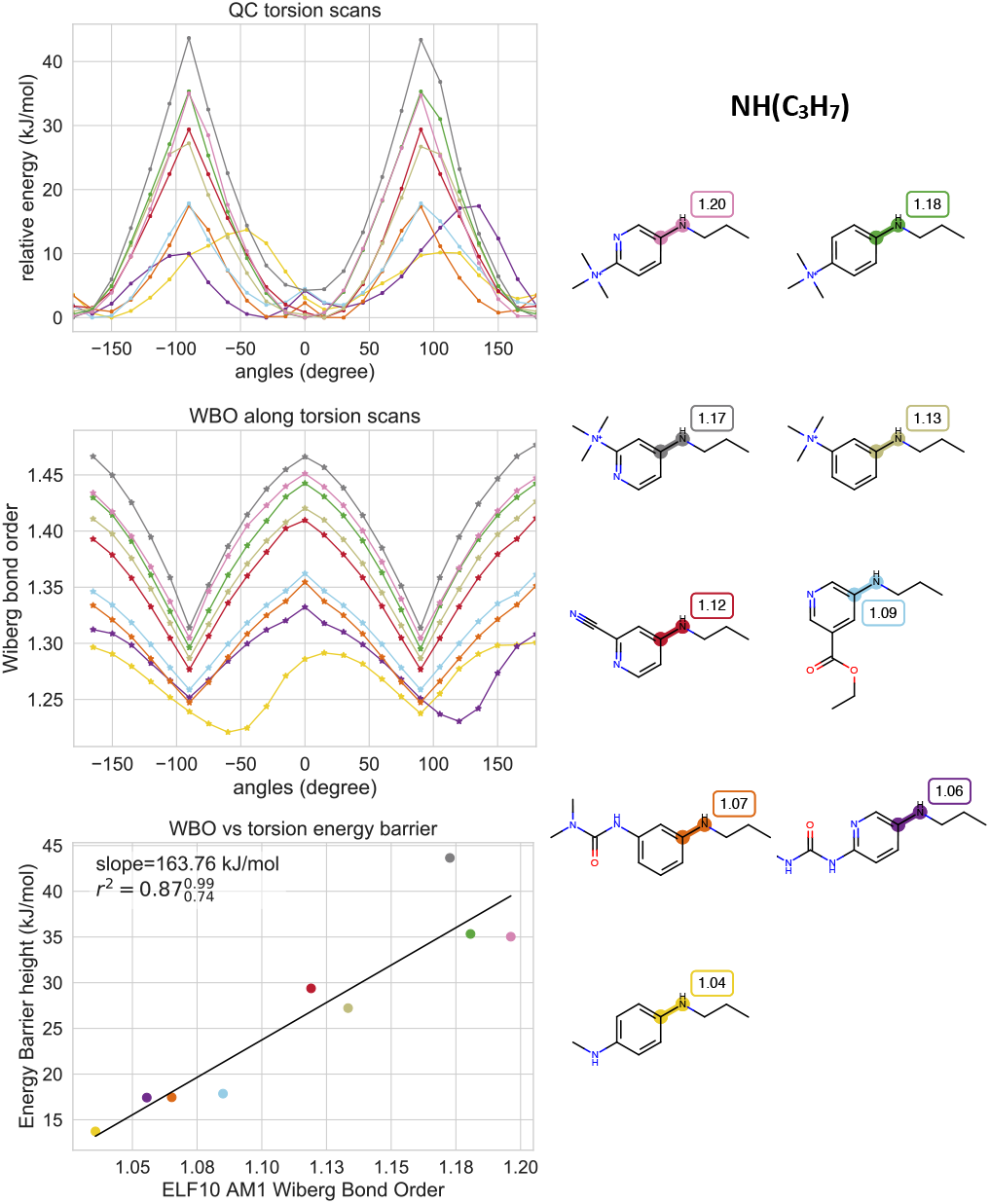
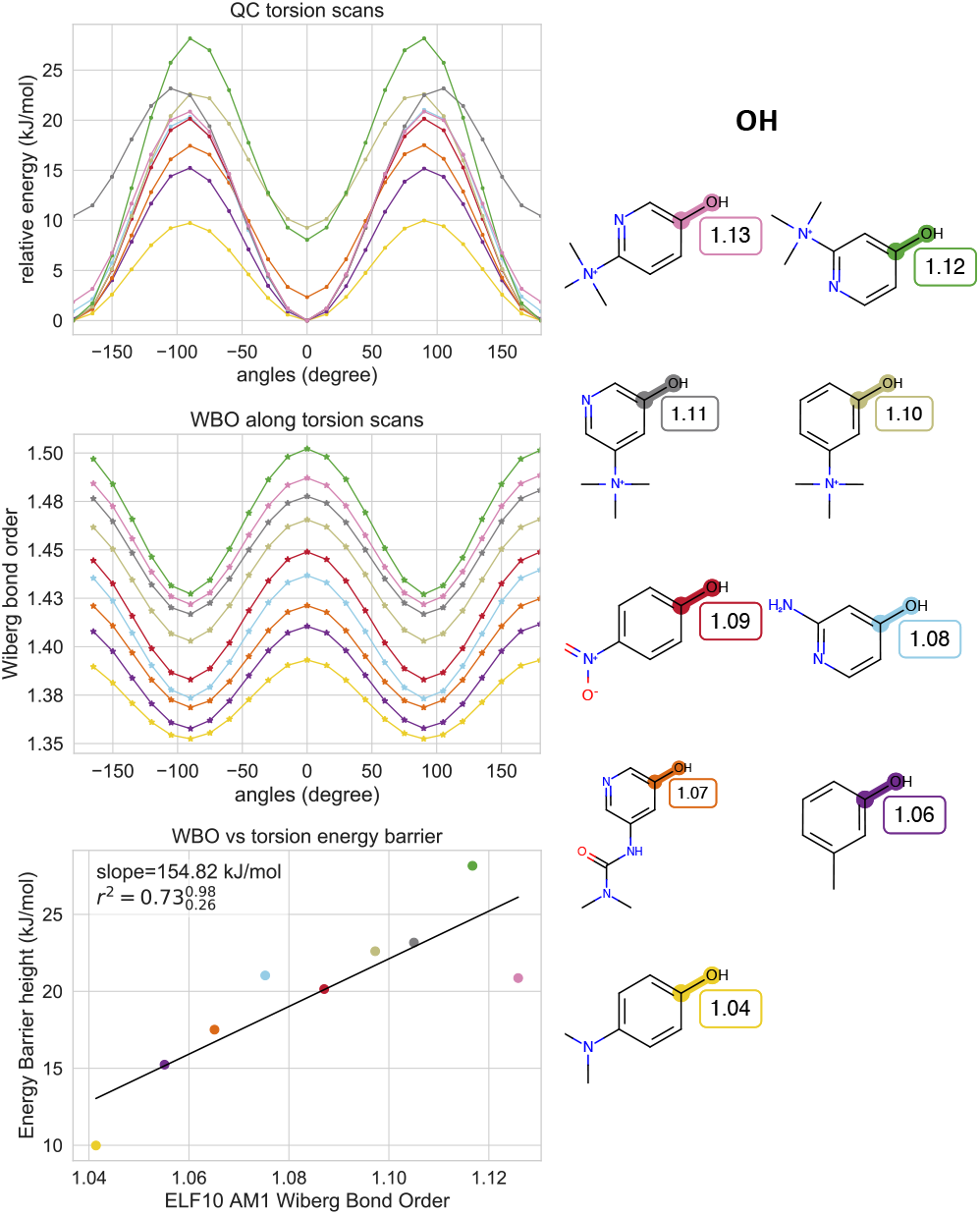
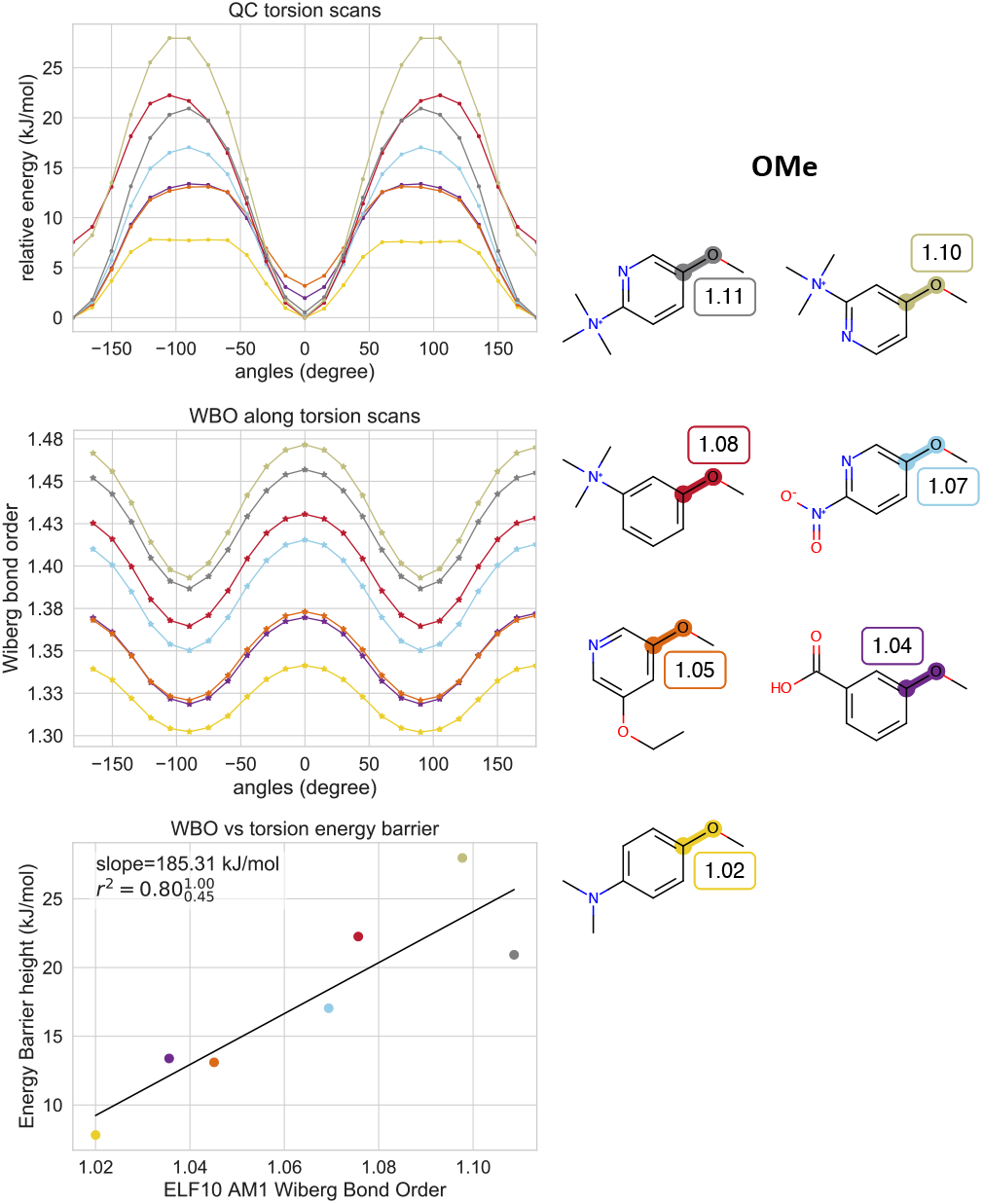
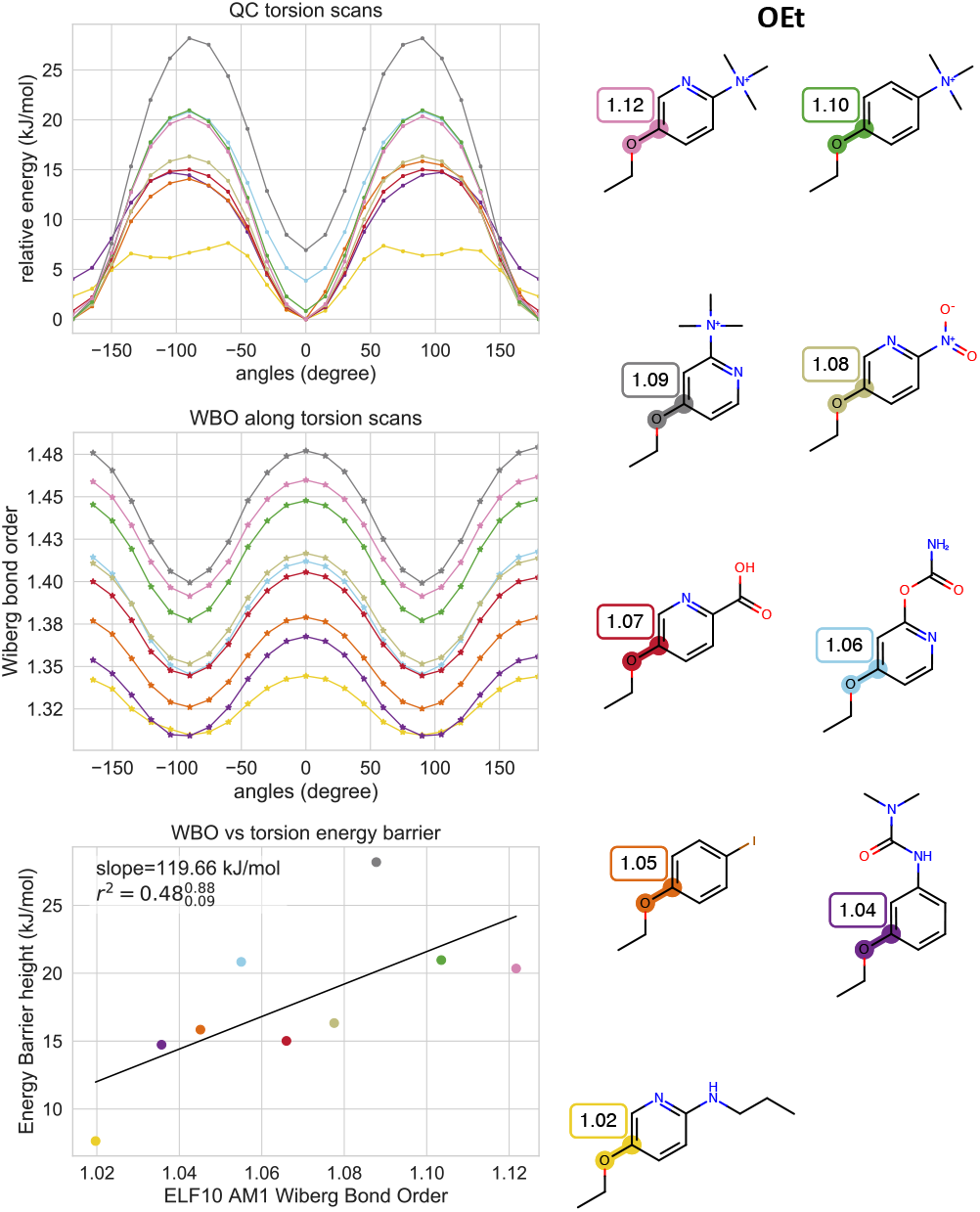
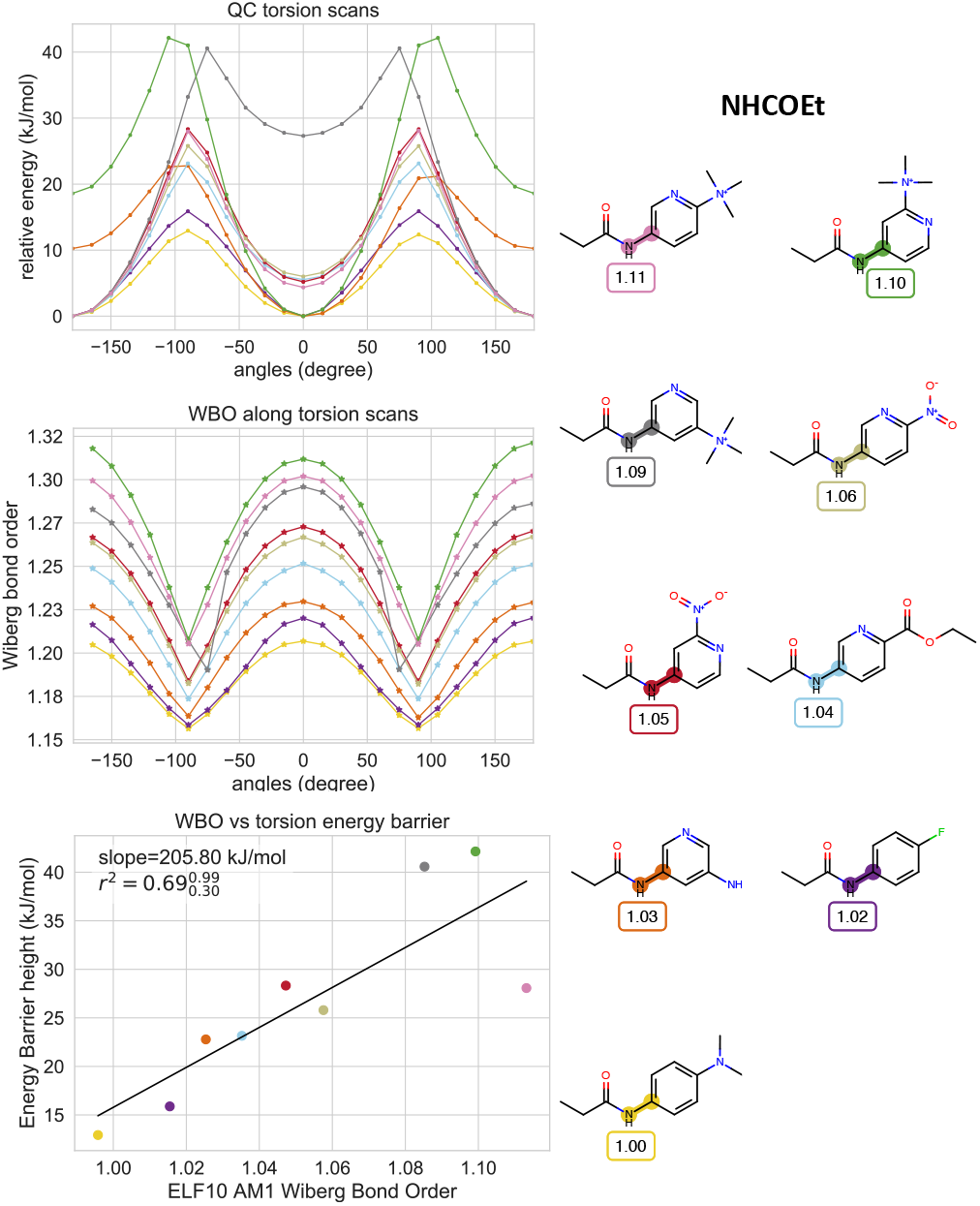
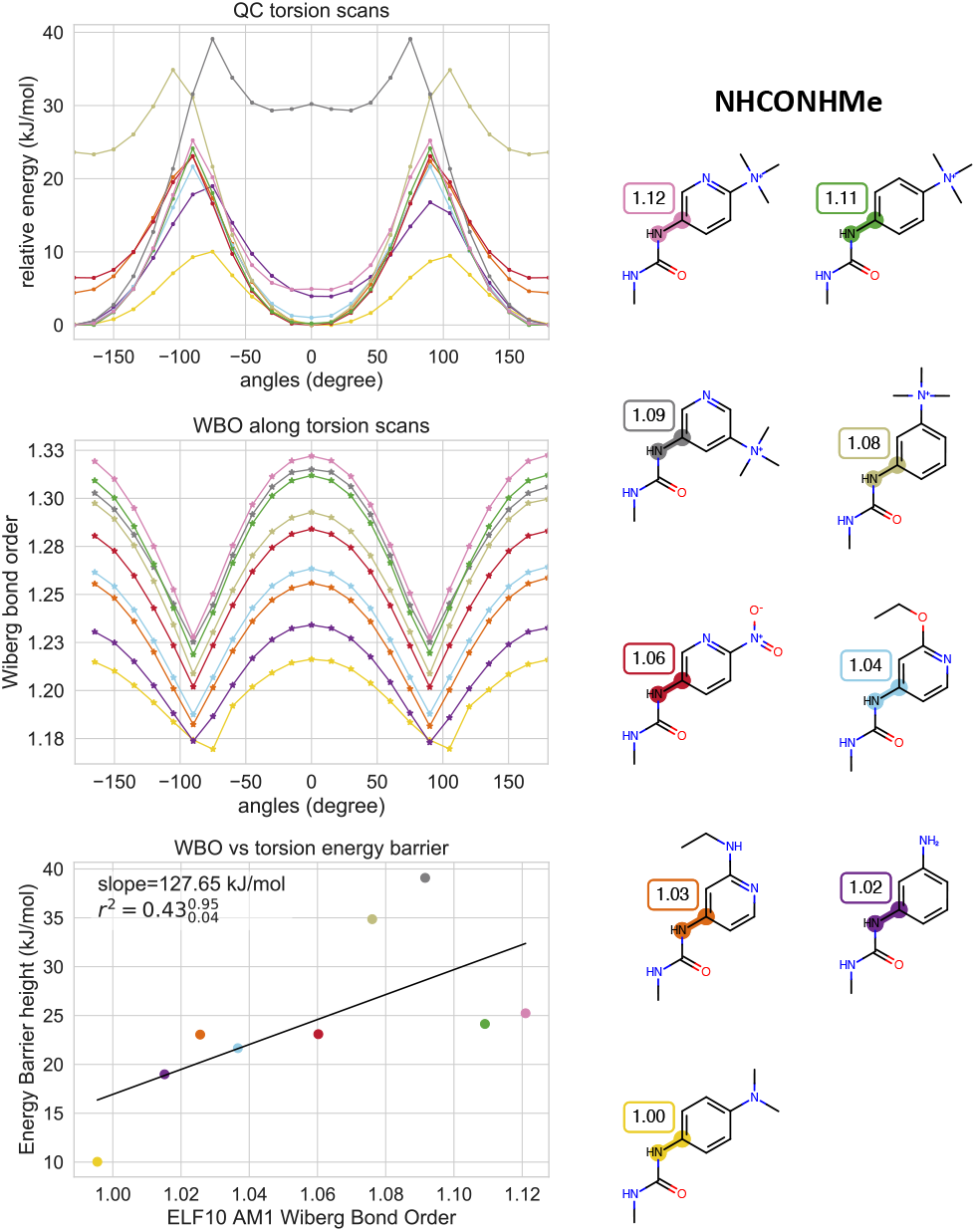
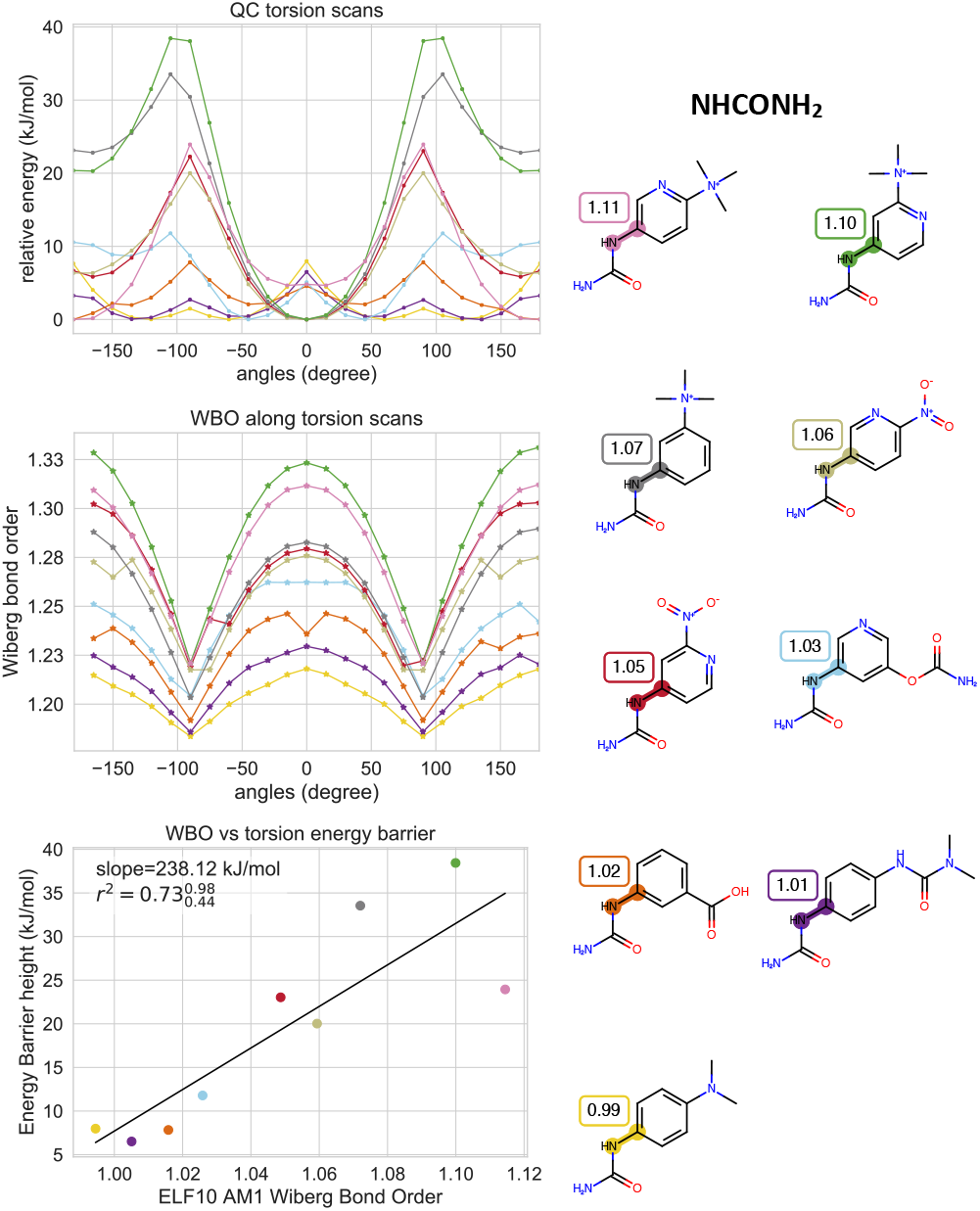
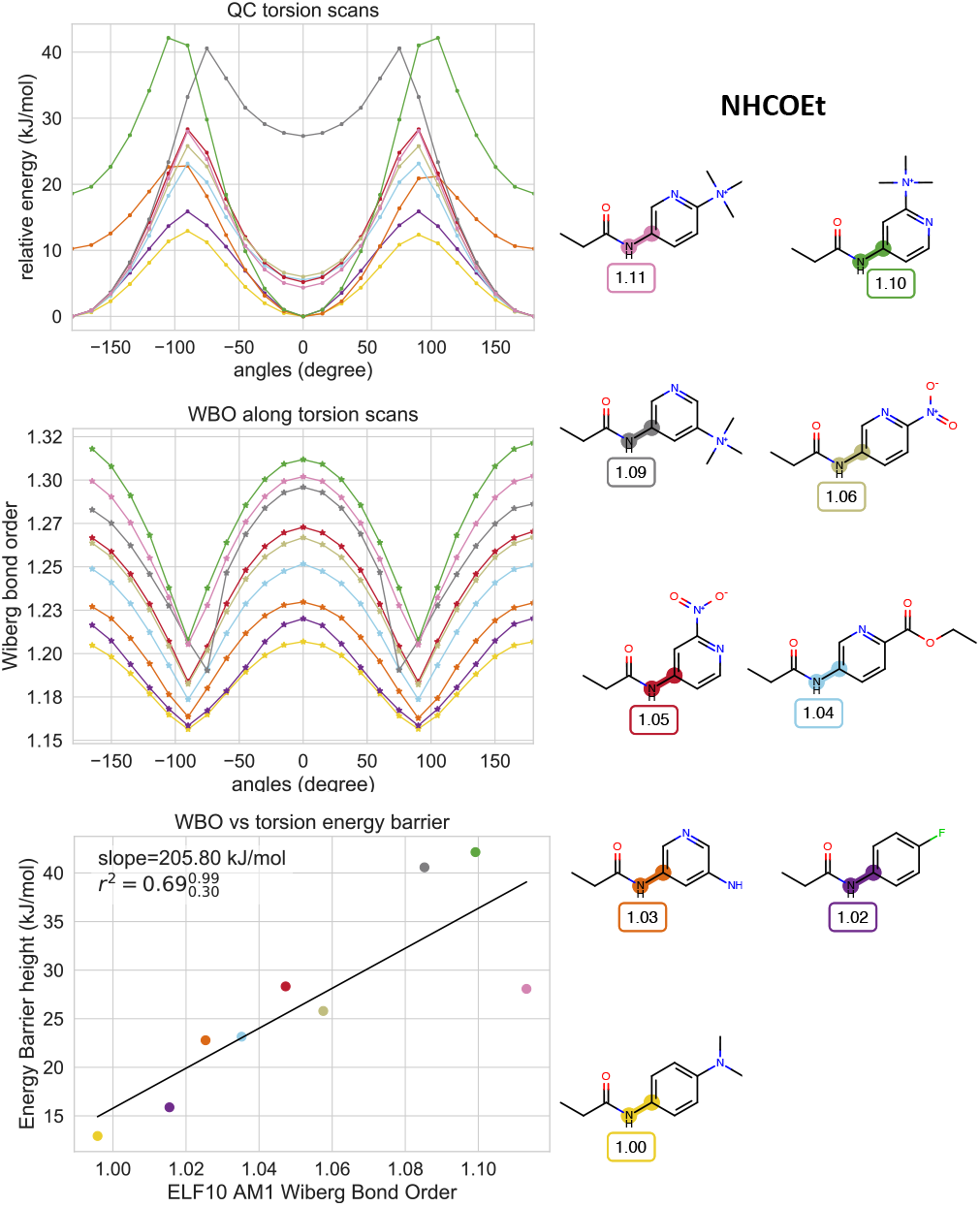
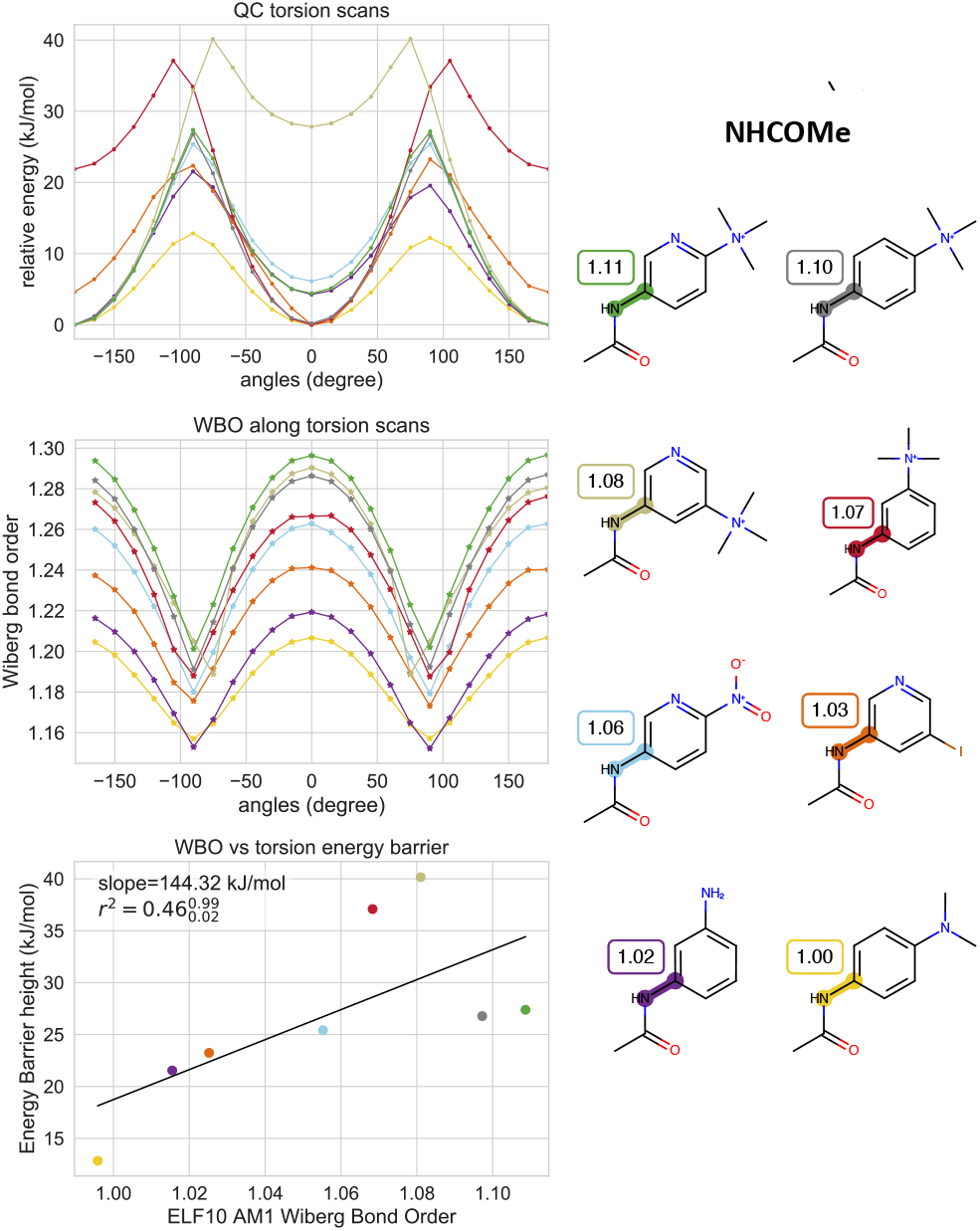
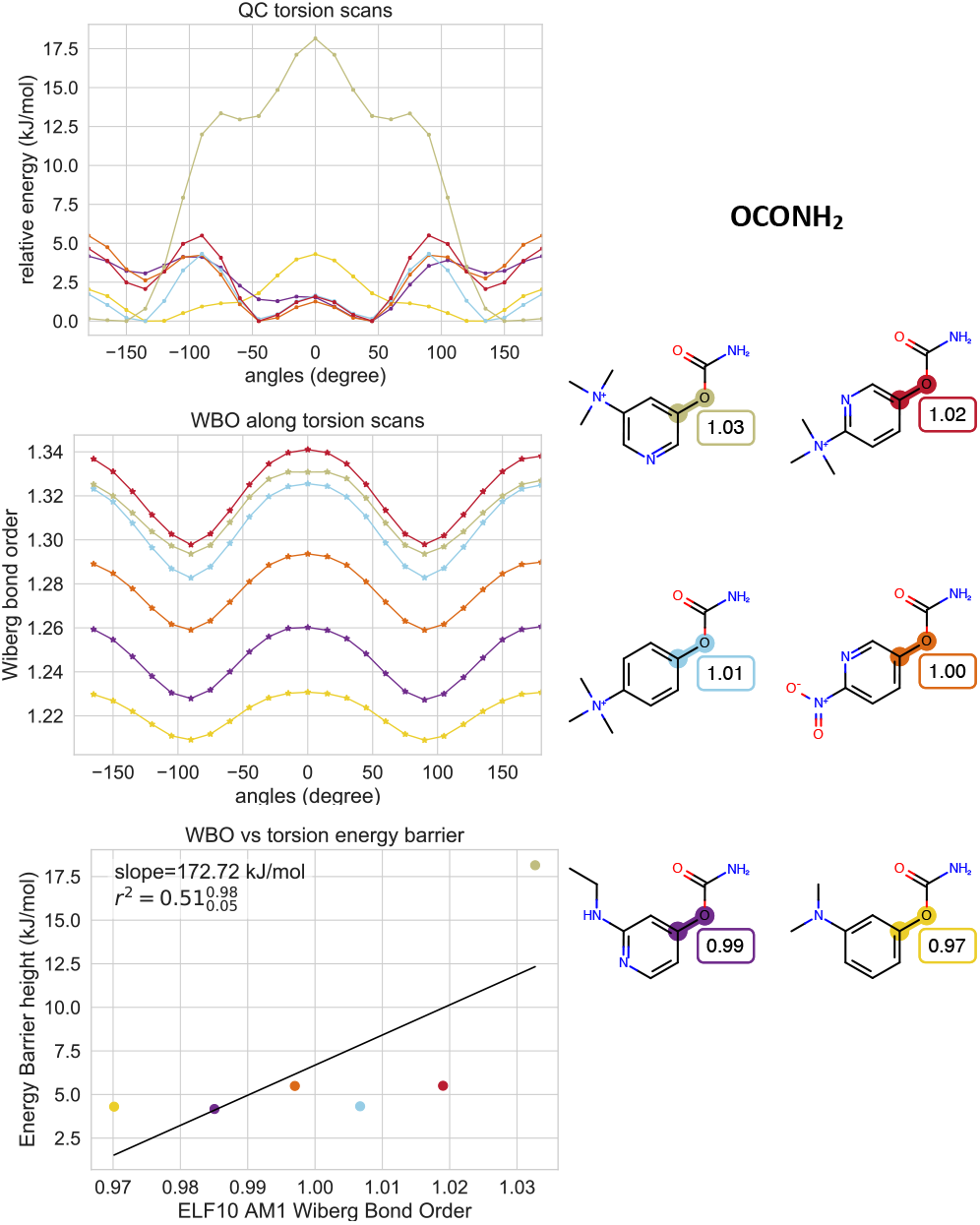
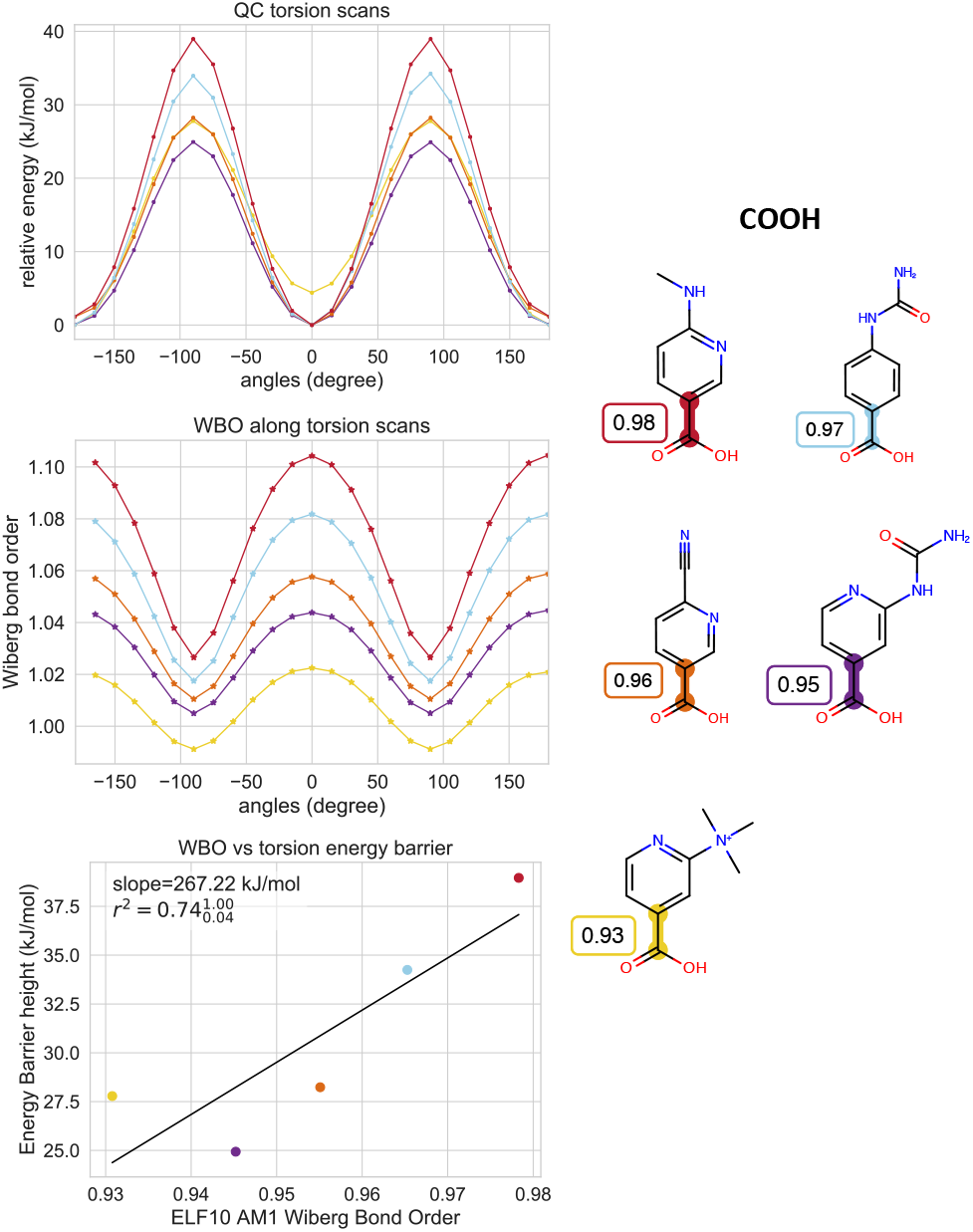
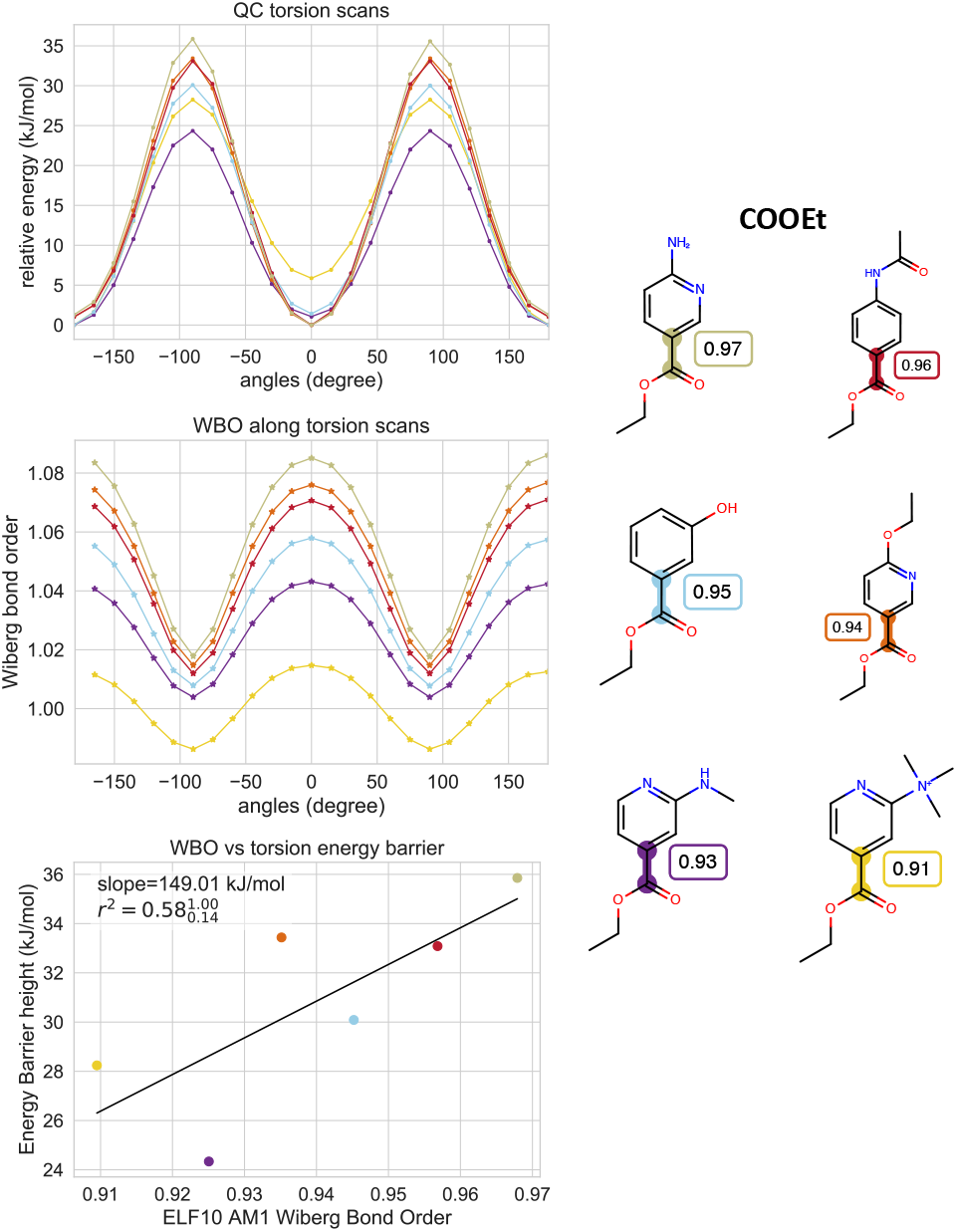
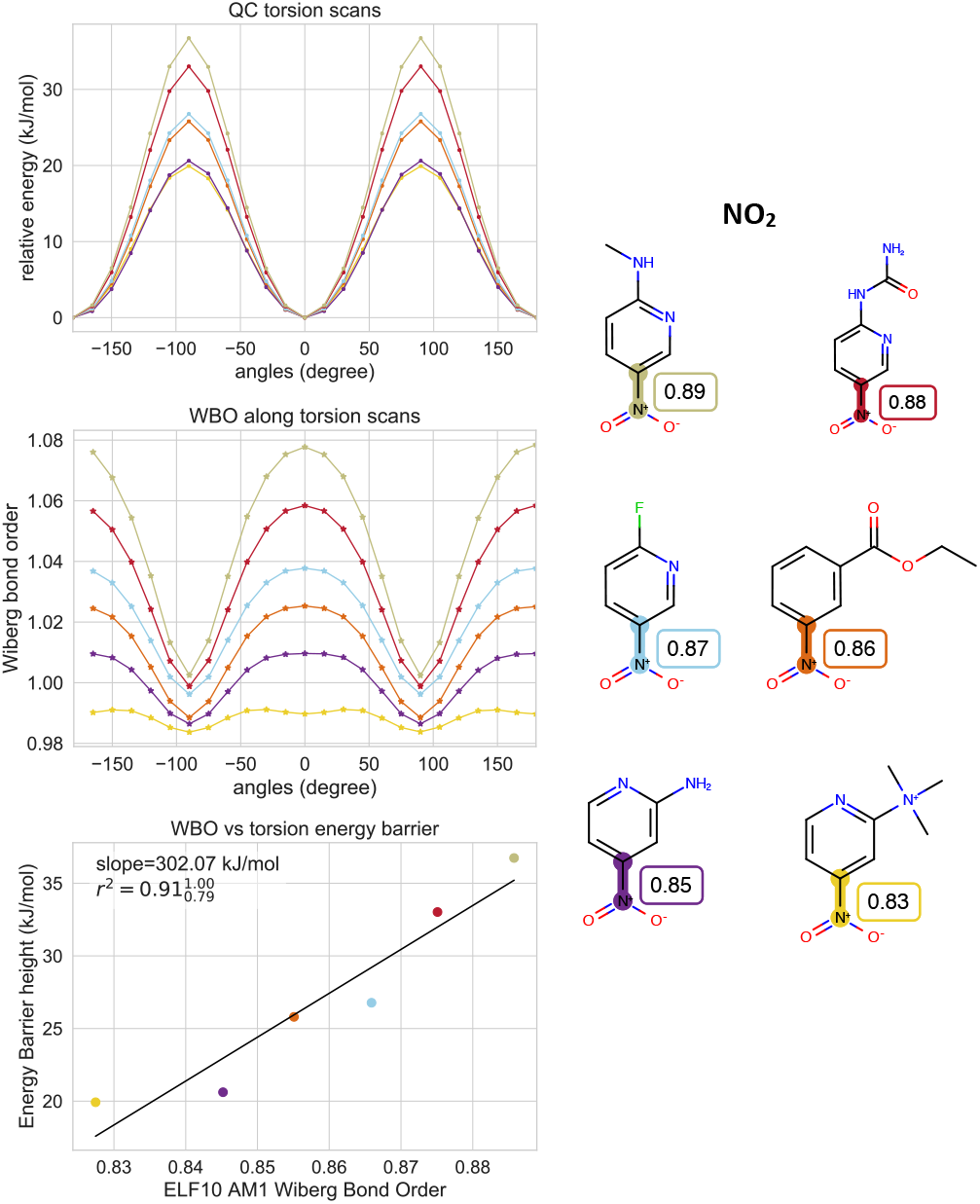
QC torsion scans and corresponding WBOs for substituted phenyl set. QC torsion scans, WBOs corresponding to scan and torsion barrier heights vs ELF10 WBOs for phenyl set. QC scan colors correspond to highlighted central bonds shown on the right. The molecules are labeled with their ELF10 WBOs.

**Appendix A Figure 7.**
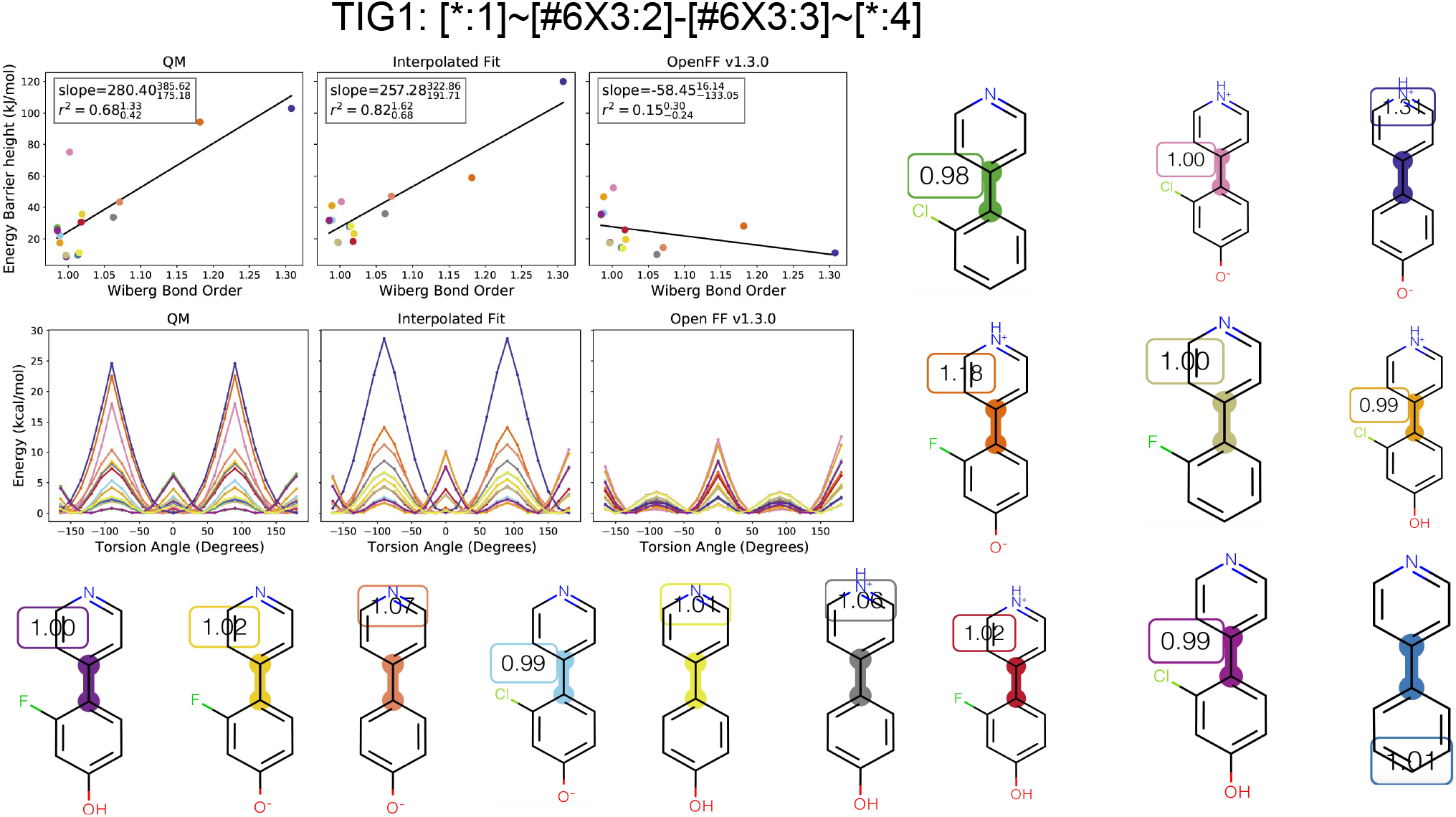
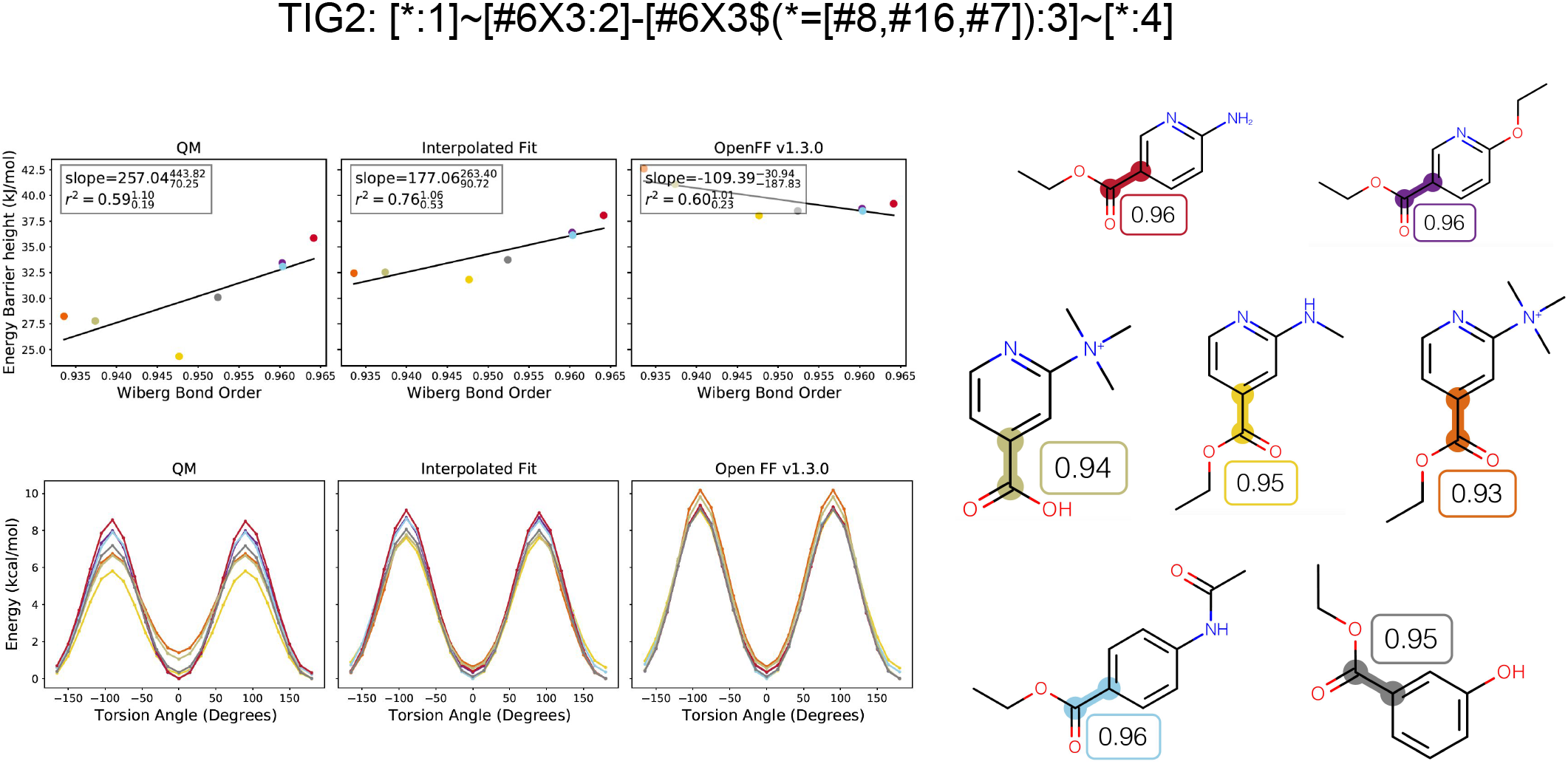
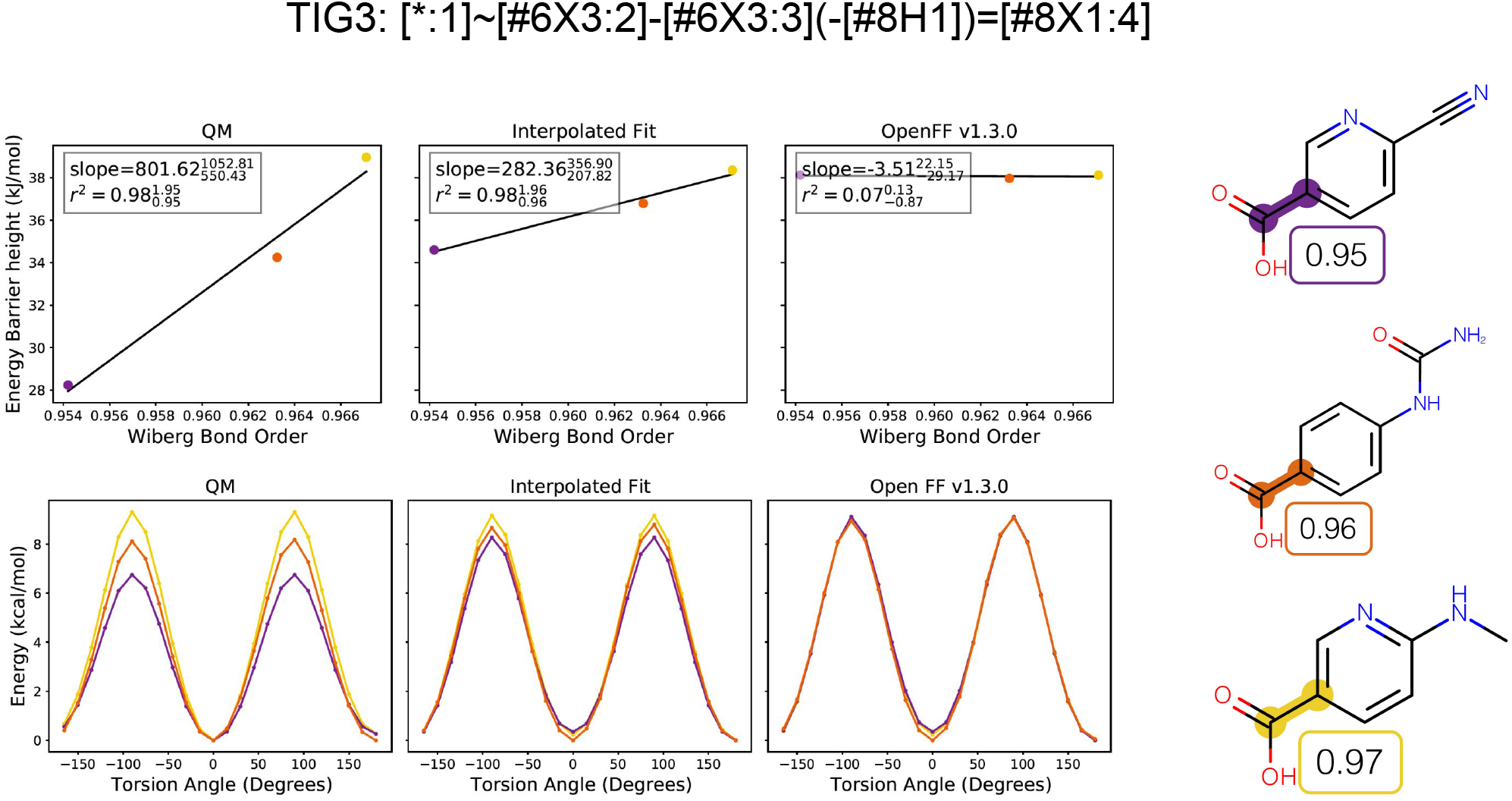
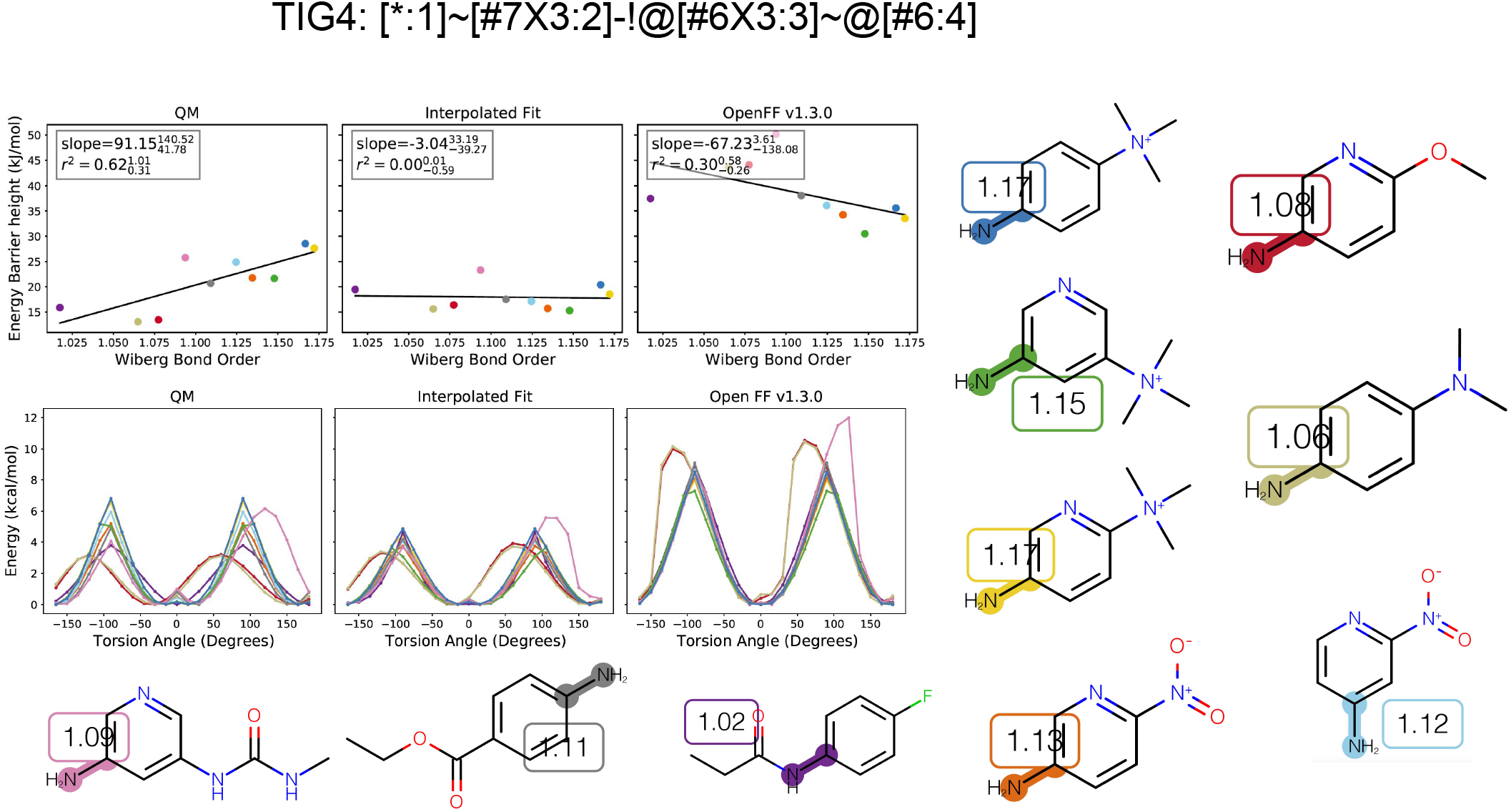
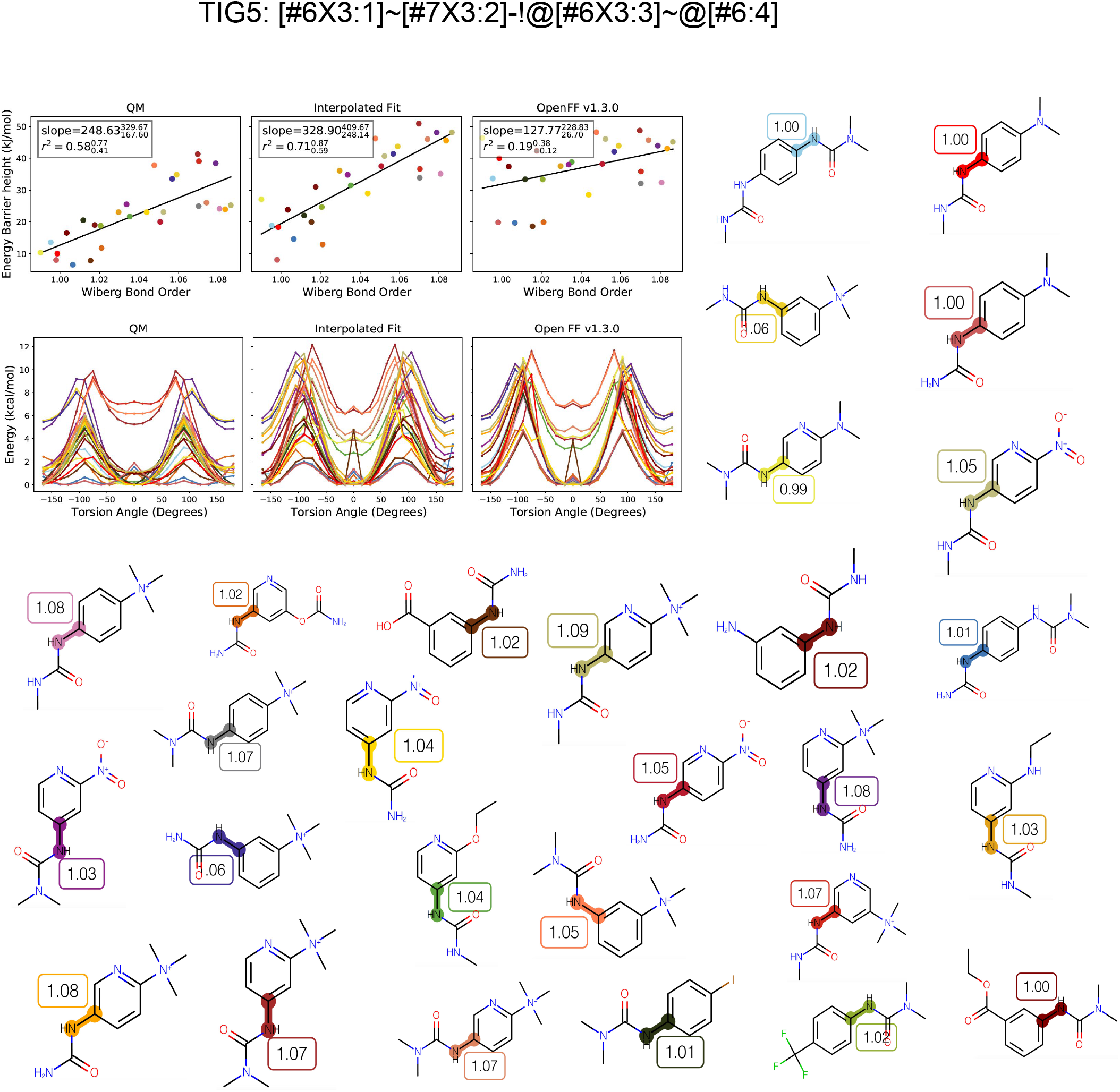
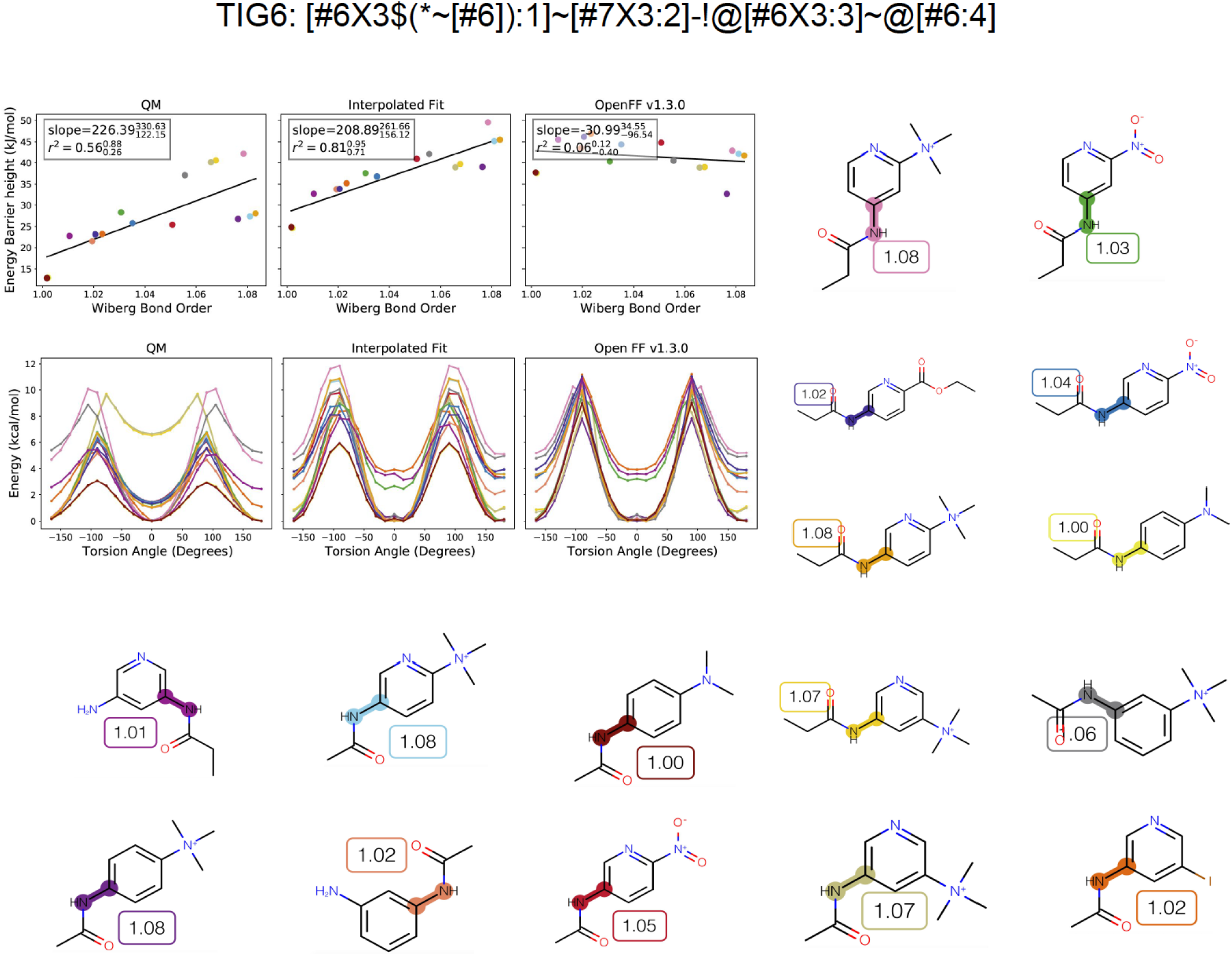
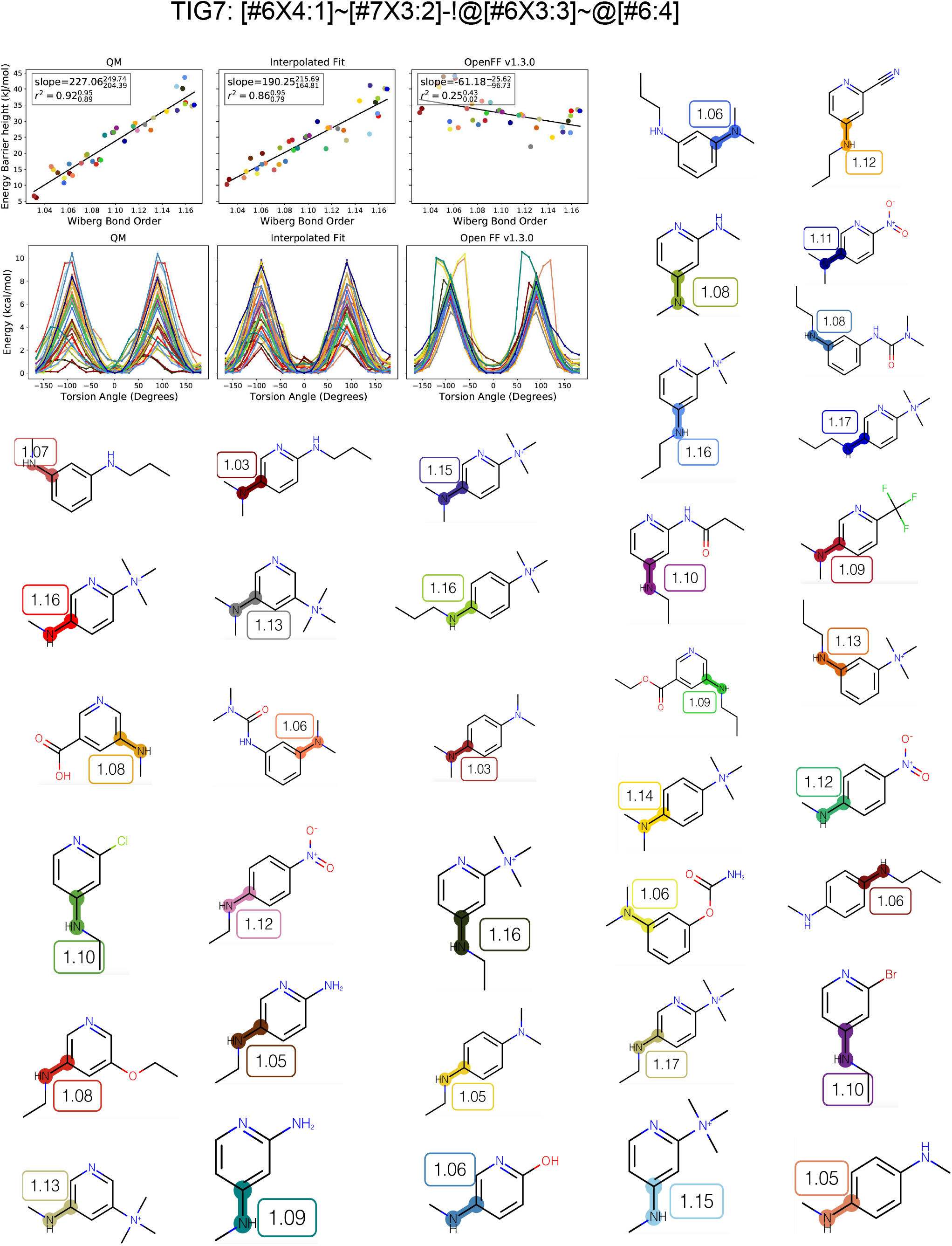
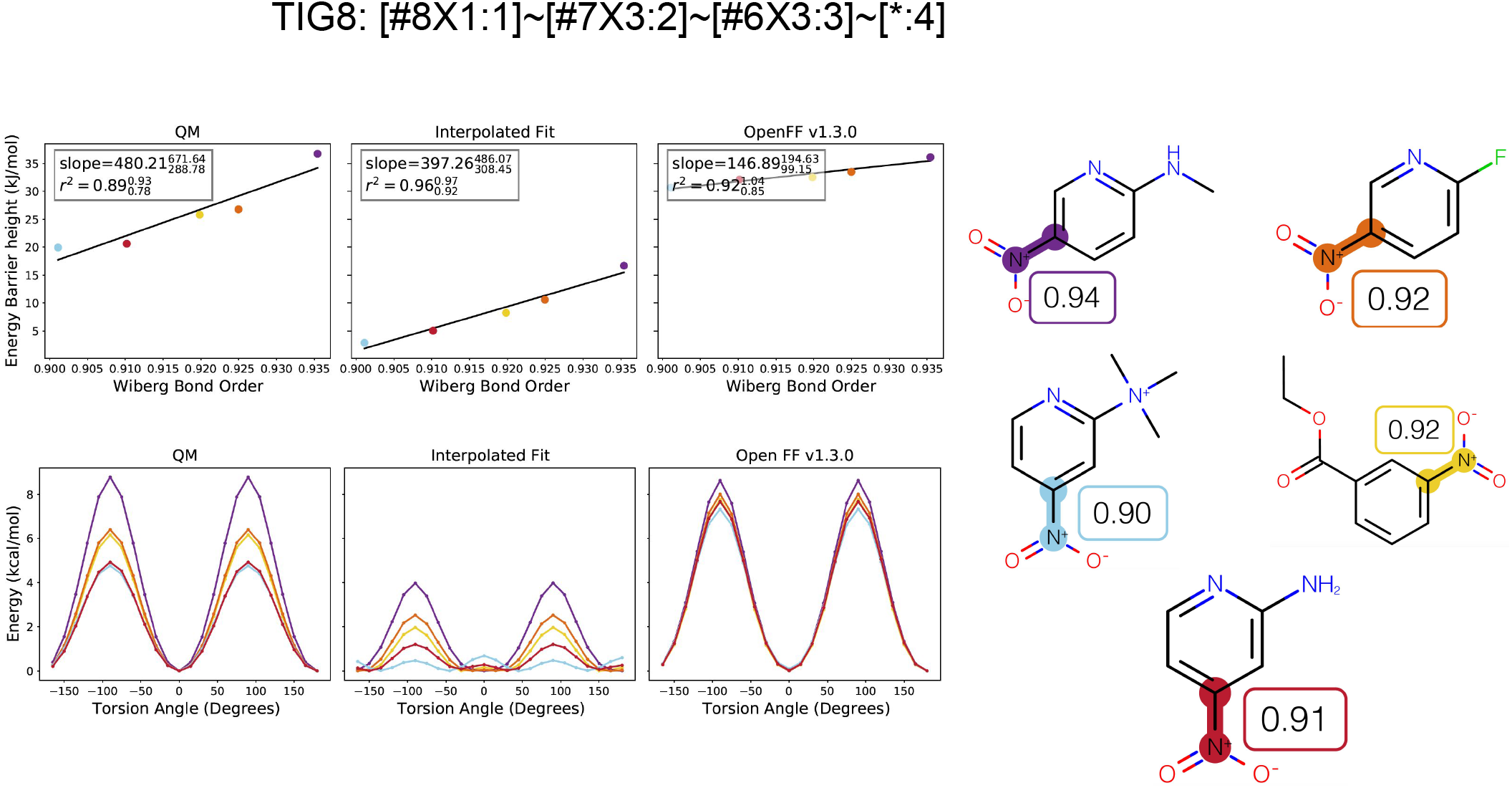
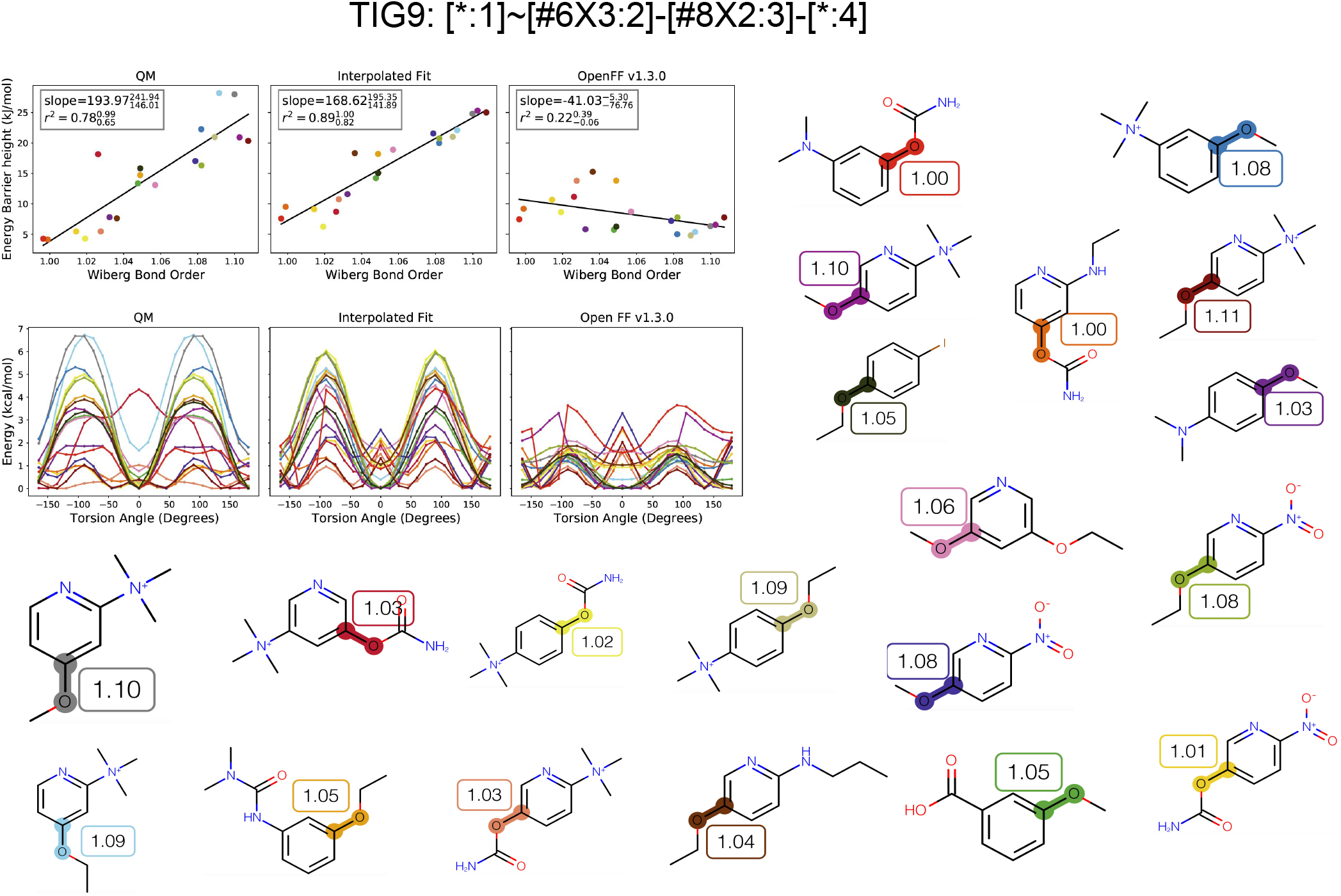
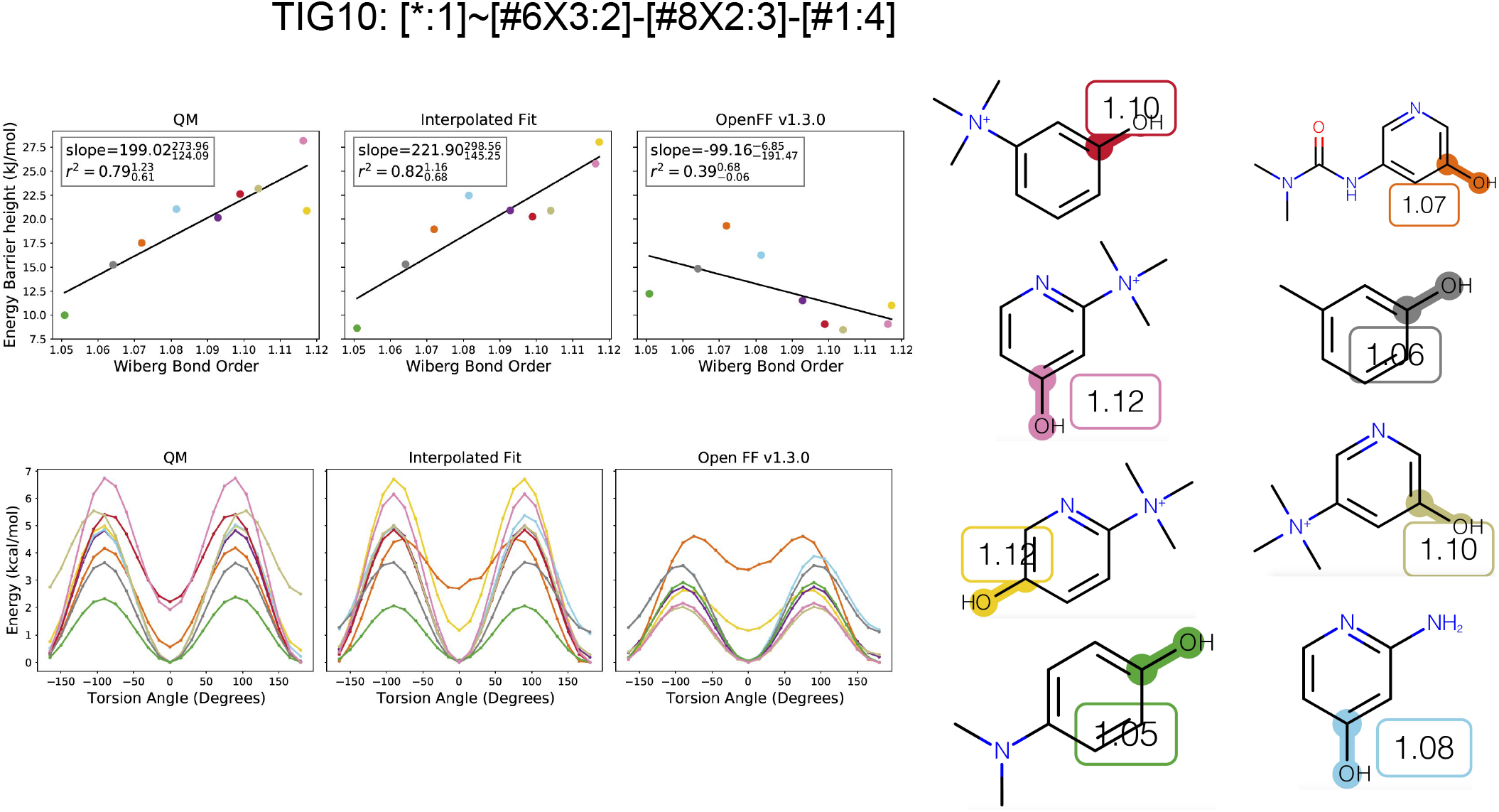
Comparing QM and MM torsion scans and torsion barrier heights for the FF with interpolation and OpenFF v1.3.0. The torsion scans correspond with the torsion barrier heights versus WBO. QC scan colors correspond to highlighted central bonds shown on the right. The molecules are labeled with their ELF10 WBOs.

## Footnotes

1 For non-minimal basis sets, AOs are often non-orthogonal and require normalization for the WBO to be valid. In the case of WBOs calculated by Psi4 [54] (used here), the Löwdin normalization [41, 51] scheme is used.

2 This is the canonical reference for the EFL10 method. The explanation provided in the paragraphs below is the most detailed description of the method

